# The relationship between diet, plasma glucose, and cancer prevalence across vertebrates

**DOI:** 10.1101/2023.07.31.551378

**Authors:** Stefania E. Kapsetaki, Anthony J. Basile, Zachary T. Compton, Shawn M. Rupp, Elizabeth G. Duke, Amy M. Boddy, Tara M. Harrison, Karen L. Sweazea, Carlo C. Maley

## Abstract

Could diet and mean plasma glucose concentration (MPGluC) explain the variation in cancer prevalence across species? We collected diet, MPGluC, and neoplasia data for 160 vertebrate species from existing databases. We found that MPGluC negatively correlates with cancer and neoplasia prevalence, mostly of gastrointestinal organs. Trophic level positively correlates with cancer and neoplasia prevalence even after controlling for species MPGluC. Most species with high MPGluC (50/78 species = 64.1%) were birds. Most species in high trophic levels (42/53 species = 79.2%) were reptiles and mammals. Our results may be explained by the evolution of insulin resistance in birds which selected for loss or downregulation of genes related to insulin-mediated glucose import in cells. This led to higher MPGluC, intracellular caloric restriction, production of fewer reactive oxygen species and inflammatory cytokines, and longer telomeres contributing to longer longevity and lower neoplasia prevalence in extant birds relative to other vertebrates.

## Introduction

Explaining patterns of cancer susceptibility among multicellular organisms is a major challenge in comparative oncology. Several life-history and environmental factors have been proposed to explain variations in cancer prevalence. Body size and longevity have been hypothesized to be predictors of cancer prevalence, based on the idea that larger animals (i.e. those with more cells), and long-lived animals (i.e. those with a longer time for mutations to accumulate^1^), should accumulate more carcinogenic mutations and thus have a higher cancer prevalence^2,3^. This has been shown in a study of free-living birds where body mass is positively correlated with tumor prevalence^4^. The majority of studies testing this hypothesis, however, show that larger and longer-lived animals do not get more cancer^5–10^. Other life history traits, such as shorter gestation length (across vertebrates^10^), and large litter size (in mammals^6^) or clutch size (in birds^8^), have been correlated with cancer prevalence. The reason for this positive correlation may be that animals may trade off resources between body maintenance (such as protection from DNA damage) and reproduction^11,12^. Other variables such as low habitat productivity (i.e., grams of glucose produced in the habitat of species per square meter per year), and low inferred metabolic rates are also correlated with cancer or neoplasia prevalence across vertebrates, but the correlations were not significant after False Discovery Rate corrections for performing multiple tests^13^. There is extensive evidence that diet affects cancer risk in humans^14–16^. Recently, researchers have begun investigating the role of diet in species’ susceptibility to cancer. Within mammals^17–19^ and across vertebrates^13^, higher trophic levels have higher cancer and neoplasia prevalence.

Glucose is a unifying factor among a number of risk factors for cancer prevalence across species, including larger litter/clutch size and carnivorous diet. The relationship between diet and blood glucose level is not simple. There is conflicting evidence about an association between diet and blood glucose levels in the literature. In humans^20^, common voles, tundra voles^21,22^, and vampire bats^23^, dietary components can change blood glucose levels. Birds, such as Noisy miners, that eat nectar and fleshy fruits have higher plasma glucose levels than birds, such as Welcome swallows, that eat insects^24,25^. Fish, such as sea bass, brown trout, and dogfish have lower blood glucose levels after fasting for a few days^26^. Yet mammals like cats and rats^27^, the “great fruit-eating bat” (*Artibeus lituratus*), and the “Jamaican fruit bat”^28^, maintain their blood glucose levels during fasting. Other studies in birds show that regardless of diet and the frequency of eating, blood glucose concentrations remain fairly constant^29^, with no significant differences in blood glucose concentrations between herbivorous, omnivorous, or insectivorous songbirds^30^. Also, many other fish species have stable blood glucose concentrations when starving^26^.

Diet appears to correlate with cancer prevalence across vertebrates^13,19^, but does not appear to correlate with glucose levels in the above studies of a few species in the literature. So why would we expect glucose levels to correlate with cancer prevalence across vertebrates? Blood glucose may be correlated with cancer prevalence across vertebrates because of: (1) the established positive relationship between trophic level (and their receptive macronutrient diets) and cancer prevalence^13,17–19^; (2) higher blood glucose being associated with increased oxidative stress, DNA damage, glycated proteins, and inflammation, all of which could increase cancer risk and development^31–36;^ and (3) the Warburg effect, in which cancers switch to a glucose-based metabolism rather than on oxygen-based metabolism^37–39^. Thus, one would expect higher blood glucose concentrations to be associated with higher cancer prevalence. This Warburg expectation, however, is in contrast with the dual role of glucose starvation in inducing both tumor suppression and tumor progression via cell-in-cell phenomena (such as cell cannibalism and entosis)^40,41^. In cross-species comparisons, birds are the taxon with the highest plasma glucose levels^42–44;^ 150–300% higher than mammals of similar body size^42^. Birds are also among the taxa with the lowest rate of neoplasia at necropsy; specifically lower than mammals and reptiles^45^. Thus, an examination of whether the Warburg hypothesis holds across species is warranted.

Associations of variables within species do not always generalize across species. For example, smaller dogs and shorter humans tend to live longer than larger dogs^46–48^ and taller humans^49^, respectively. However, there is a clear positive correlation between body size and lifespan across species (e.g.,^50–52)^. Thus, it may be the case that within humans and within other species, blood glucose levels may be positively correlated with cancer risk, but across species, blood glucose levels might be negatively correlated with cancer prevalence.

We sought to test three hypotheses: (1) diet influences mean plasma glucose concentration (MPGluC) across species; (2) cancer prevalence is higher in species with lower mean plasma glucose concentration; and (3) cancer prevalence is altered by (a) trophic level (herbivore, invertivore, primary carnivore, and secondary carnivore), (b) percentage of food type in the diet (fruit, invertebrates, plant, seeds, vertebrate ectotherms, and vertebrate endotherms), (c) percentage of animal-based and plant-based foods in the diet, and (d) overall diet type (herbivore, omnivore, carnivore), with or without controlling for mean plasma glucose concentration. We expected mean plasma glucose concentration to be negatively correlated with cancer and neoplasia prevalence across vertebrates, and more negatively (steeper slope and smaller *P*-value) correlated with gastrointestinal cancer and neoplasia prevalence. This expectation is based on the fact that the gastrointestinal tract, or the liver in the case of birds^53^, is the largest organ system that animals have^54^ in contact with the external environment (e.g. food), the largest immune organ in the body^55^, and the role of the largest mucosal absorption surface area in the body in the breakdown and absorption of glucose^56^. Even though the associations between diet and cancer prevalence have been previously studied across vertebrates^13^, we reevaluate previous results in a subset of species for which we had plasma glucose data and examine the unknown association between diet and cancer when controlling for mean plasma glucose concentration across vertebrates. We test these associations using literature resources on diet, Species360 data on glucose concentrations in the plasma, and cancer prevalence data from 160 species that were necropsied under veterinary care.

## Results

Mean plasma glucose concentration varied considerably across the examined vertebrate species (Supplementary Material). The burmese python (*Python bivittatus*) had the lowest mean plasma glucose concentration (1.22 mmol/L), whereas the bird “blue-bellied roller” (*Coracias cyanogaster*) had the highest mean plasma glucose concentration (21.08 mmol/L). When comparing different classes, the mean of the mean plasma glucose concentrations in Amphibia (4 species) was 2.05 ± 0.42 (mean ± standard deviation) mmol/L, 4.56 ± 4.01 mmol/L in Reptilia (35 species), 6.67 ± 1.54 mmol/L in Mammalia (69 species), and 15.31 ± 3.12 mmol/L in Aves (51 species).

### Relationships Between Diet and Plasma Glucose

There is no association between diet and mean plasma glucose concentration across vertebrate species included in this study after correcting for the false discovery rate of multiple testing (Fig. 1). Trophic levels do not explain the diversity in the concentration of glucose in the plasma across 160 species (Fig. 1A). Similarly, the percentage of animal-based vs. plant-based foods (n = 44 and n = 50 species, respectively), or diet type (n = 67 species), is not significantly associated with differences in mean plasma glucose concentration (Fig. 1C, 1D).

**Figure 1.**
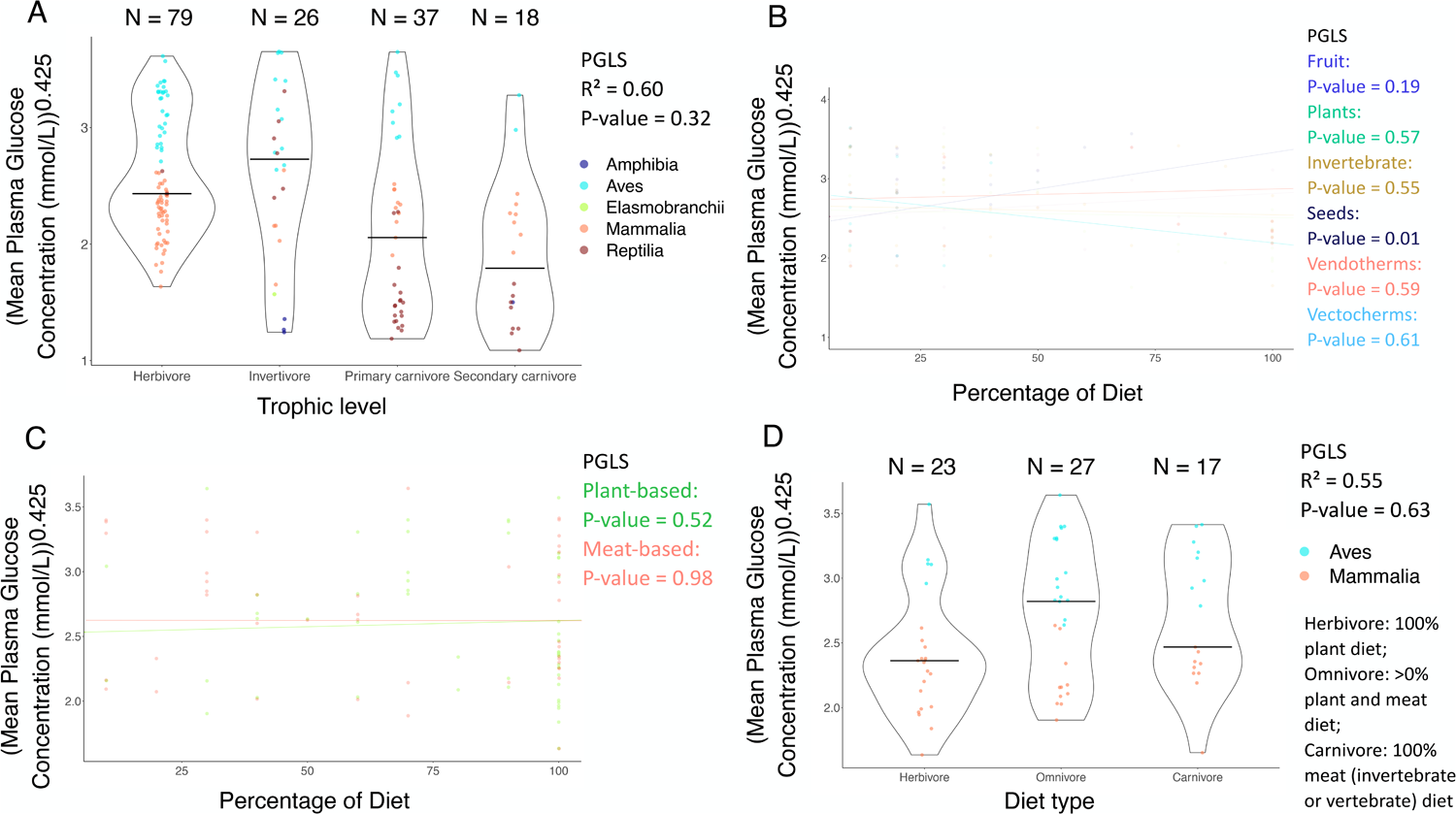
The relationship between mean plasma glucose concentration and diet across vertebrates. (**A**) Trophic level is not significantly correlated with mean plasma glucose concentrations across 160 species (PGLS: *P*-value = 0.32; R^2^ = 0.60). (**B**) The percentage of seeds in a species’ diet is positively correlated with mean plasma glucose concentrations for 18 species (PGLS: *P*-value = 0.01; R^2^ = 0.56), but not after correcting for multiple testing (Table 1A). There was no significant correlation between the percentage of fruit, plants, invertebrates, endothermic vertebrates (Vendotherms), or ectothermic vertebrates (Vectotherms), and mean plasma glucose concentrations for 25, 39, 35, 15, or 11 species, respectively (PGLS: *P*-value > 0.05). (**C**) The percentage of plant-based foods or meat-based foods in a species’ diet was not significantly correlated with mean plasma glucose concentrations for 50 or 44 species, respectively (PGLS: *P*-value > 0.05). (**D**) Diet type is not significantly correlated with mean plasma glucose levels for 67 species (PGLS: *P*-value = 0.63; R^2^ = 0.55). The horizontal black line in each diet category shows the mean plasma glucose concentration in that trophic (plot A) or diet category (plot D). Each dot shows the mean plasma glucose concentration and diet category of one species. N shows the number of species per diet category (plots A & D). We added minimal jitter in the plots in order to better visualize individual data points.

### Relationships Between Plasma Glucose and Cancer

Mean plasma glucose concentration is not correlated with cancer prevalence (including all tissues) (Fig. 2; Supp. Fig. 1A). However, mean plasma glucose concentration is negatively correlated with neoplasia prevalence (including all tissues) when controlling the analysis for body weight (Table 1B). Mean plasma glucose concentration is also negatively correlated with gastrointestinal neoplasia prevalence (Supp. Fig. 1B; Table 1B), even when controlling for birds vs. non-birds, and body weight (Table 1B). Mean plasma glucose concentration is negatively correlated with gastrointestinal cancer prevalence (measured from a total of 242 records of gastrointestinal malignancies across 108 species), even after controlling for weight, birds versus non-birds (Fig. 3; Table 1B). Mean plasma glucose concentration is negatively correlated with non-gastrointestinal neoplasia prevalence only when we control the analyses for weight (Table 1B). Within mammals (Supp. Fig. 5; Table 1C; n = 48–68 species), birds (Supp. Fig. 6; Table 1D; n = 31–51 species), and reptiles (Supp. Fig. 7; Table 1E; n = 25–35 species) there were no significant correlations between mean plasma glucose concentrations and cancer prevalence or neoplasia prevalence across tissues, gastrointestinal cancer prevalence or gastrointestinal neoplasia prevalence, and non-gastrointestinal cancer prevalence or non-gastrointestinal neoplasia prevalence after applying multiple testing corrections.

**Figure 2.**
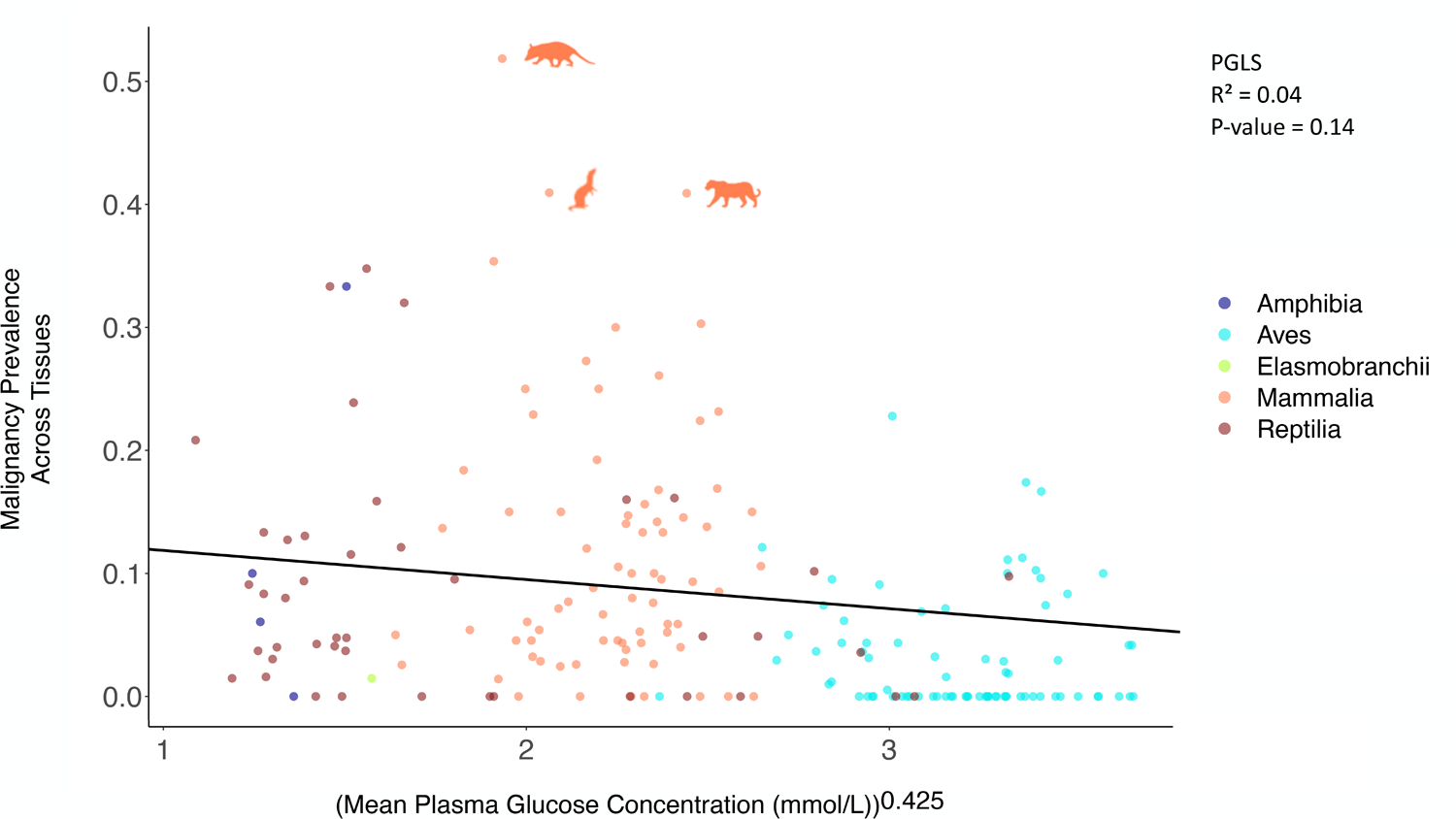
The relationship between mean plasma glucose concentration and overall cancer prevalence. This non-significant correlation (PGLS: *P*-value = 0.14) was across 160 species. Each dot represents the average plasma glucose concentration and the malignancy prevalence across tissues of one species; Amphibia: Dark Blue; Aves: Blue; Elasmobranchii: Green; Mammalia: Orange; Reptilia: Red. For ease of interpretation of significant outliers (Rosner’s test), we show images of outlier species. Animal silhouettes from PhyloPic (http://www.phylopic.org/).

**Table 1.**
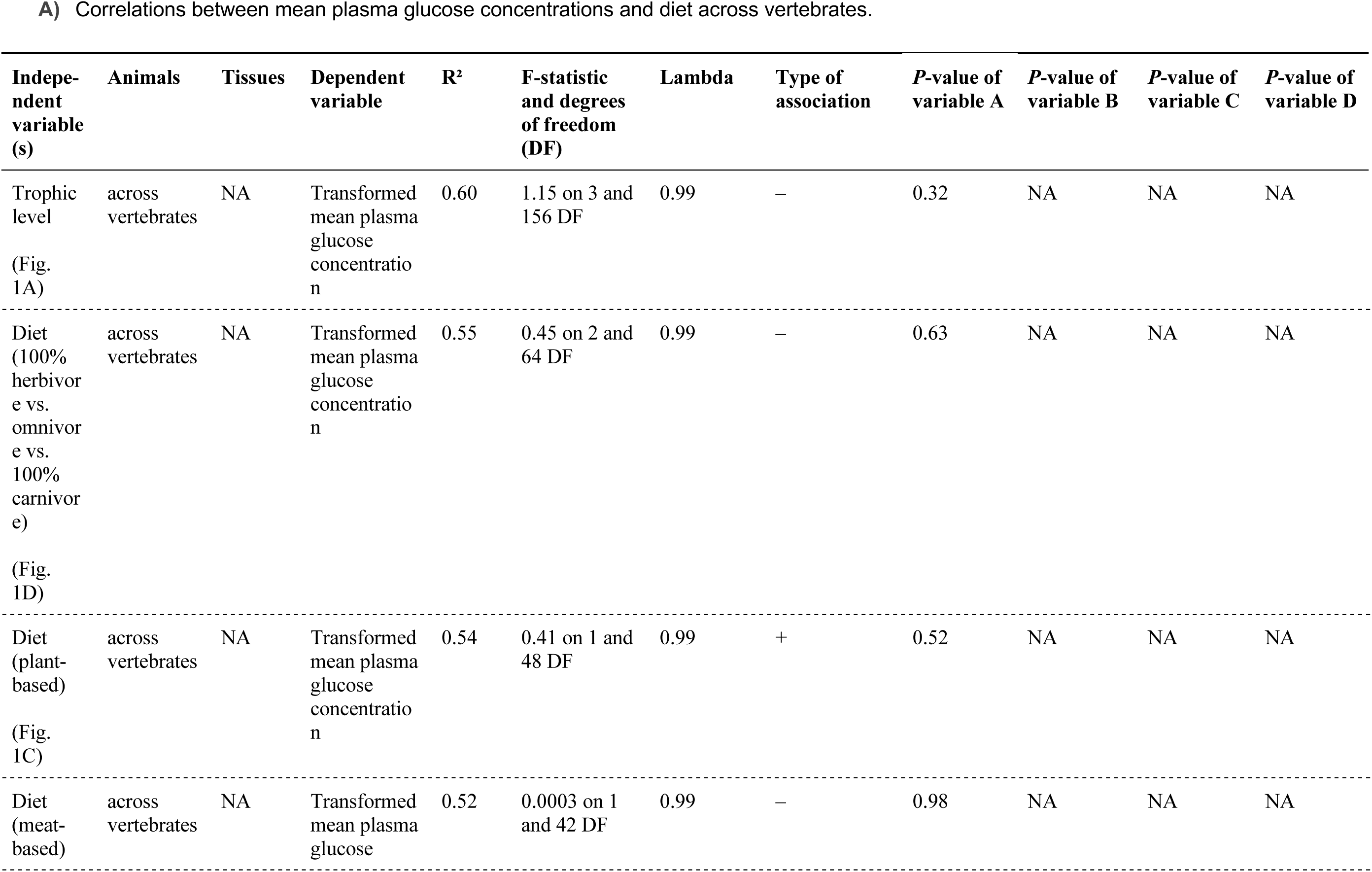
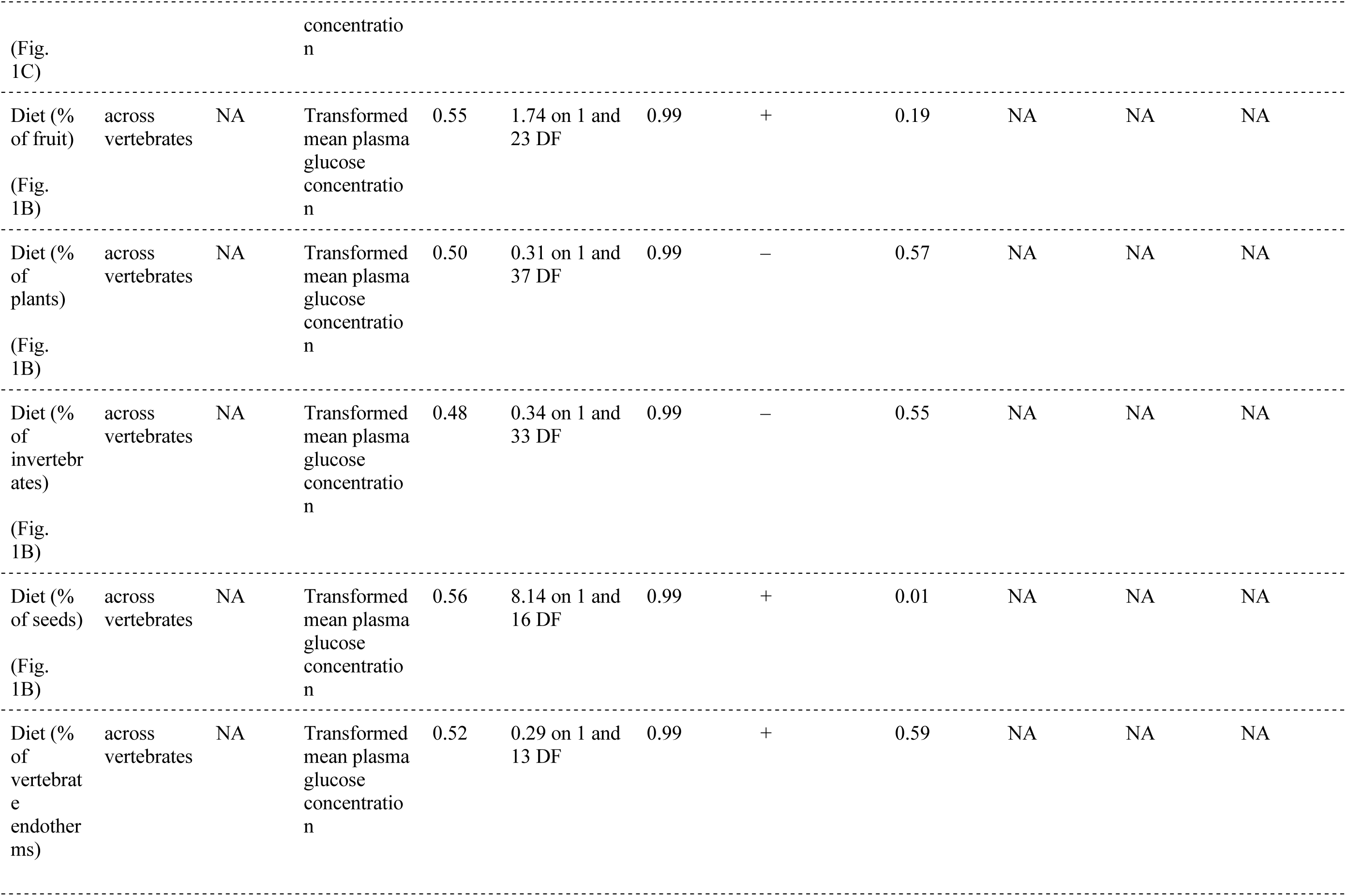
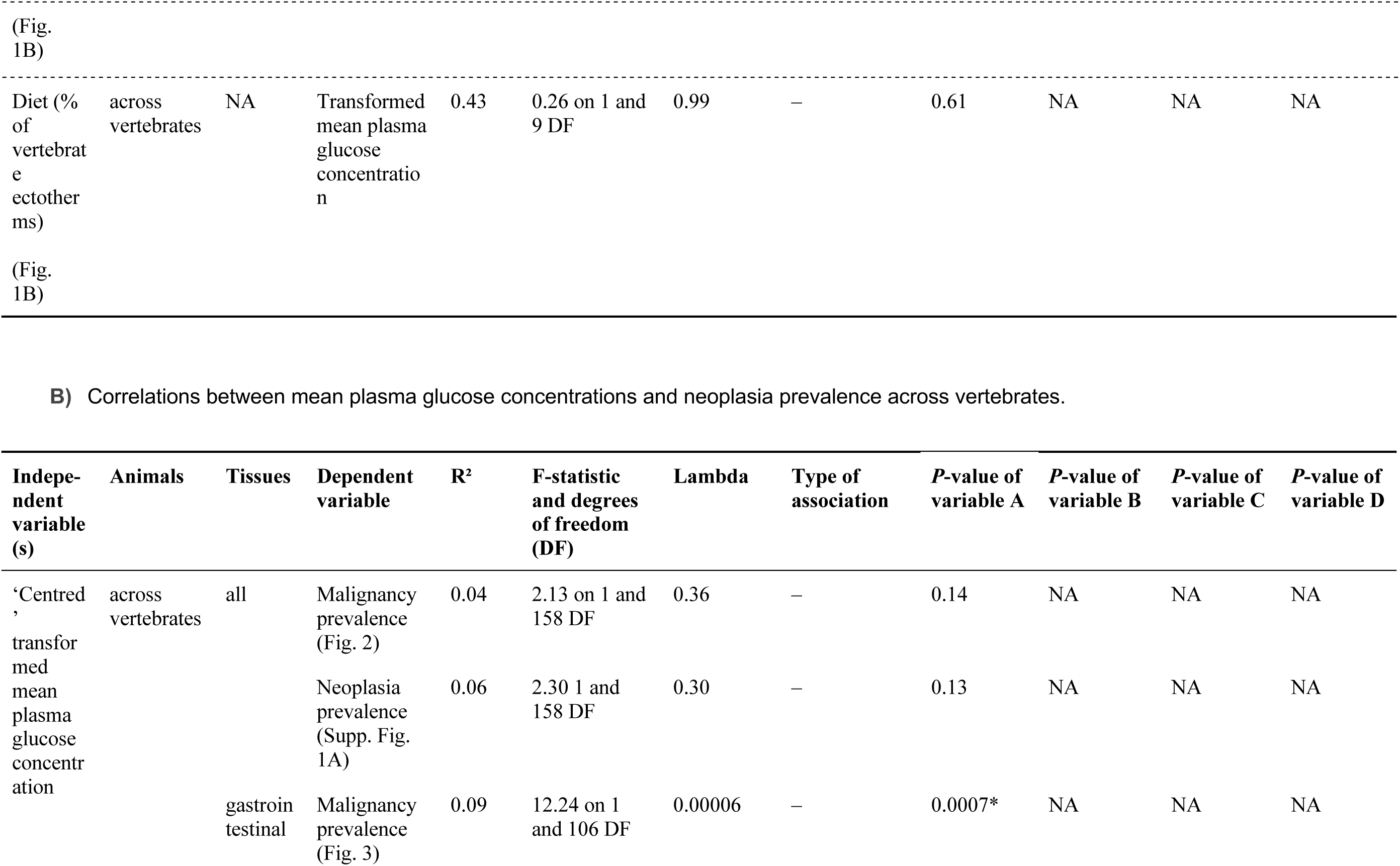
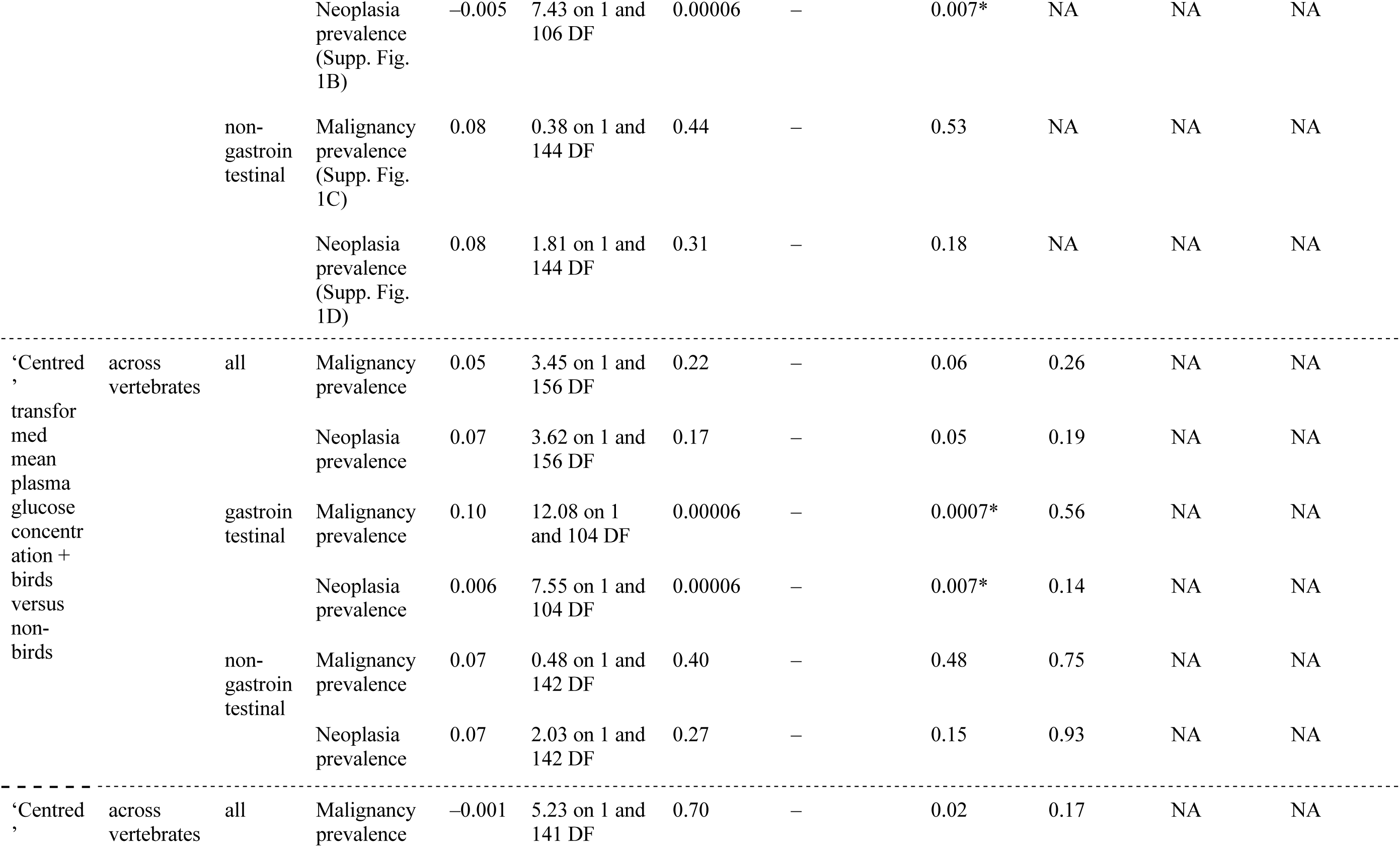
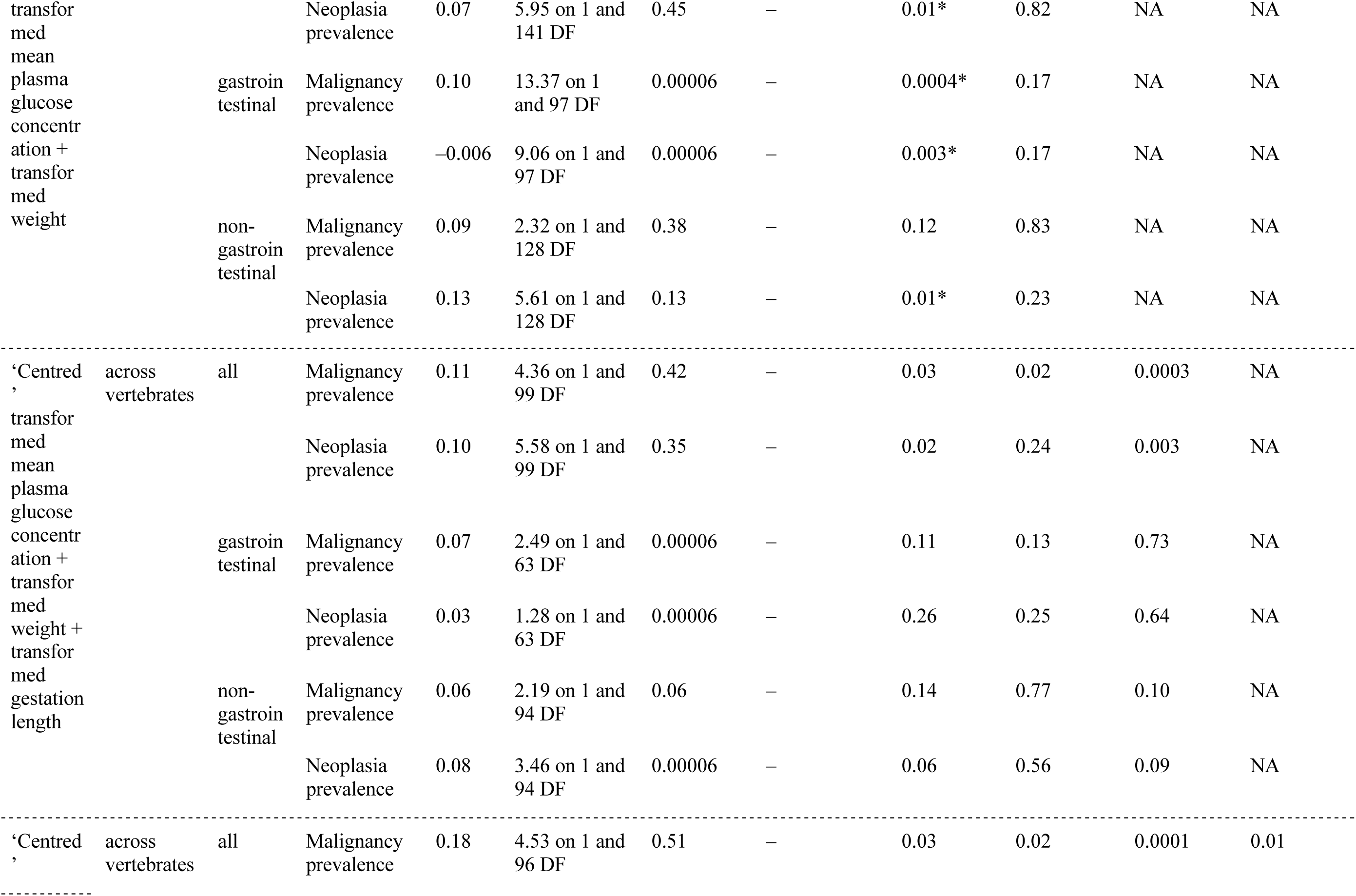
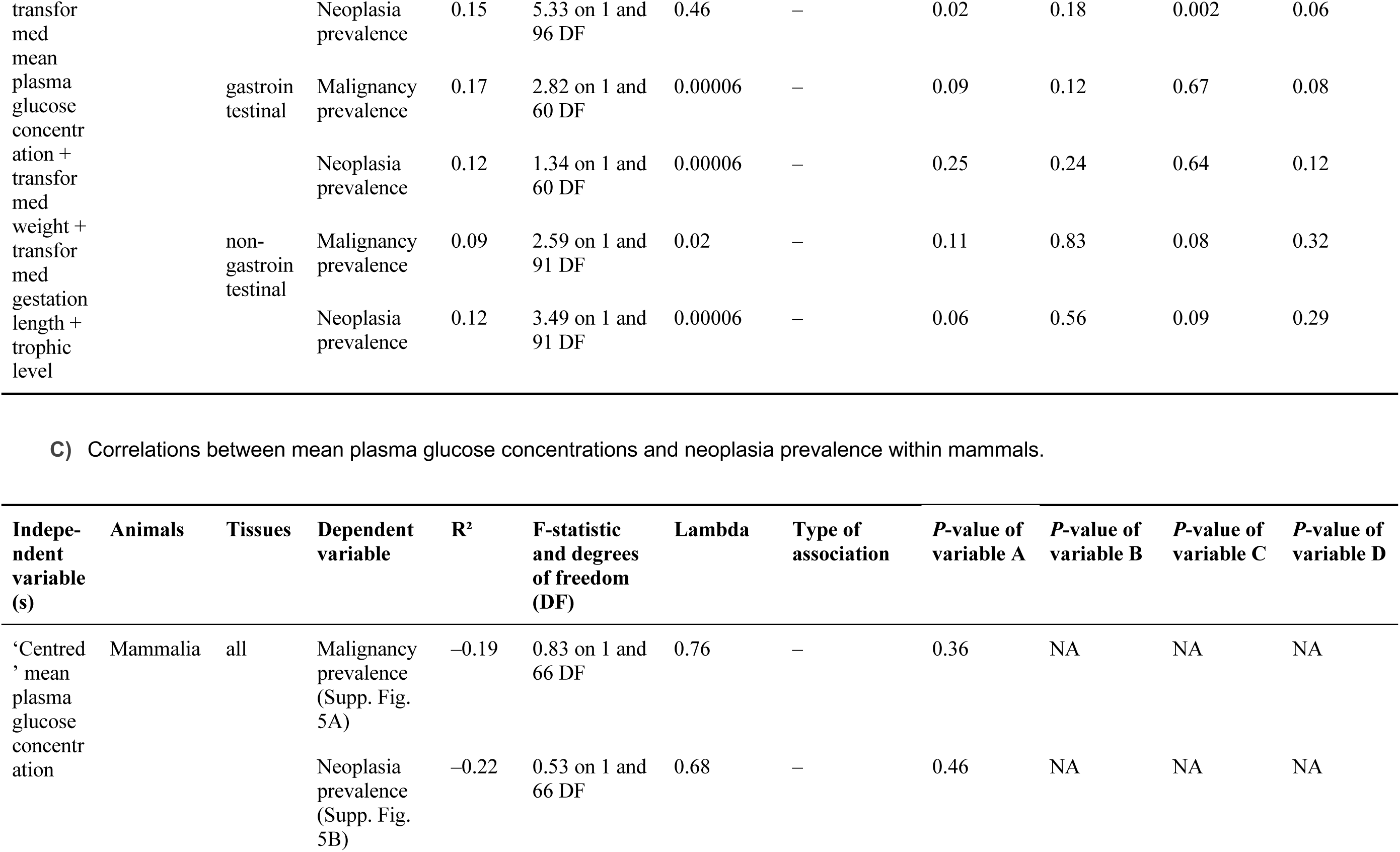
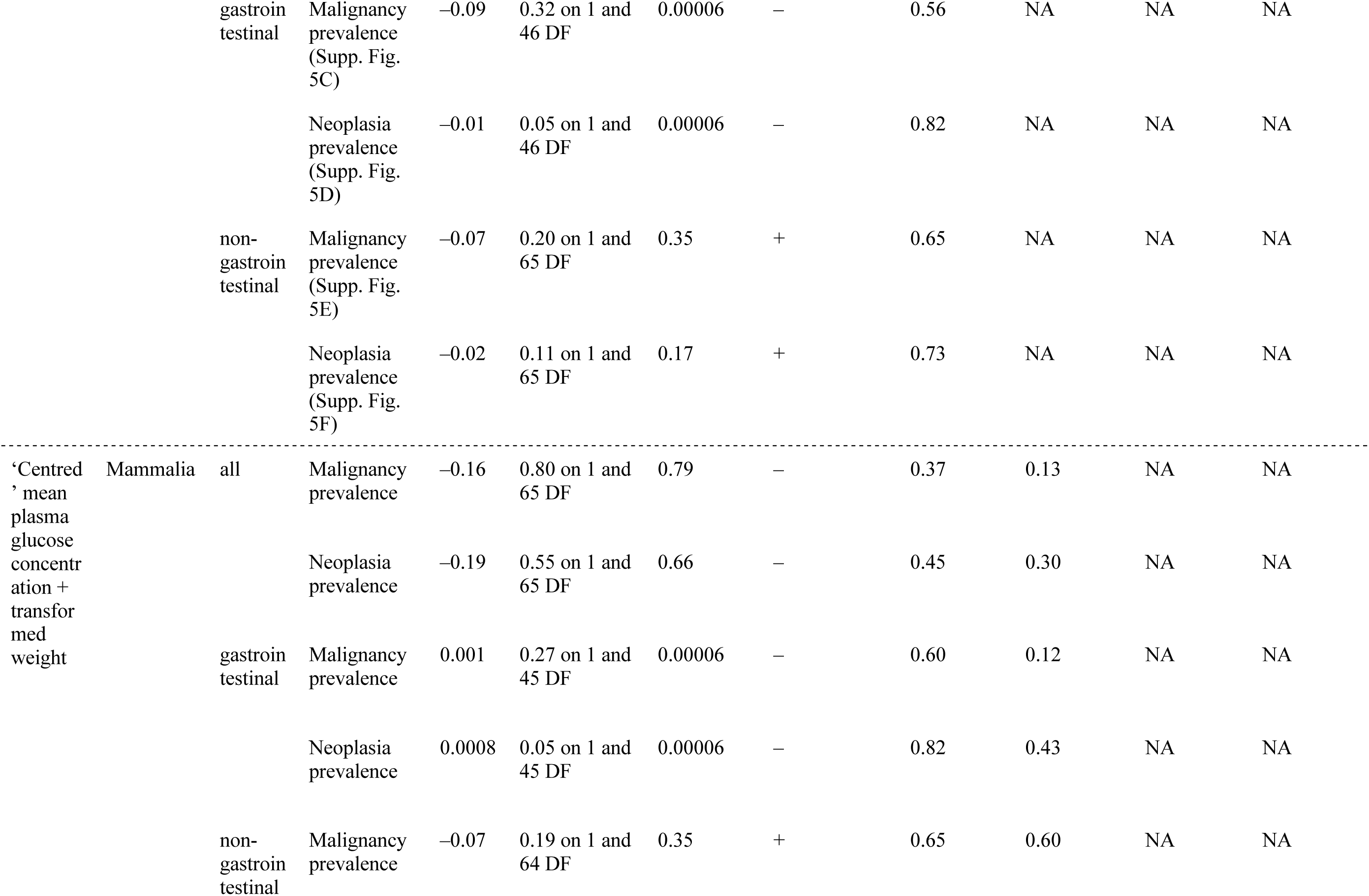
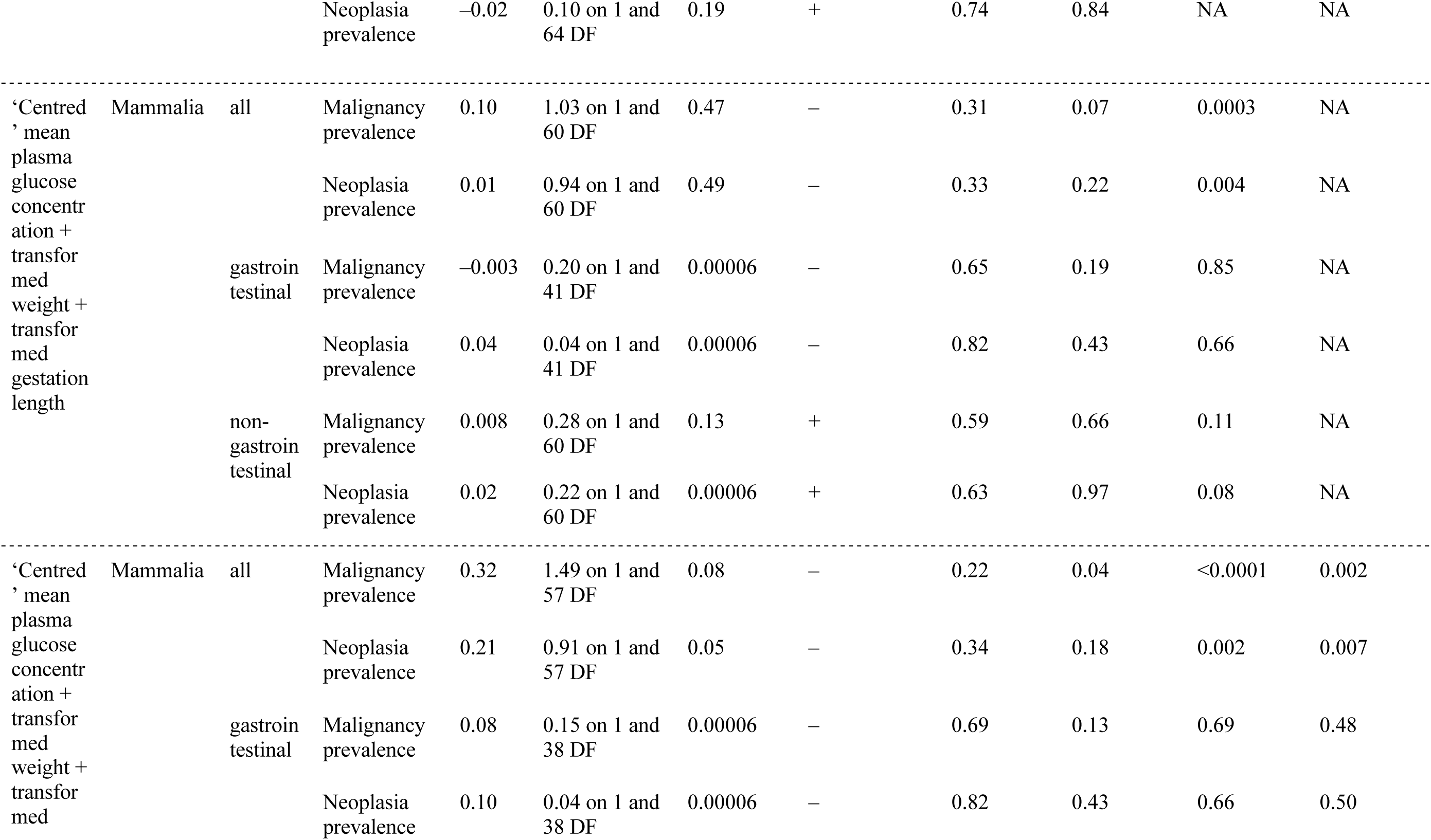
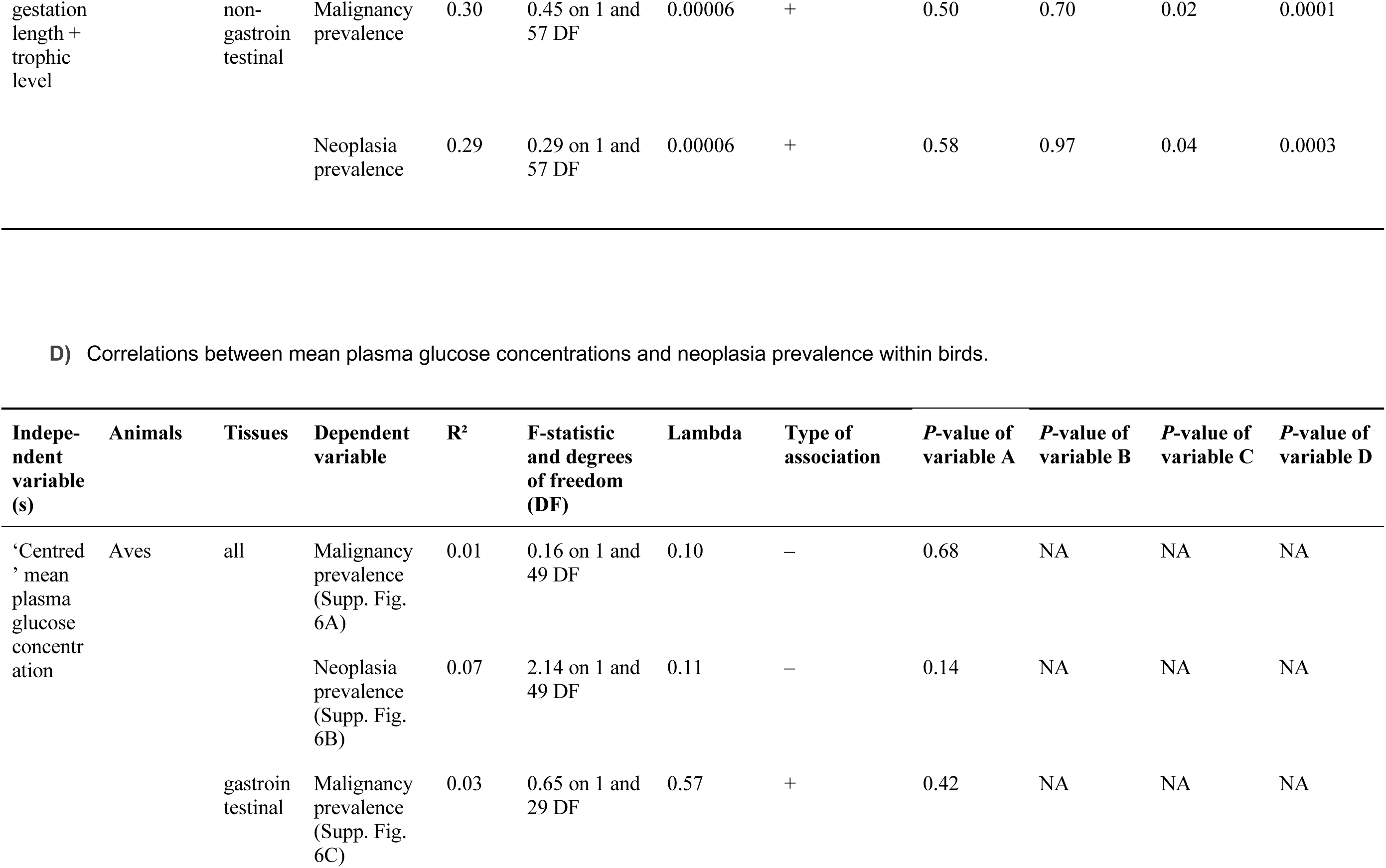
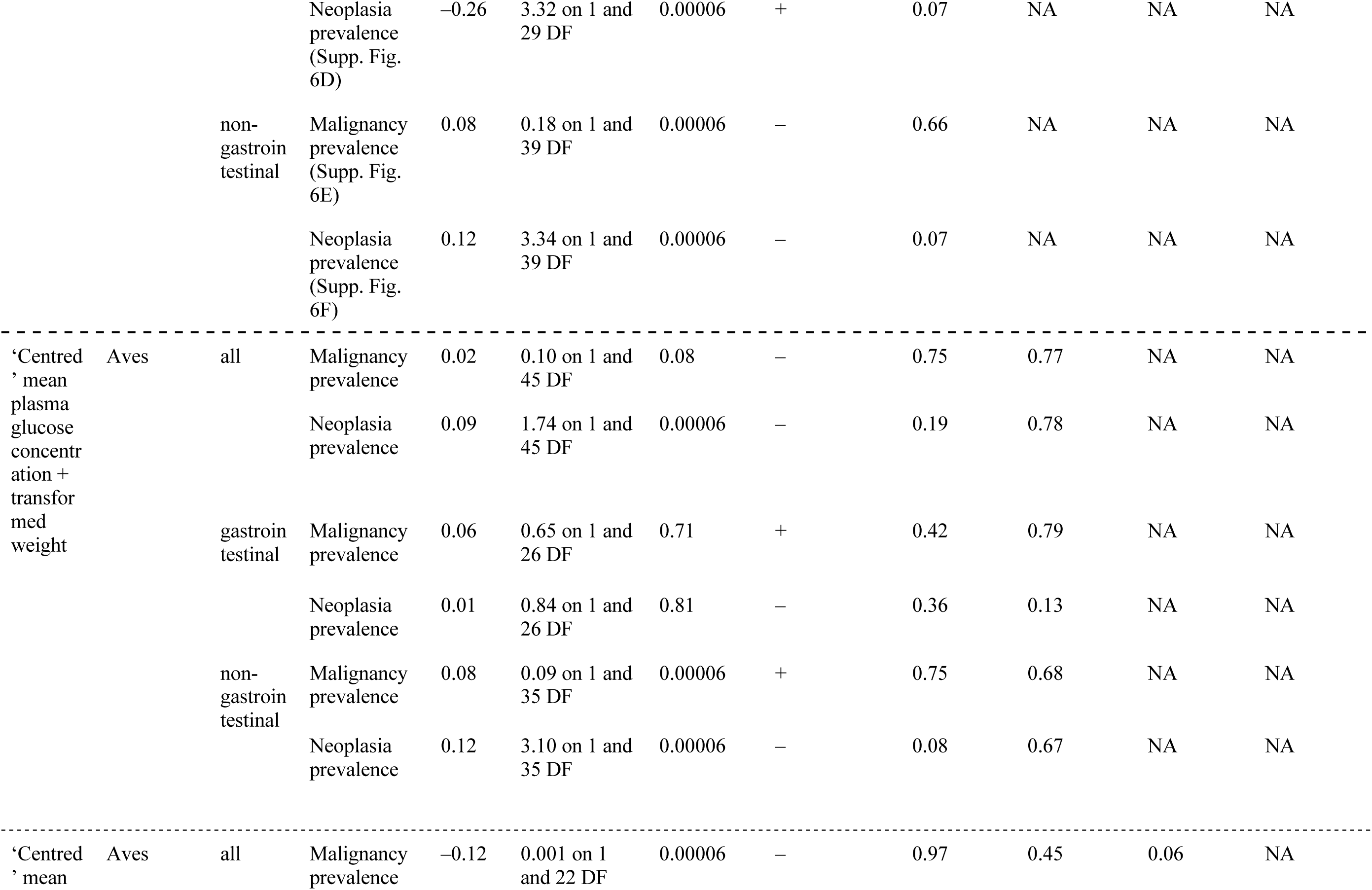
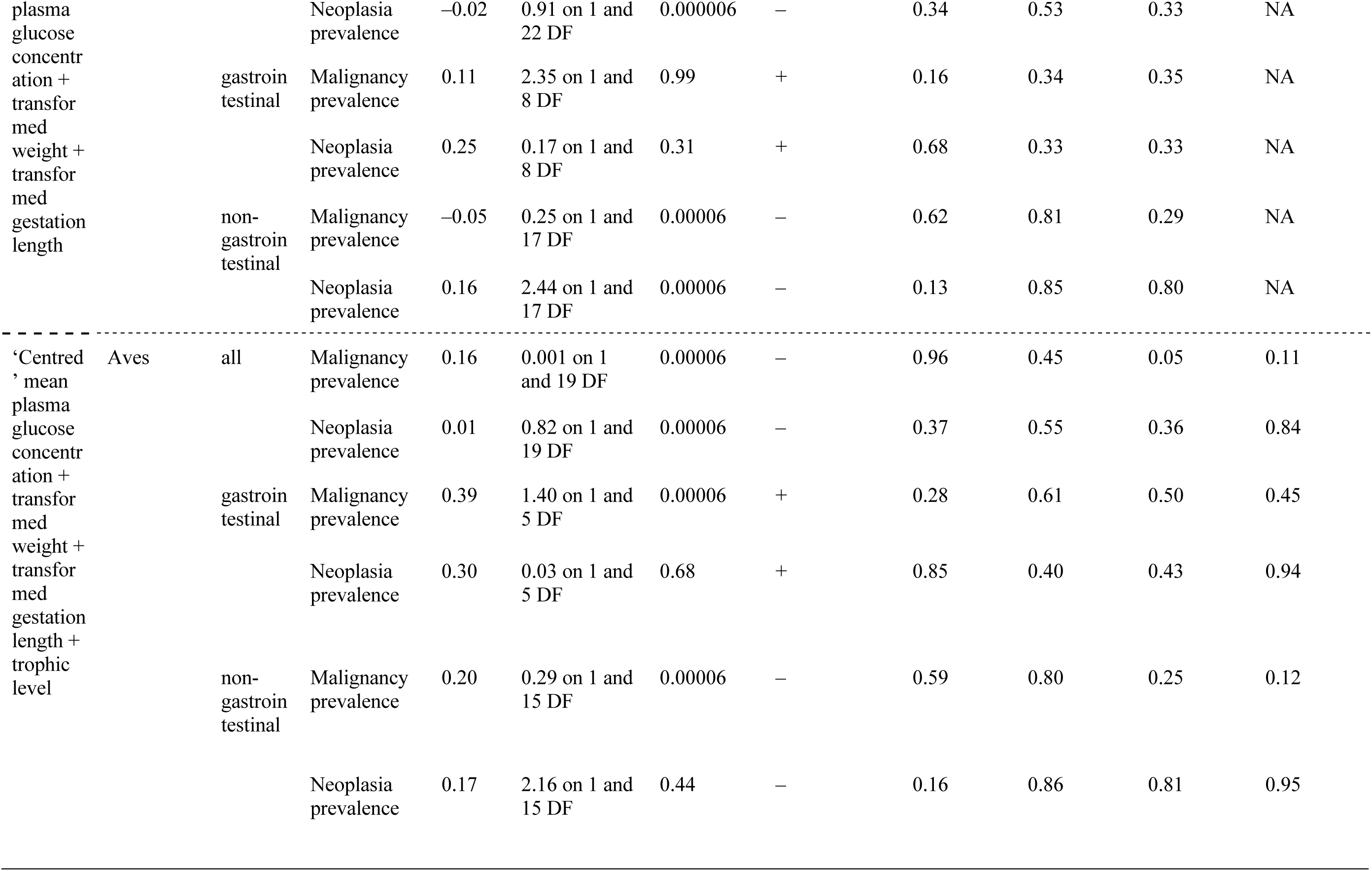
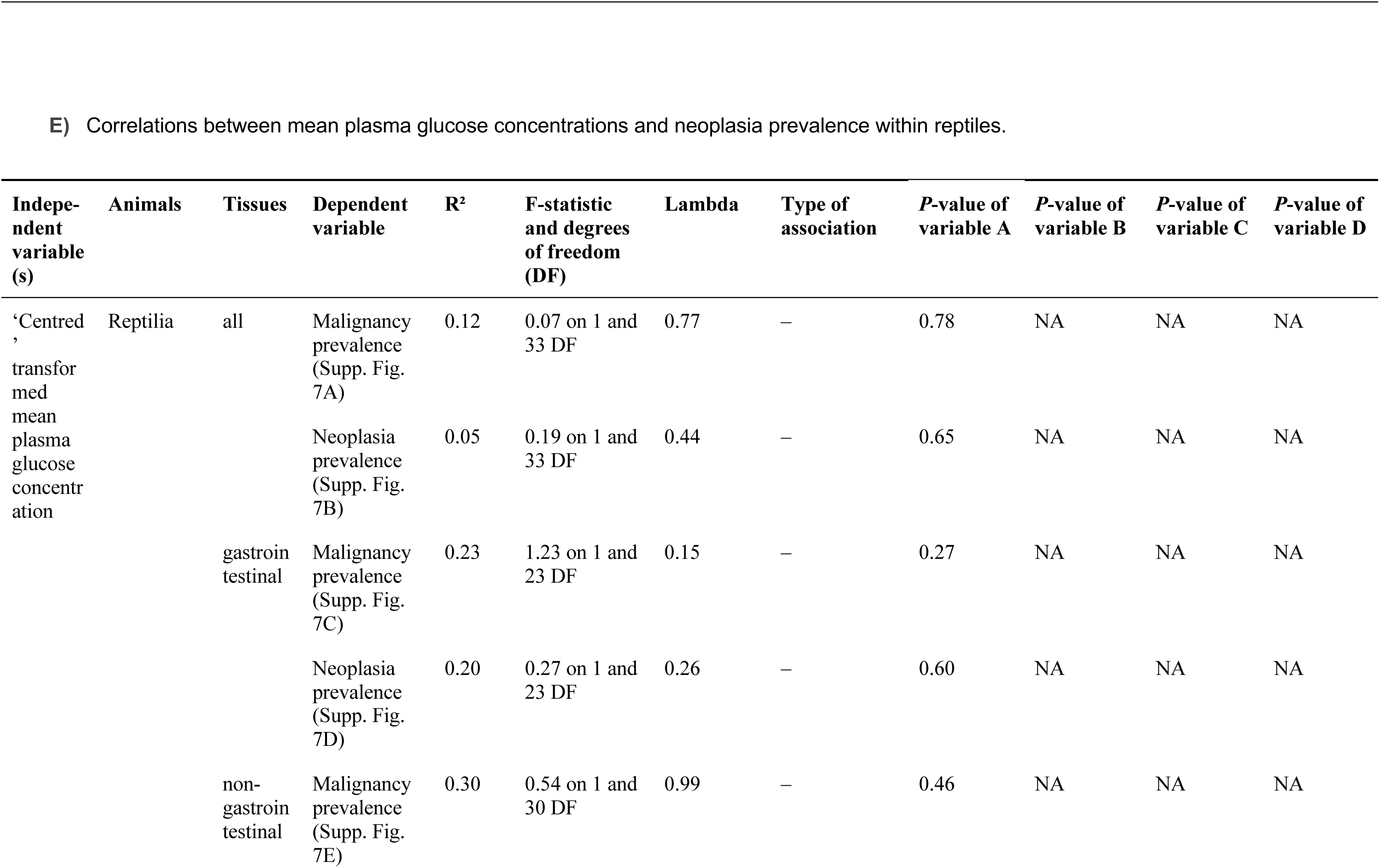
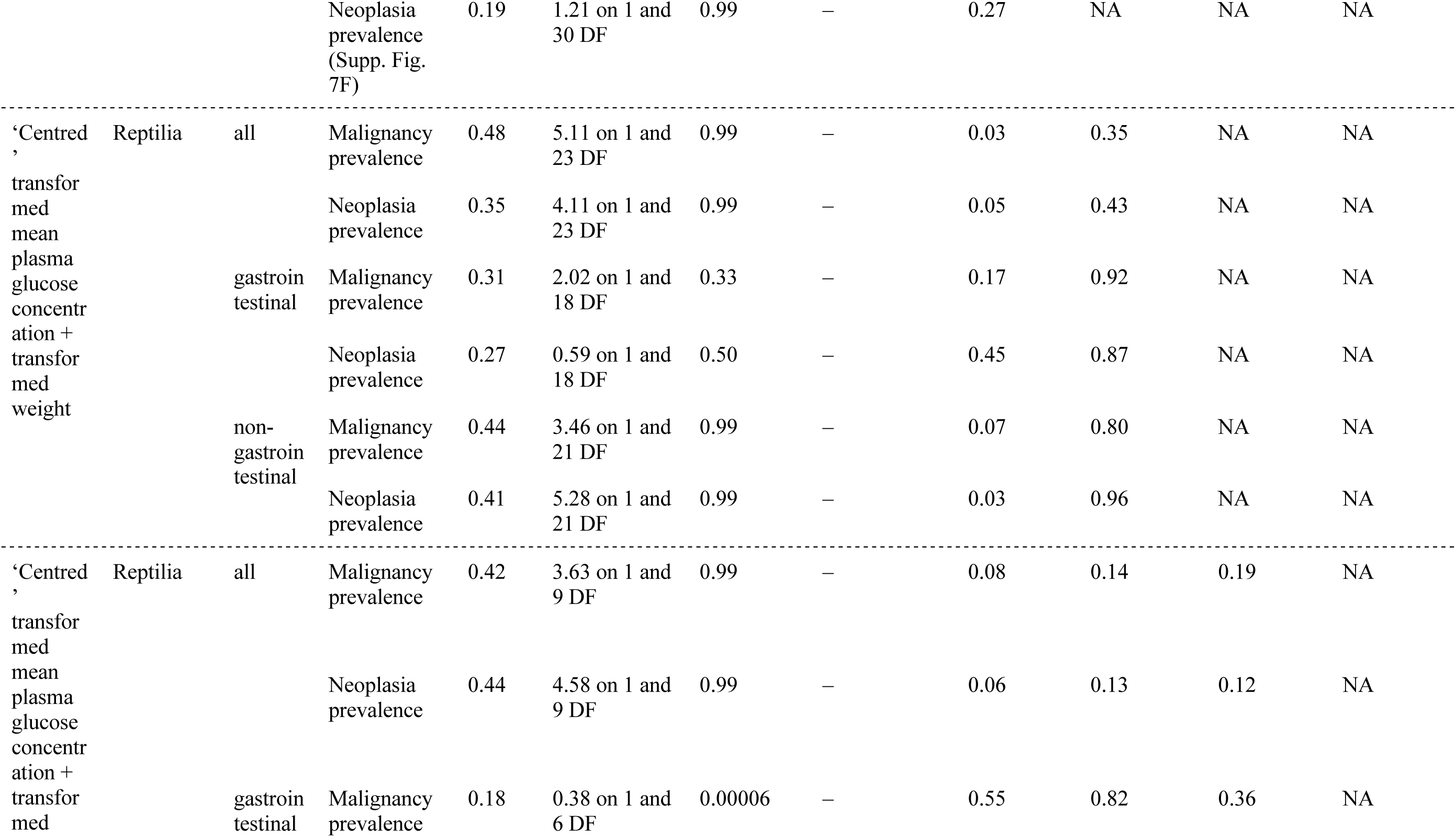
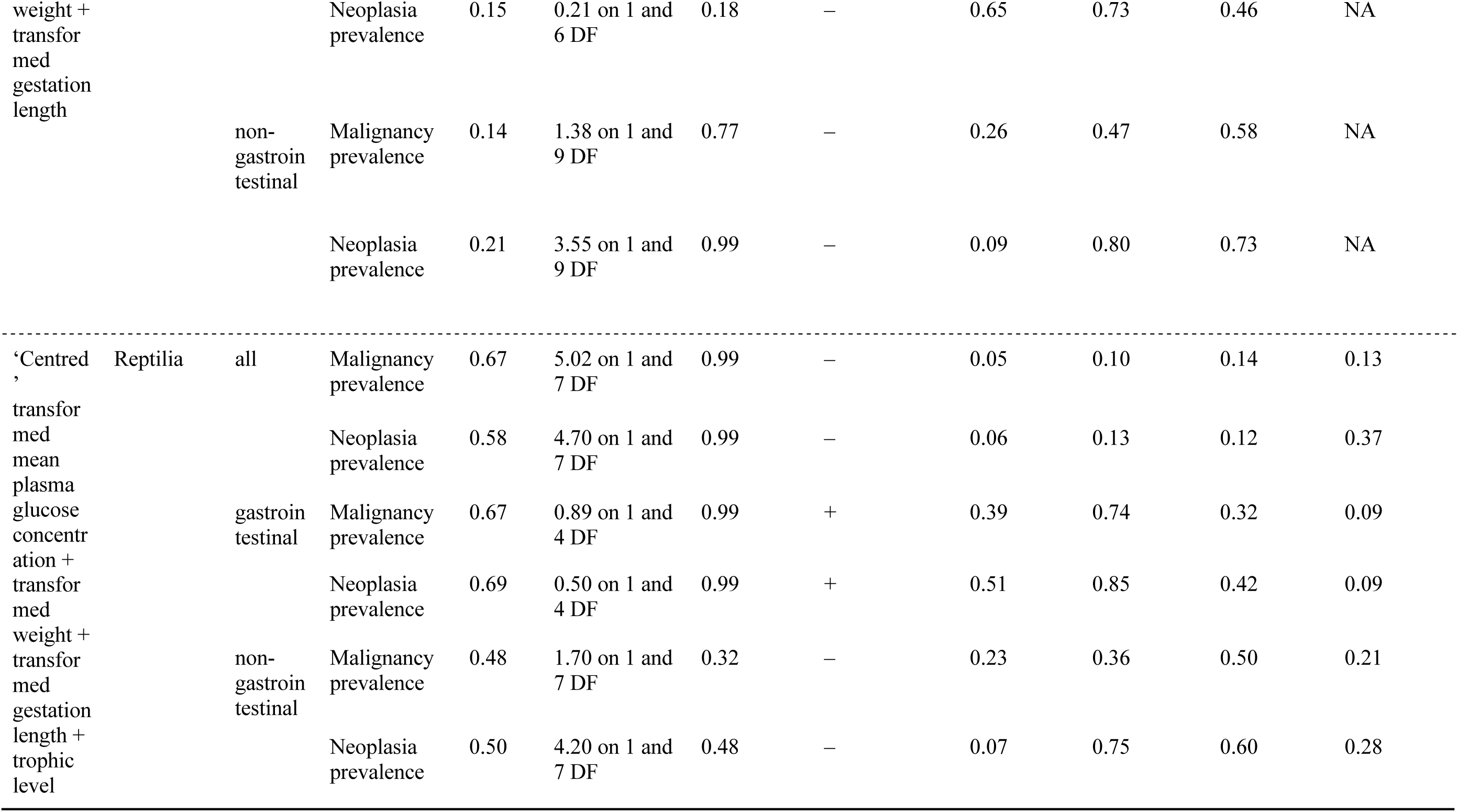
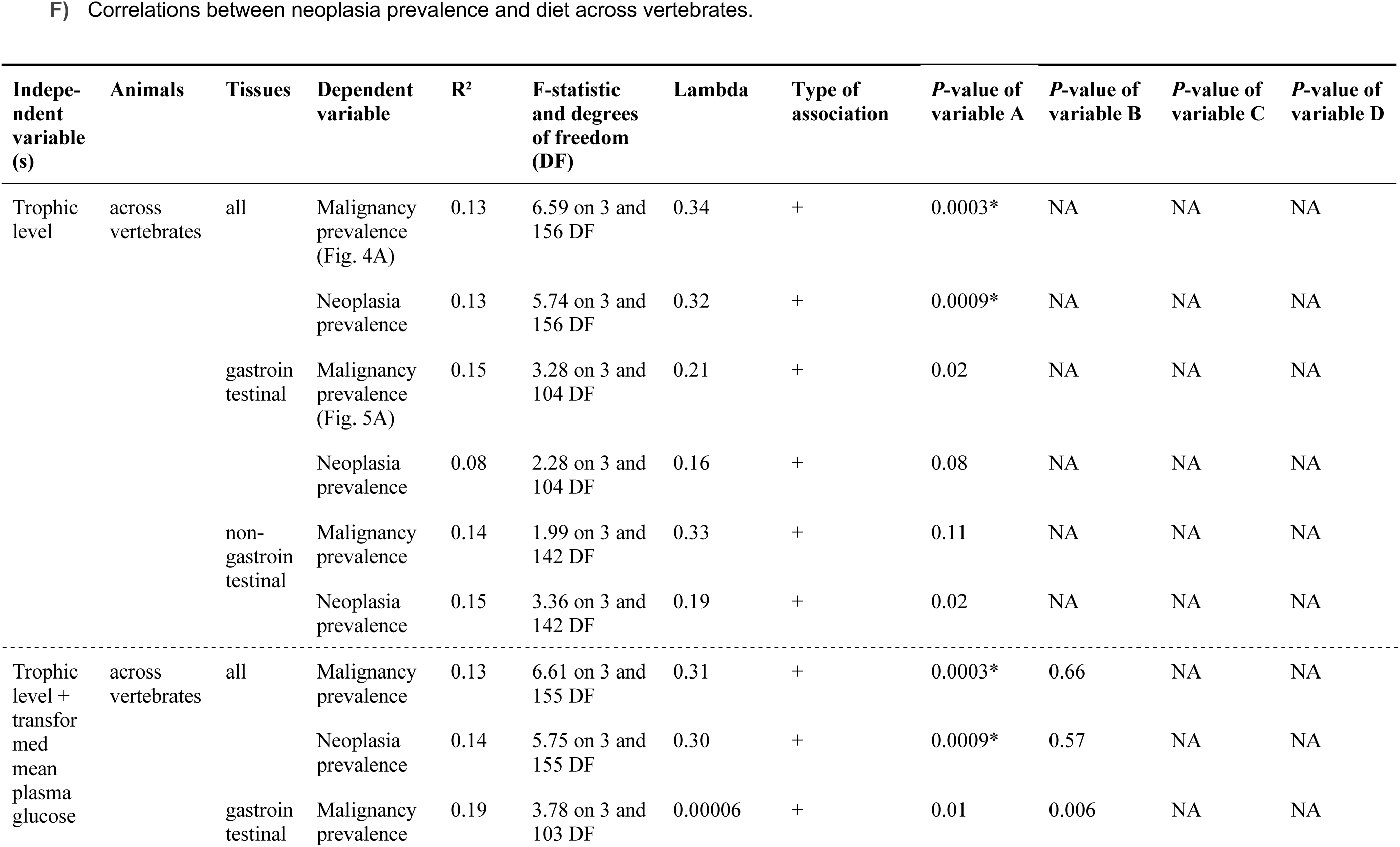
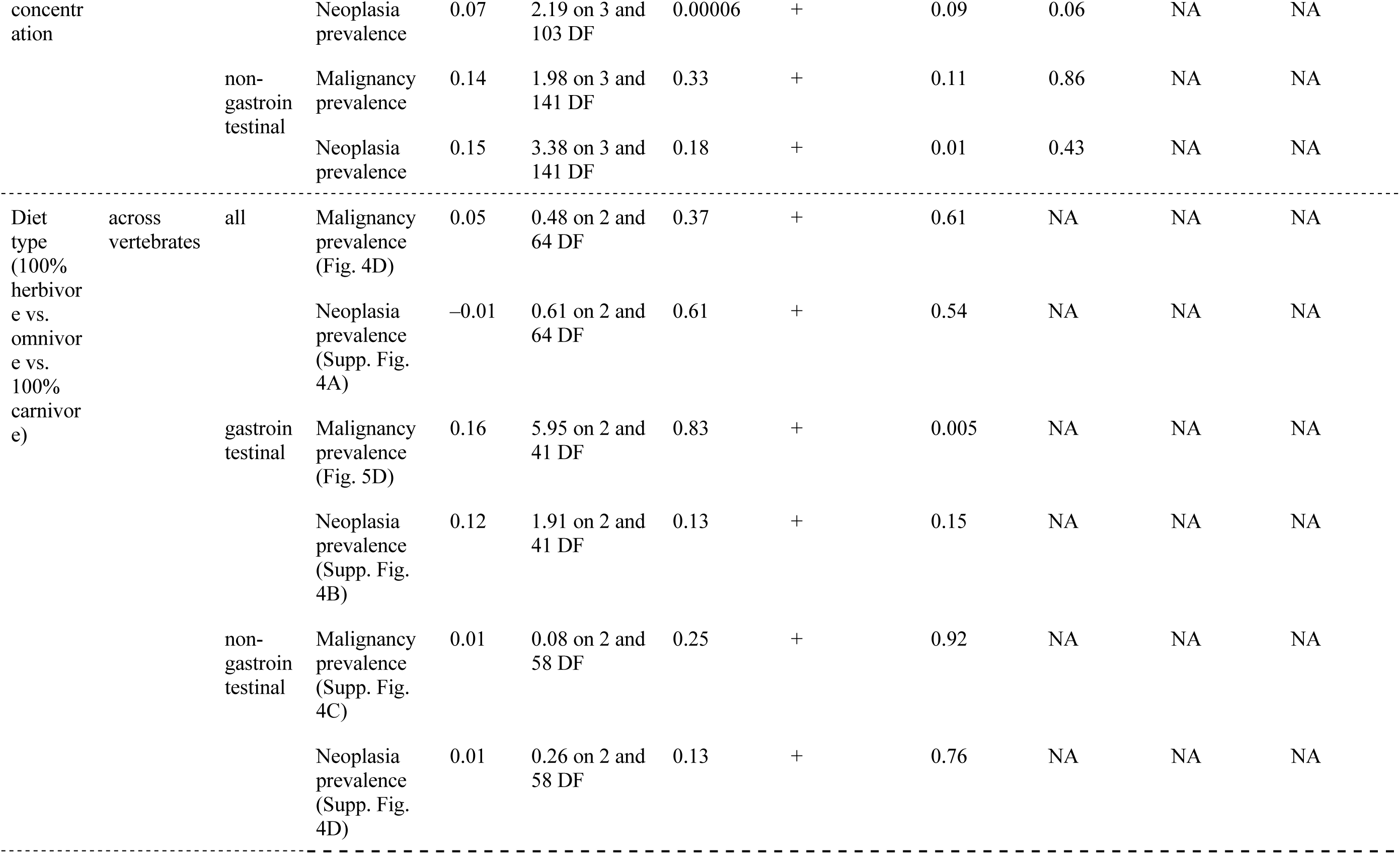
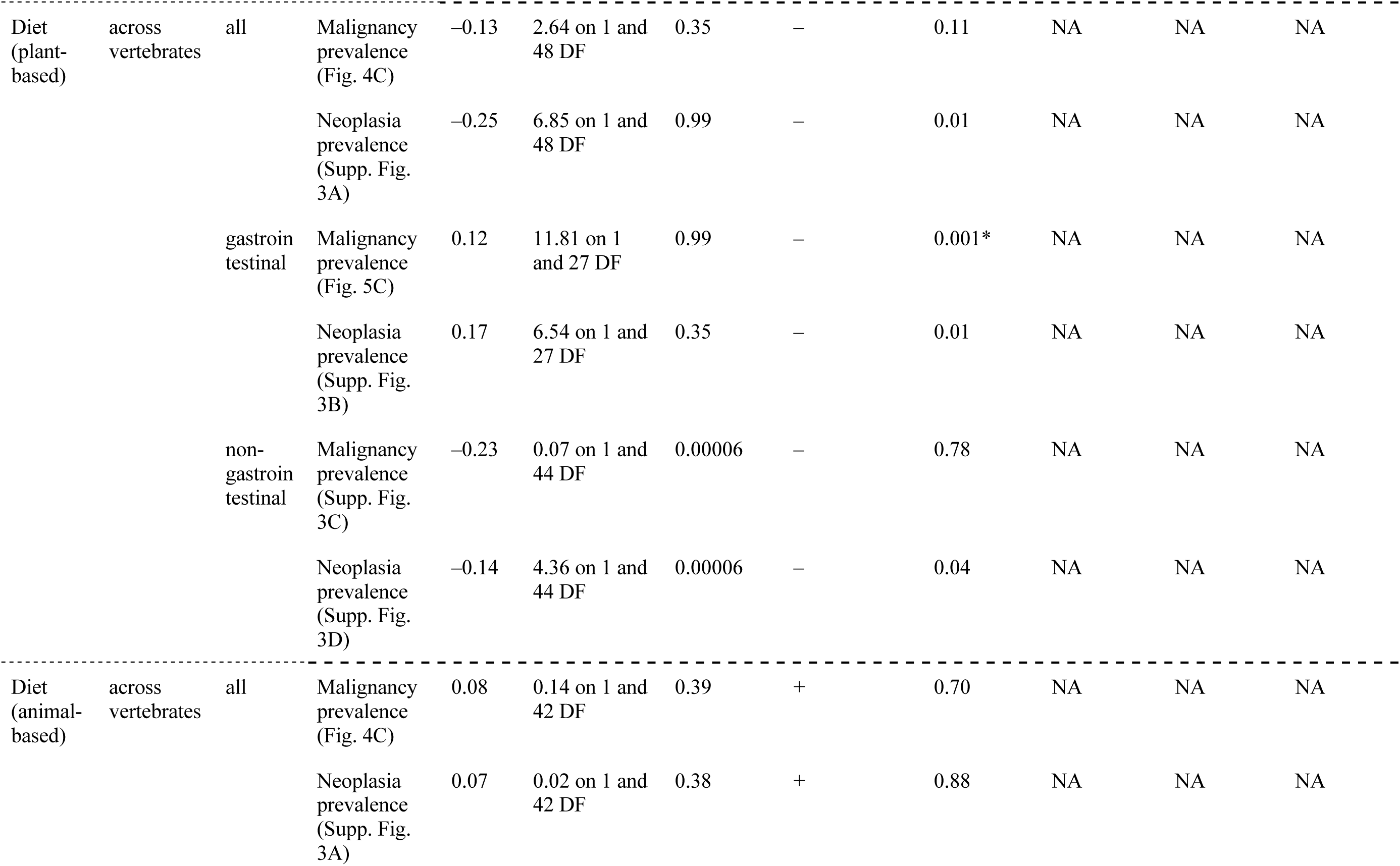
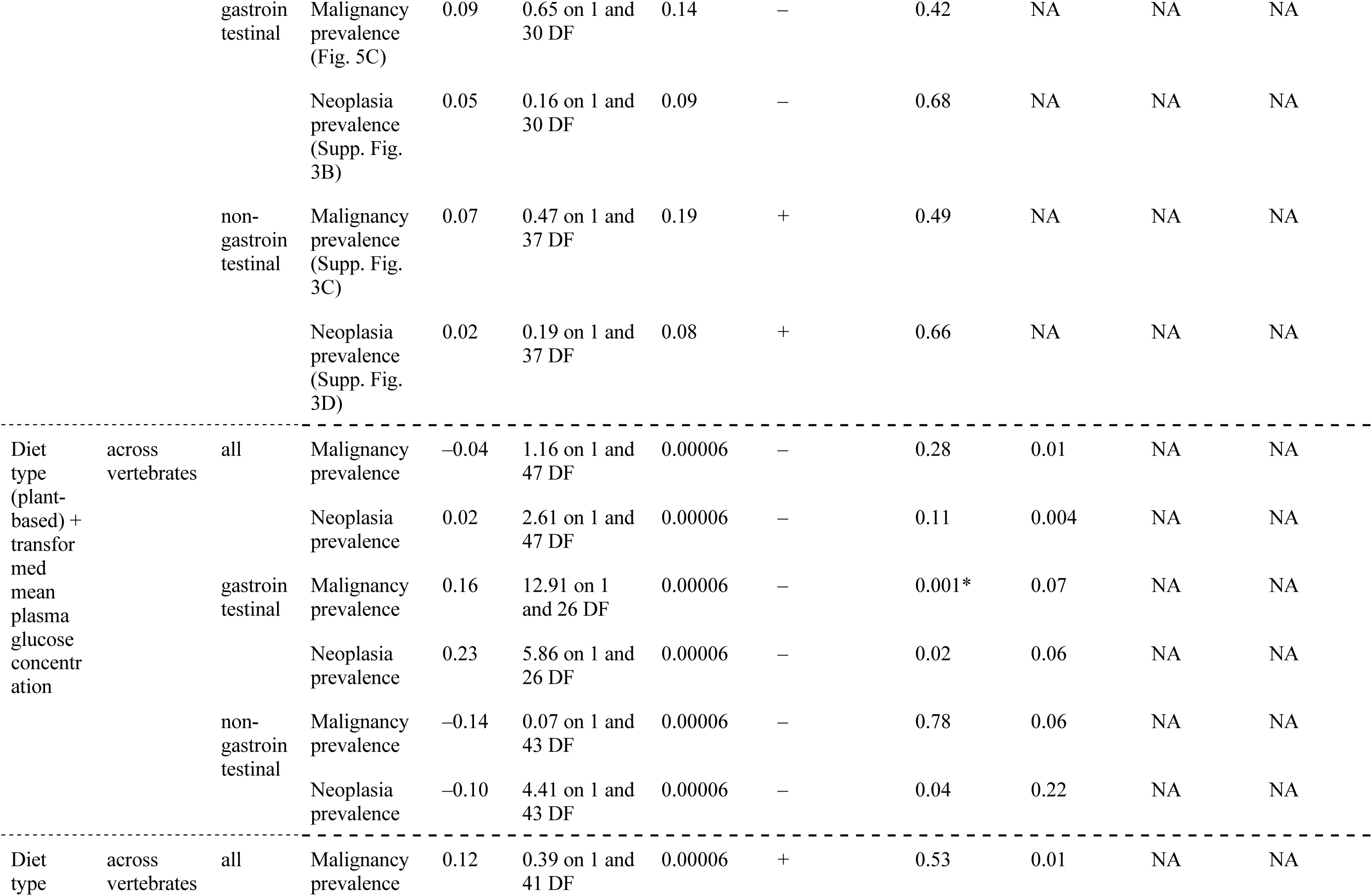
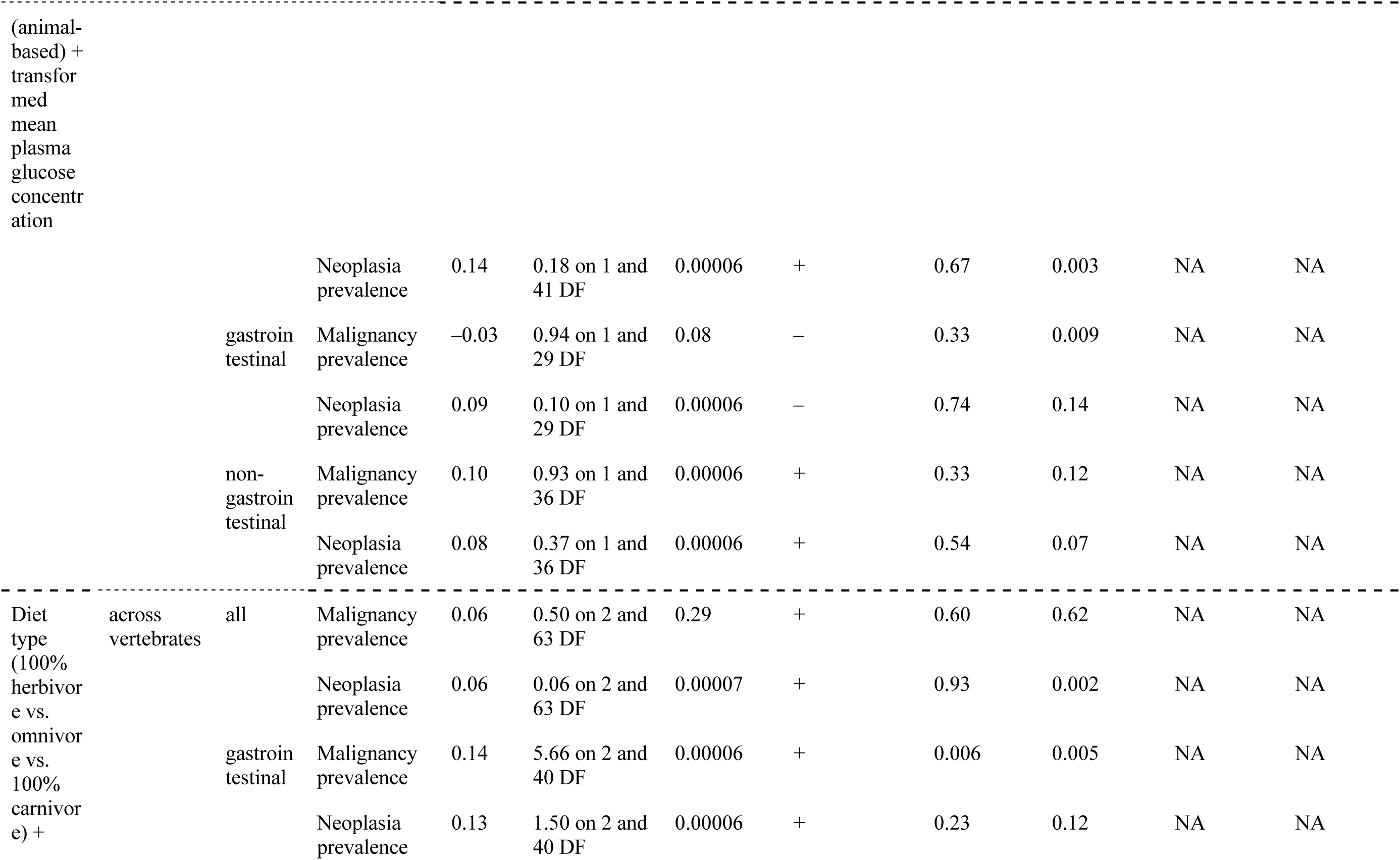
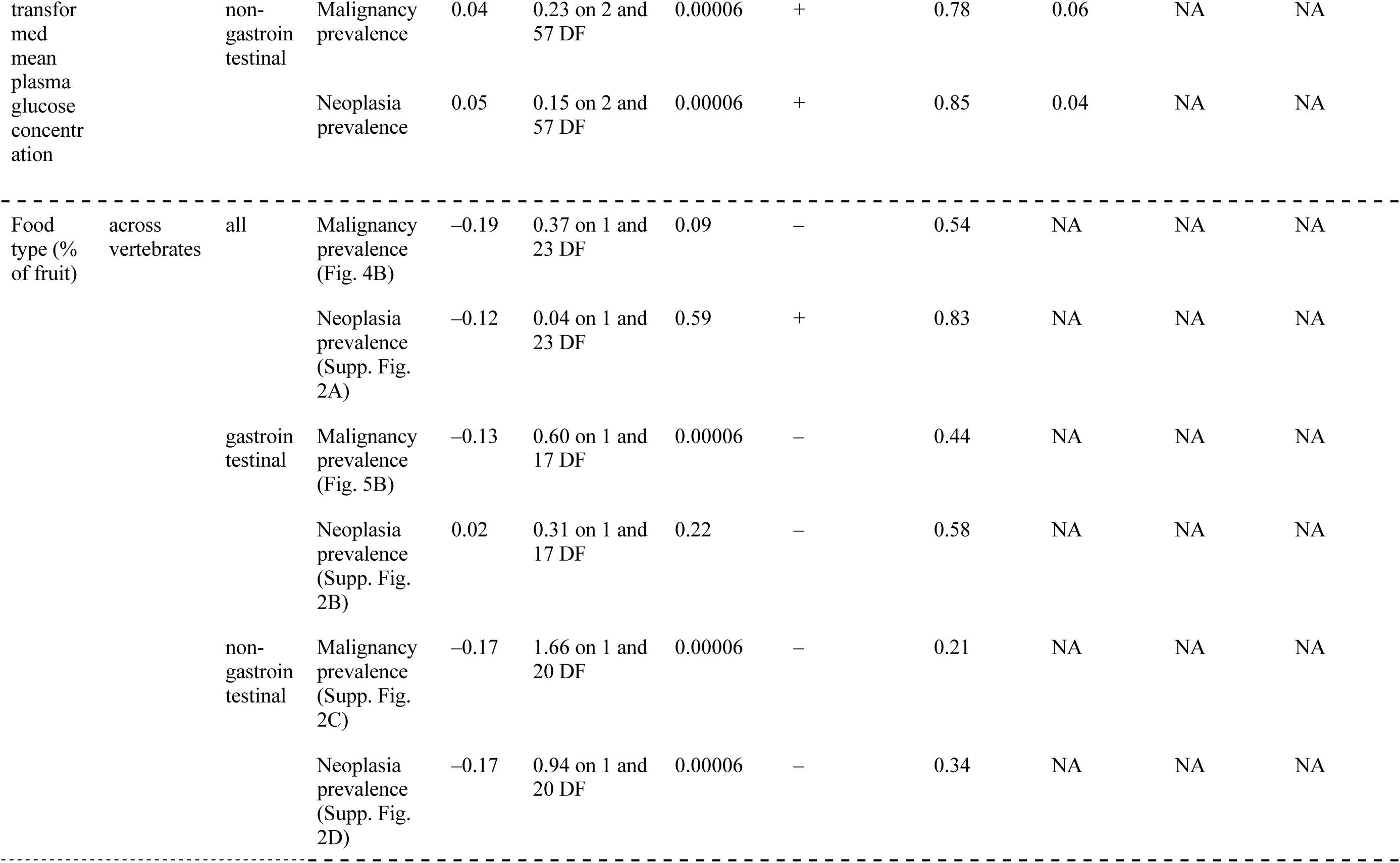
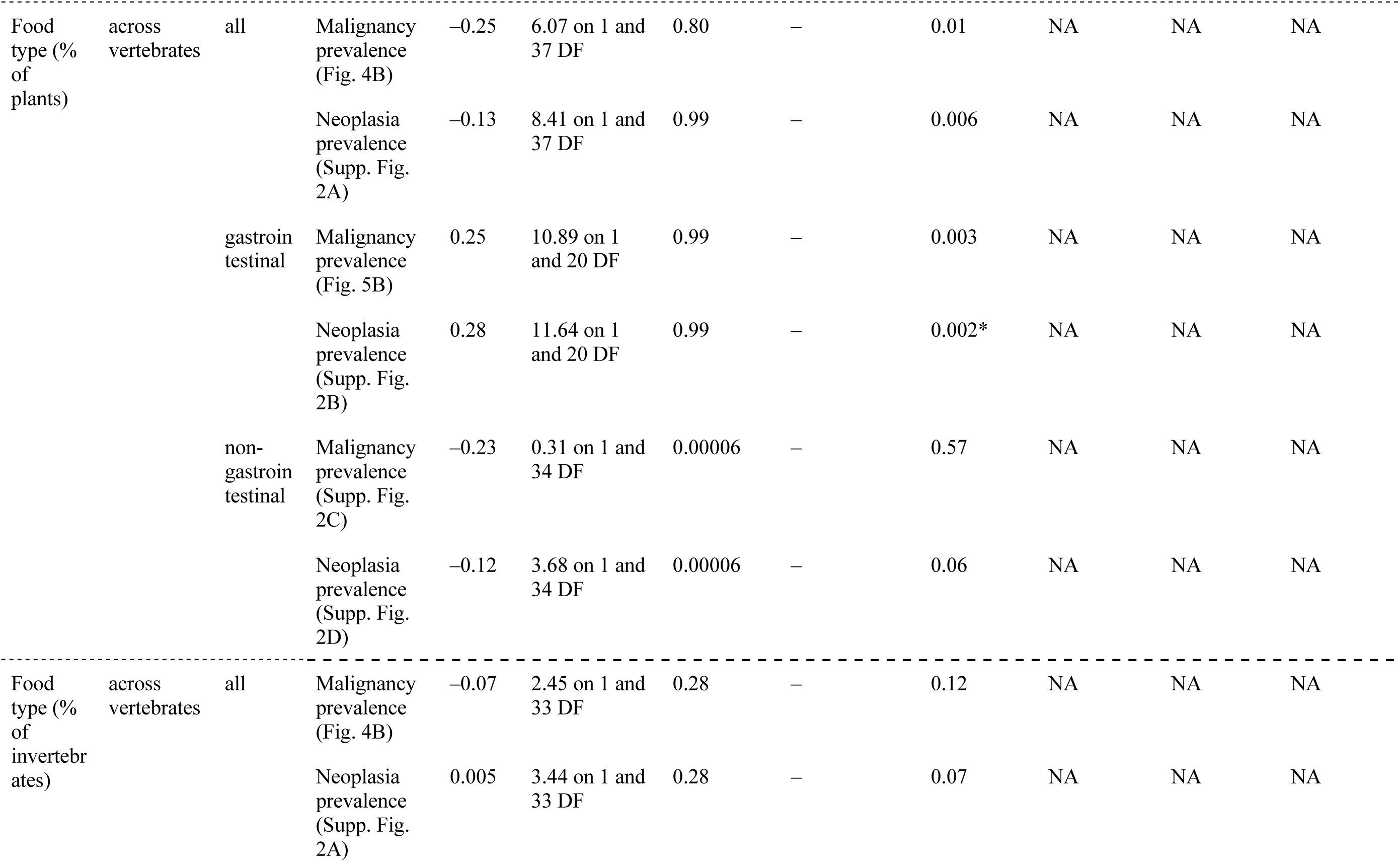
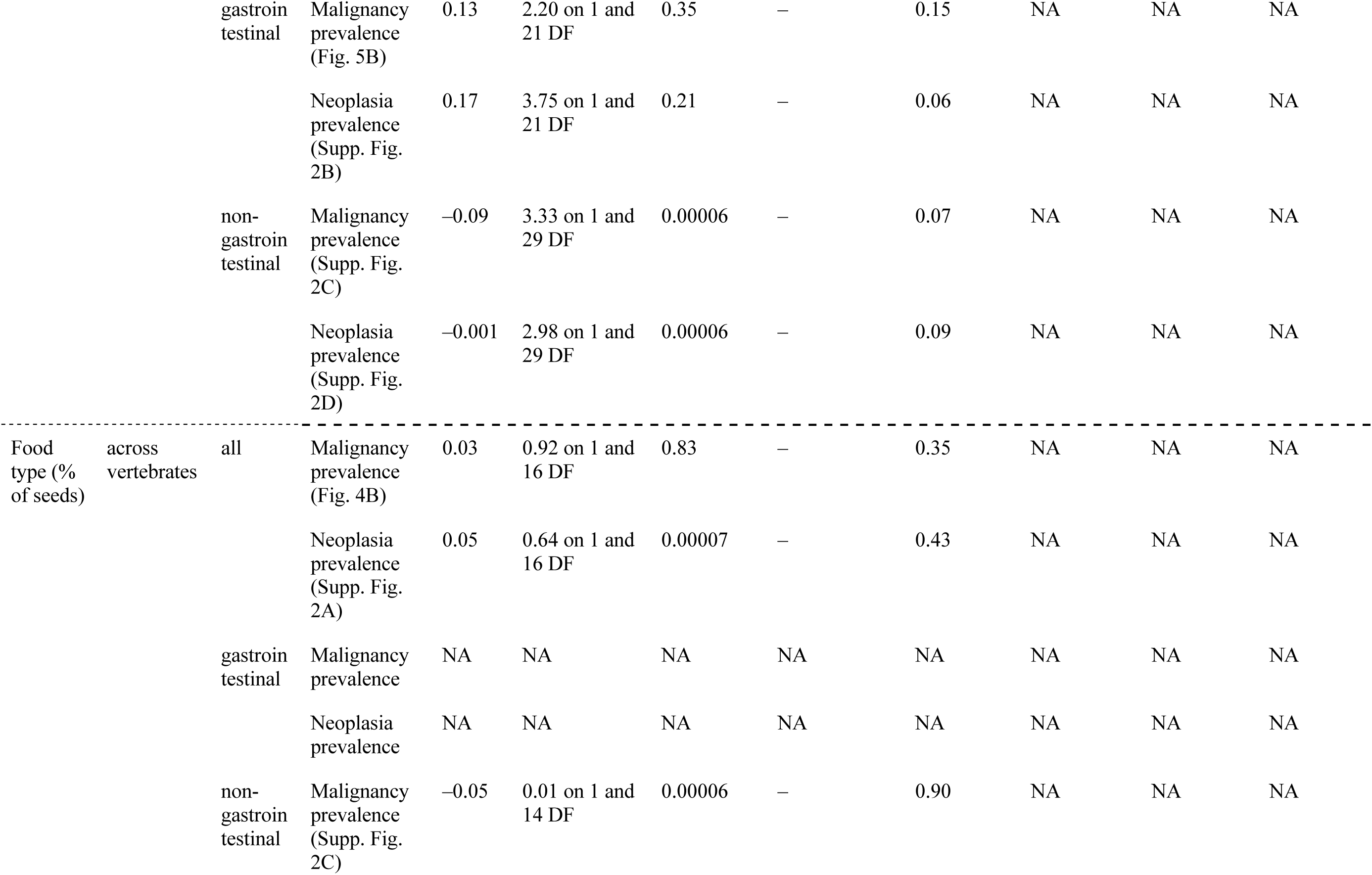
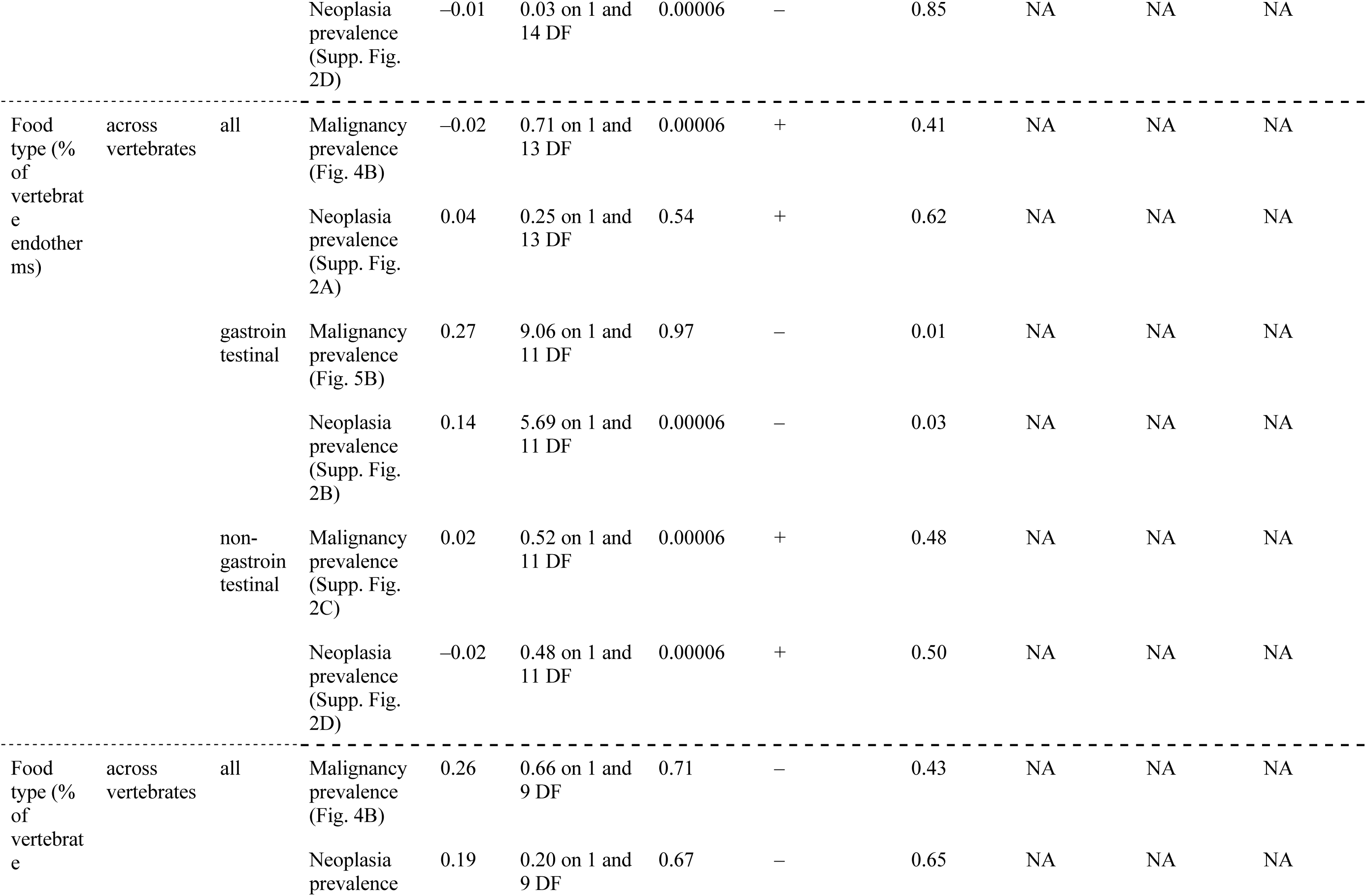
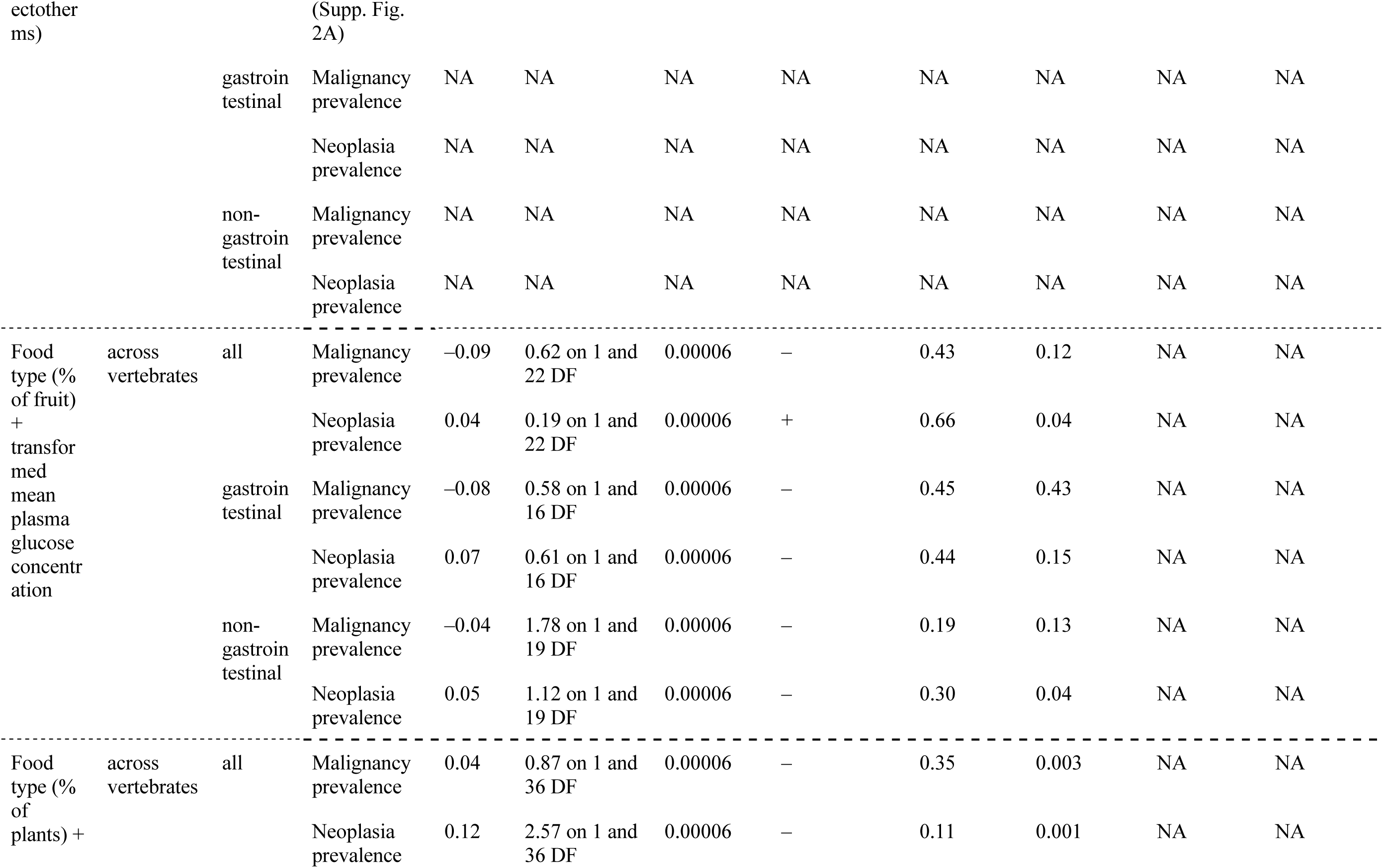
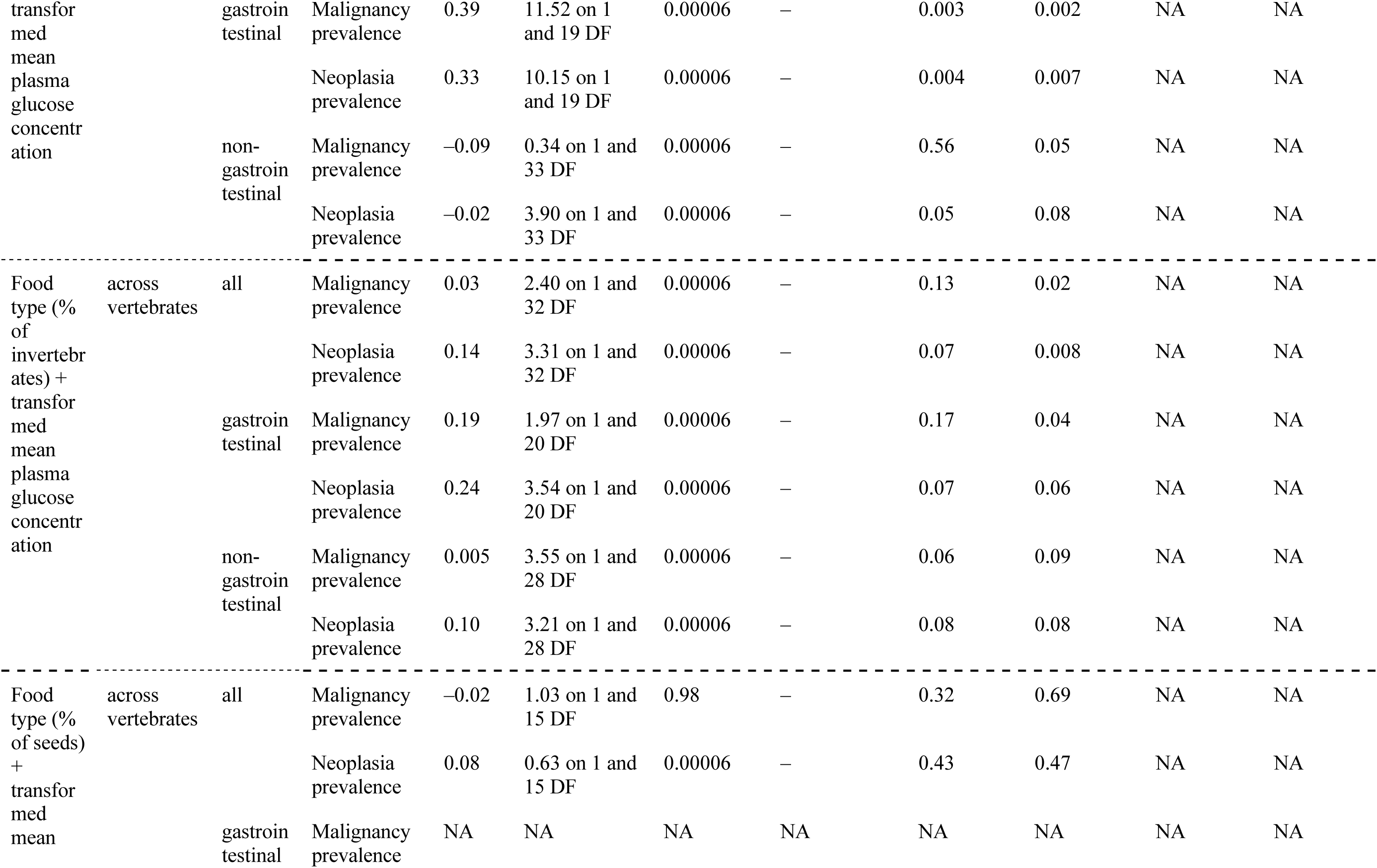
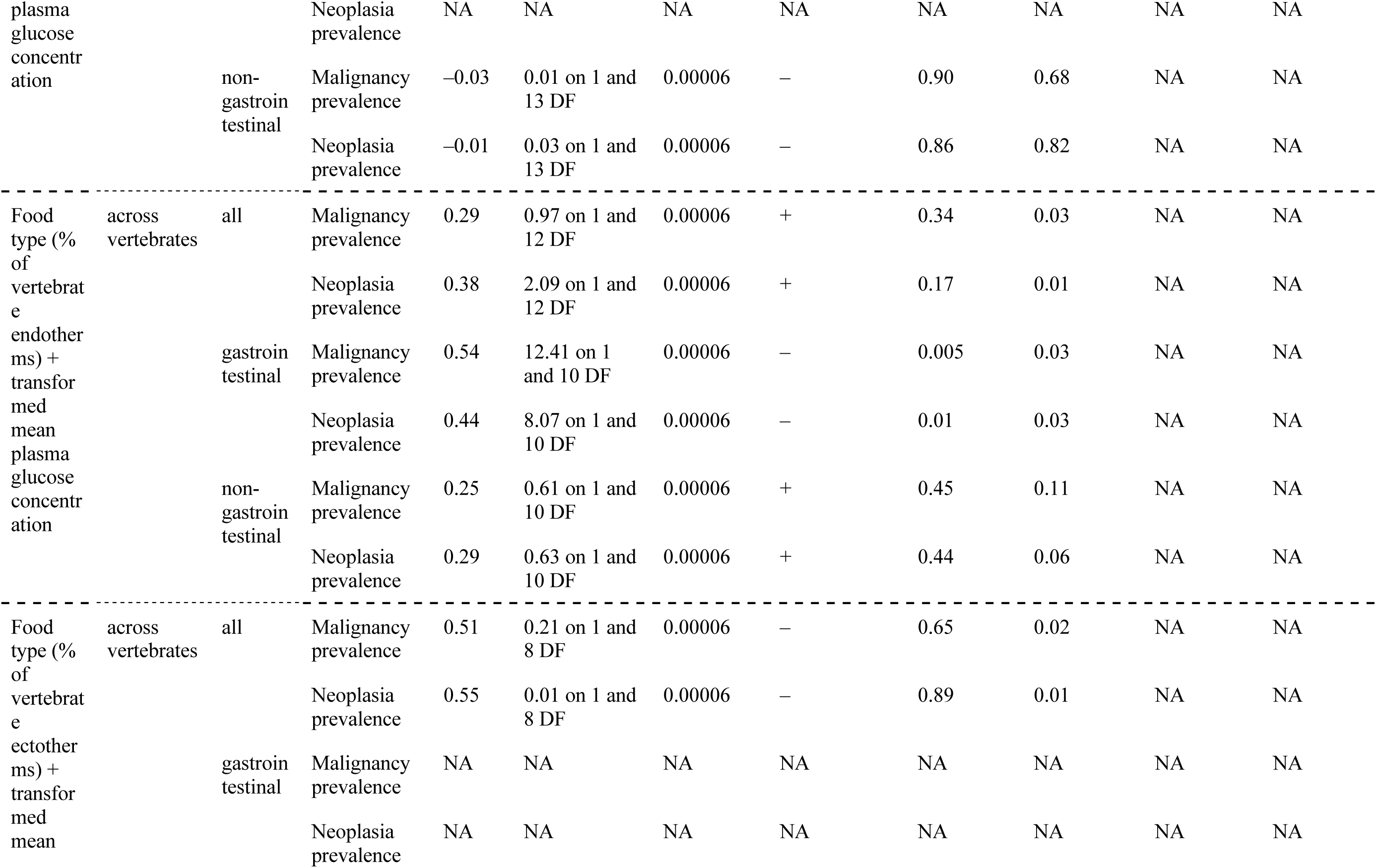
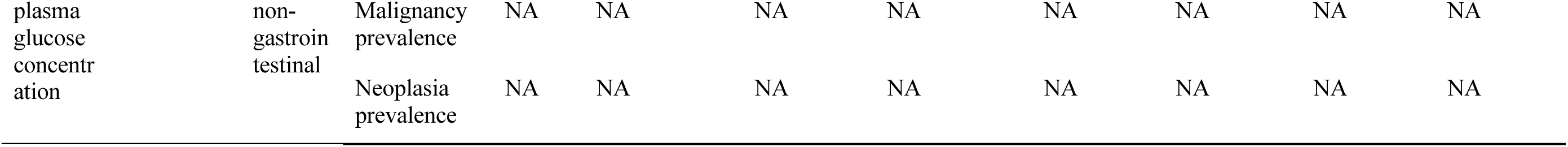
Phylogenetic generalised least squares (PGLS) regression summary results of the figures in the main text and in the supplementary materials. High values of lambda indicate that the signals can be mainly explained by common ancestry among species. In rows where all columns show NA, there were <10 species to conduct a powerful analysis. In the “Type of Association” column we report whether there is a positive (+) or negative (–) association between the independent variable A and the dependent variable. When the independent variable is categorical, we report the sign (+ or –) of the majority of between-group comparisons. If 50% of the between-group comparisons have a positive (+) association and 50% of the between-group comparisons have a negative (–) association, we report both signs. For multivariate analyses, in the 1st *P*-value column we report the *P*-value of the independent variable A, in the 2nd *P*-value column we report the *P*-value of the independent variable B, in the 3rd *P*-value column we report the *P*-value of independent variable C, in the 4th *P*-value column we report the *P*-value of variable D, and in the F-statistics column we report the F-statistics of the independent variable A. We mark *P*-values that passed the False Discovery Rate (FDR) correction in column “P-value of variable A” with an asterisk (*). Based on our hypotheses, we performed a separate FDR correction in the group of analyses as separated in the tables below (A-F).

**Figure 3.**
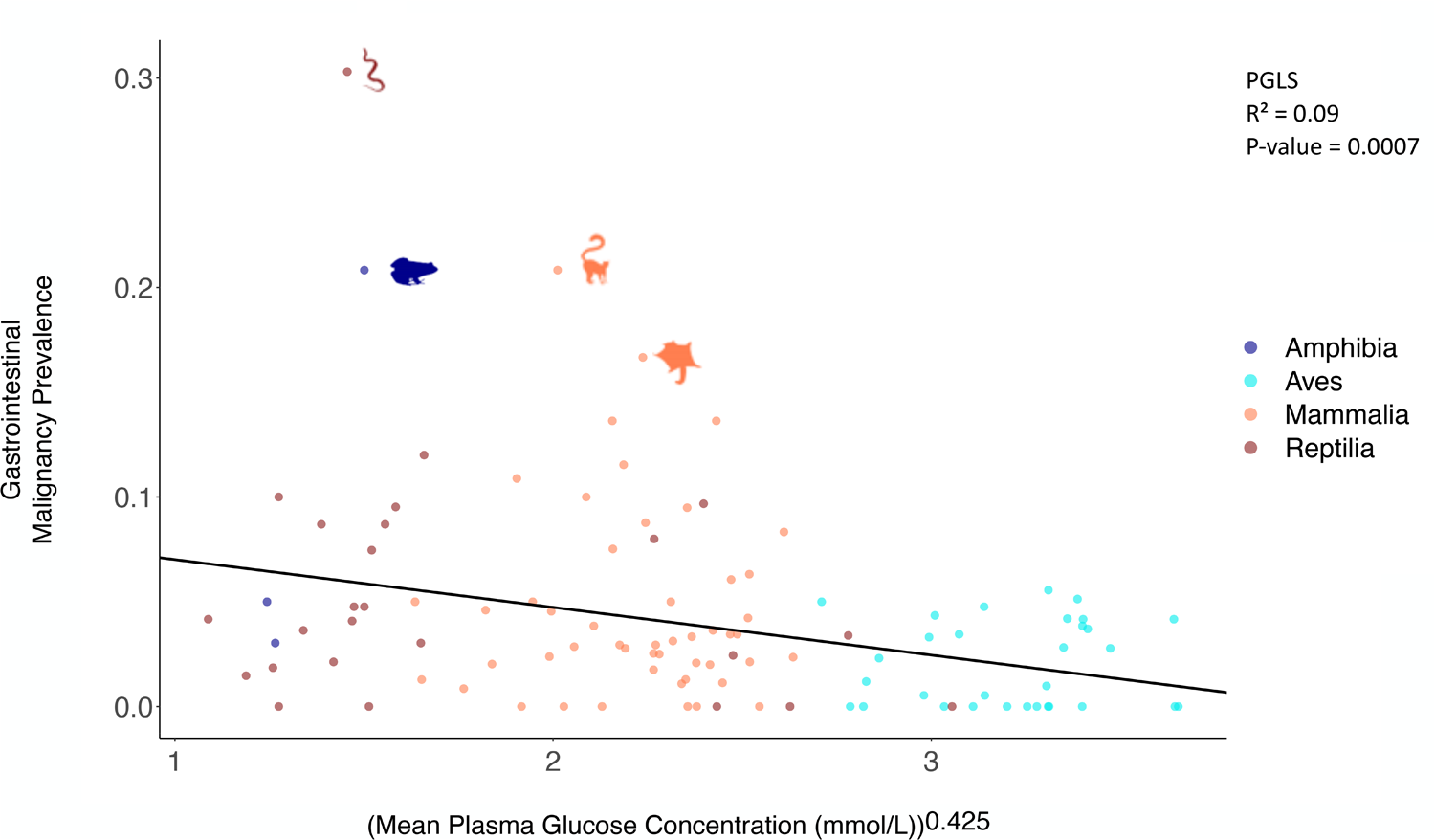
Relationship between mean plasma glucose concentration and gastrointestinal malignancy prevalence. The relationship (negative) was statistically significant (PGLS: *P*-value = 0.0007) across 108 species, even after correcting for multiple testing (Table 1B). Each dot represents the average plasma glucose concentration and the gastrointestinal malignancy prevalence of one species; Amphibia: Dark Blue; Aves: Blue; Mammalia: Orange; Reptilia: Red. We show images of significant outlier species (Rosner’s test). Animal silhouettes from PhyloPic (http://www.phylopic.org/).

### Relationships Between Cancer and Trophic Levels, Food Type, and Diet Type

Trophic levels are positively correlated with cancer prevalence and neoplasia prevalence across tissues among 160 species, with and without controlling for the variance in species’ plasma glucose concentrations (Fig. 4A), even after correcting for multiple testing (Table 1F). There is no significant correlation, however, between trophic levels and gastrointestinal malignancy and neoplasia prevalence, non-gastrointestinal cancer prevalence, or non-gastrointestinal neoplasia prevalence after applying corrections for multiple testing (Table 1F).

**Figure 4.**
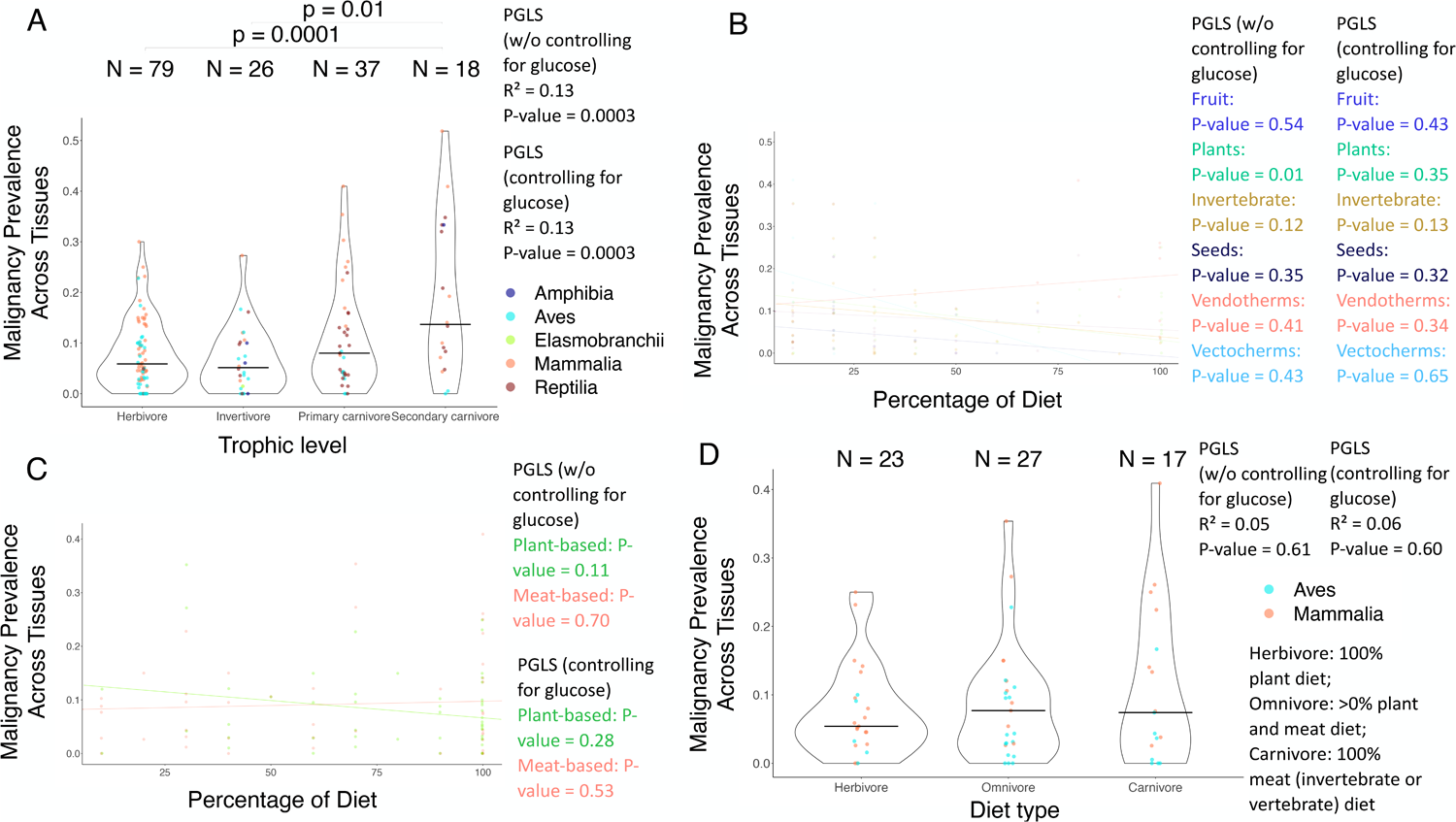
Relationships between overall cancer prevalence and diet across vertebrates. (**A**) Trophic level is positively correlated with malignancy prevalence across tissues for 160 species, even when controlling for variations in glucose concentrations in their plasma and correcting for multiple testing (PGLS: *P*-value = 0.0003; Table 1F; *P*-values ≤ 0.01 of between-trophic level comparisons are shown). **(B)** The percentage of plants in a species’ diet is negatively correlated with malignancy prevalence across tissues for 39 species only when not controlling for variations in glucose concentrations in their plasma (PGLS: *P*-value = 0.01). This result, however, does not remain significant when we correct for multiple testing. Also, there is no significant correlation between the percentage of fruit, invertebrates, seeds, endothermic vertebrates (Vendotherms), or ectothermic vertebrates (Vectotherms) in a species’ diet and malignancy prevalence across tissues for 25, 35, 18, 15, or 11 species, respectively (PGLS: *P*-value > 0.05). (**C**) The percentage of plant-based food in a species’ diet is not significantly correlated with malignancy prevalence across tissues for 50 species (PGLS: *P-*value > 0.05). Also, there is no significant correlation in the percentage of animal-based food in a species’ diet and malignancy prevalence across their tissues for 44 species (PGLS: *P*-value > 0.05). (**D**) Diet type is not significantly correlated with malignancy prevalence for 67 species (PGLS: *P*-value > 0.05). The horizontal black line in each trophic level (plot A) or diet category (plot D) shows the median malignancy prevalence across tissues in that category. Each dot shows the malignancy prevalence across tissues and diet category of a species. N shows the number of species per diet category (plots A & D). We added minimal jitter in the plots in order to better visualize individual data points.

The percentage of plants in a species’ diet is negatively correlated with gastrointestinal neoplasia prevalence among 22 species (Supp. Fig. 2B), but not after controlling for the variation in species plasma glucose concentrations (Supp. Fig. 2B; Table 1F). The percentage of fruit, plants, invertebrates, seeds, endothermic vertebrates, or ectothermic vertebrates in a species’ diet is not correlated with cancer prevalence and neoplasia prevalence across tissues (Fig. 4B; Supp. Fig. 2A), gastrointestinal malignancy prevalence (Fig. 5B), or non-gastrointestinal malignancy prevalence and non-gastrointestinal neoplasia prevalence (Supp. Fig. 2C, D) after correcting the analyses for multiple testing (Table 1F).

**Figure 5.**
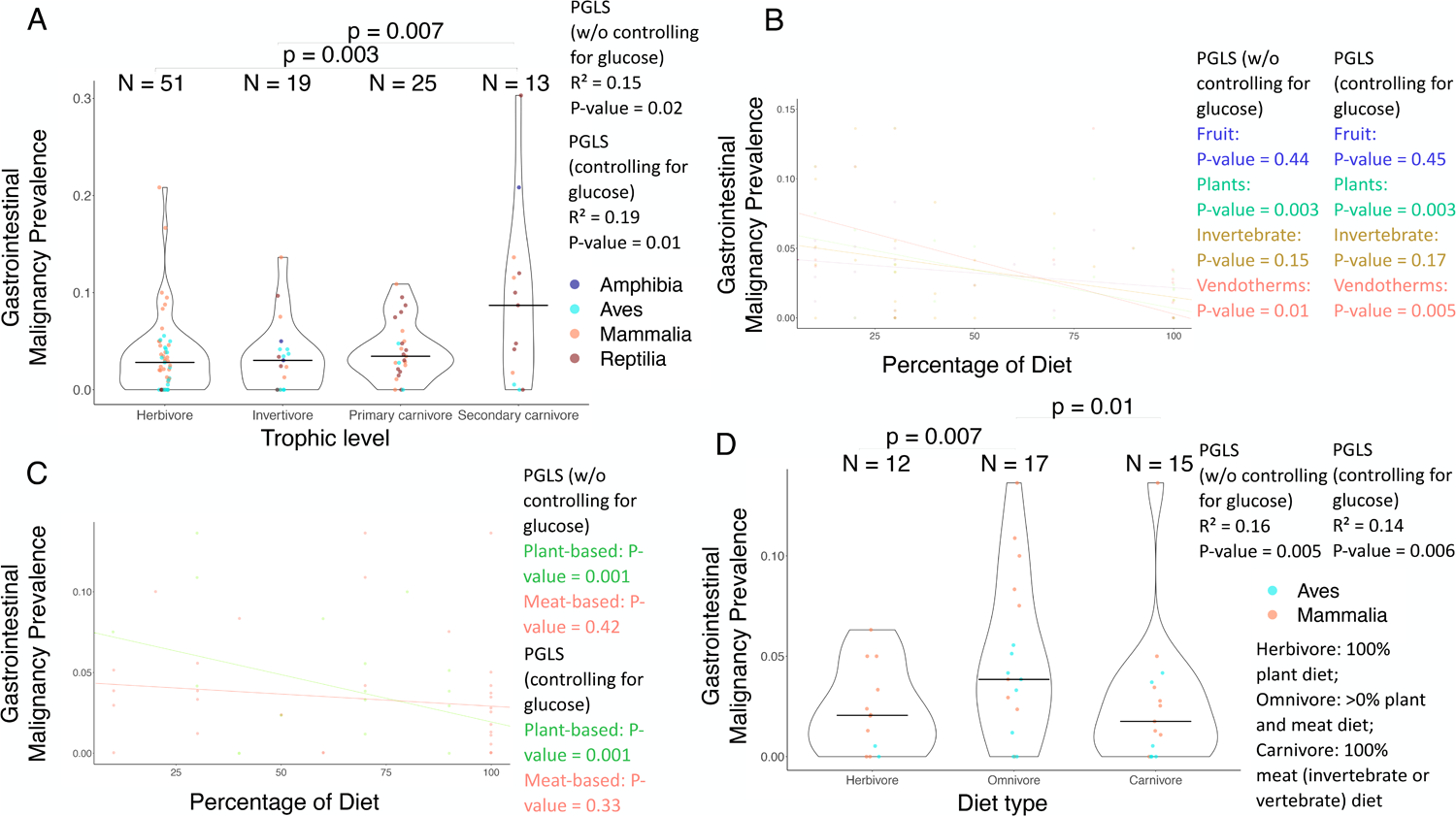
Relationship between gastrointestinal cancer prevalence and diet across vertebrates. **(A)** Trophic level is not significantly correlated with gastrointestinal malignancy prevalence for 108 species after correcting for multiple testing The percentage of plants in a species’ diet is negatively correlated with gastrointestinal malignancy prevalence for 22 species (PGLS: *P*-value < 0.05), but not after correcting for multiple testing (Table 1B; *P*-values ≤ 0.01 of between-trophic level comparisons are shown). **(B)**. The percentage of endothermic vertebrates (Vendotherms) in a species’ diet is negatively correlated with gastrointestinal malignancy prevalence for 13 species (PGLS: *P*-value < 0.05), but not after correcting for multiple testing. There is no significant correlation between the percentage of fruit or invertebrates in a species’ diet and gastrointestinal malignancy prevalence for 19 or 23 species, respectively (PGLS: *P*-value > 0.05). (**C**) The percentage of plant-based food in a species’ diet is negatively correlated with gastrointestinal malignancy prevalence for 29 species (PGLS: *P*-value = 0.001), even when controlling for the variance in the glucose concentrations in their plasma (PGLS: *P*-value = 0.001), and after correcting for multiple testing (Table 1B). There is no significant correlation in the percentage of animal-based food in a species’ diet and gastrointestinal malignancy prevalence for 32 species (PGLS: *P*-value > 0.05). (**D**) Diet type is significantly correlated with gastrointestinal malignancy prevalence for 44 species (PGLS: *P*-value < 0.05), but not after corrections for multiple testing (Table 1B). Omnivores have higher gastrointestinal malignancy prevalence than herbivores (PGLS: *P*-value = 0.007), and carnivores have lower gastrointestinal malignancy prevalence than omnivores (PGLS: *P*-value = 0.01). These correlations, however, do not remain significant after applying corrections for multiple testing (Table 1B). The horizontal black line in each trophic level (plot A) and diet category (plot D) shows the median gastrointestinal malignancy prevalence in that category. Each dot shows the gastrointestinal malignancy prevalence and diet category of a species. N shows the number of species per category (plots A & D). We added minimal jitter in the plots in order to better visualize individual data points.

The percentage of plant-based food in a species’s diet is negatively correlated with gastrointestinal cancer prevalence among 29 species even after controlling for the variation in the species’ plasma glucose concentration and correcting for multiple testing (Fig. 5C; Table 1F). However, the percentage of plant-based or animal-based food in a species’ diet is not significantly correlated with cancer prevalence across tissues and neoplasia prevalence across tissues (Fig. 4C; Supp. Fig. 3A; Table 1F), gastrointestinal neoplasia prevalence (Supp. Fig. 3B; Table 1F), or non-gastrointestinal cancer prevalence and non-gastrointestinal neoplasia prevalence, when correcting for multiple testing (Supp. Fig. 2C,D; Table 1F).

There is no association between diet type and cancer prevalence among 67 species (Fig. 4D), neoplasia prevalence across tissues among 67 species (Supp. Fig. 4A), gastrointestinal cancer prevalence among 44 species (Fig. 5D), gastrointestinal neoplasia prevalence among 44 species (Supp. Fig. 4B), non-gastrointestinal cancer prevalence among 61 species (Supp. Fig. 4C), or non-gastrointestinal neoplasia prevalence among 61 species (Supp. Fig. 4D), with or without controlling for the variance in plasma glucose concentrations and after applying corrections for multiple testing (Table 1F).

## Discussion

Similar to previous studies testing glucose concentrations in the plasma and whole blood across vertebrates^42,43^, we found that birds have the highest mean concentration of plasma glucose, followed by mammals, reptiles, and amphibia. In our cross-vertebrates diet, plasma glucose, and cancer association studies we found two main associations. Our results do not support our first hypothesis, as we found that diet is not correlated with mean plasma glucose concentration across vertebrates. Our results support our second hypothesis that there is a negative correlation between mean plasma glucose concentrations and neoplasia and cancer prevalence. Specifically, we found a negative correlation between mean plasma glucose concentrations and gastrointestinal neoplasia and cancer prevalence across vertebrates. We also found a negative correlation between mean plasma glucose concentration and non-gastrointestinal neoplasia prevalence only when controlling the analyses for weight. Our results also support our third hypothesis that trophic levels are positively correlated with cancer prevalence across tissues and neoplasia prevalence across tissues. This result holds with and without controlling for variations in plasma glucose concentrations across vertebrates. Also, the percentage of plant-based foods and the percentage of plants in a species’ diet are negatively correlated with gastrointestinal cancer prevalence and gastrointestinal neoplasia prevalence, respectively. Overall, these results indicate that mean plasma glucose concentration and trophic level/diet can be viewed as factors affecting cancer prevalence across species. We next explore a potential evolutionary explanation for our results, starting with why and how do birds have the highest mean plasma glucose levels across vertebrates.

Approximately 252 million years ago, during the Permian-Triassic boundary, there was a sharp decrease in oxygen concentration in the atmosphere^57,58^, from 30% to 11–15%^57,59^. This was followed by a gradual increase in the concentration of atmospheric oxygen in the late Jurassic, Cretaceous, and Tertiary^57,59^. According to Takumi Satoh, these environmental conditions led to a divergence in the metabolic pathways of birds relative to mammals and reptiles^60^. Specifically, there was a reduction in bird genome size and loss of some genes (e.g. encoding the protein omentin^61^) or absence of the GLUT4 immunoreactive glucose transporter protein in most tissues^42,62^, reducing import of glucose inside the cells, and providing protection from hypoxia^60^. Given this loss or downregulation of genes related to glucose import inside bird cells, among other adaptations, today there is a higher plasma glucose concentration in birds versus other vertebrates^42–44^ (Fig. 2 & 3; Supp. Fig. 1; 50 out of the 78 vertebrate species with the highest glucose levels are birds: Supplementary data), which does not change significantly with diet^29,30^.

The reduced import of glucose inside avian cells has multiple intracellular effects that contribute in stabilizing and protecting DNA, suppressing cellular growth, and slowing down ageing and ageing-related diseases in birds relative to mammals^63,64^ and reptiles^65^. First, there is comparatively lower intracellular storage of glucose as glycogen in birds^42^. Second, there is caloric restriction inside bird cells^66^, and we know that caloric restriction or “eating [living] like a bird”^63^ is one of the strategies humans have implemented in order to increase human longevity and decrease the risk of cancer^67,68^. Third, the constitutive inactivation of the ageing hormone insulin/IGF-1^69,70^ approximately doubles bird lifespan^60^. Fourth, birds lack certain inflammatory cytokines such as TNF-a^71,72^. Fifth, birds have evolved longer telomeres. Sixth, birds have adapted to thrive on fatty acids rather than glucose, mostly during long-distance flights and migration^73,74^. Seventh, birds have adaptations such as being more physically active; more than 99% of species of birds fly^63^. Eighth, birds have erythrocytes with a nucleus and high metabolic activity, thus increasing their intracellular concentration of haemoglobin relative to mammals and reptiles, and maximizing the use of the limited oxygen in the atmosphere during the Permian-Triassic^60^. Ninth, birds are known to have higher metabolic rates than most mammals^60,75–78^. In a study across vertebrates, including birds and mammals, higher inferred metabolic rates were correlated with lower neoplasia prevalence though these results did not remain significant after multiple testing corrections^13^. Tenth, birds produce fewer reactive oxygen species via mitochondrial respiration in comparison to mammals and reptiles^65^. Eleventh, there are higher levels of the potential antioxidant “uric acid” in birds than mammals^79,80^. Twelvth, the antioxidant NRF2 is constitutively expressed in most birds as an evolutionary consequence of the loss of a binding domain in its repressor KEAP1 during the Permian-Triassic period^60^. These features may explain the increased lifespan^44,81^ and lower cancer prevalence^18,45,82,83^ in birds relative to mammals and reptiles, and our findings that vertebrate species with higher plasma glucose concentrations have lower gastrointestinal neoplasia and cancer prevalence. These features may also explain why fowl, which have maintained a functional KEAP1, are amongst the birds with the highest cancer and neoplasia prevalence^8^ (Supplementary Data; Supp. Fig. 6A, 6E–F).

The relationship between higher plasma glucose levels and lower gastrointestinal neoplasia and cancer prevalence, however, may not be entirely explained by phylogenetic relatedness. As seen in previous studies, a mammal, the Egyptian fruit bat, has around 5X higher blood glucose levels than a healthy rat^84^. Further explanations for this negative correlation between glucose levels and gastrointestinal neoplasia and cancer prevalence, apart from phylogeny, may be a shared ancestral environment by some of these species. This may mean that they share a similar microbiome, which is known to affect metabolic rates in some taxa^85–87^, and thus may also affect blood glucose levels. A shared environment may also mean that the animals are exposed to similar pollutants affecting their cancer prevalence. Also, the less host-specific gut microbiome of birds and bats, and the potential subsequent higher proportion of nutrients absorbed directly by bird and bat cells, relative to other vertebrates^88^, may explain the relatively constant high blood glucose levels in birds and bats. Additionally, a non-selective force of evolution, random genetic drift acting during domestication may have led to the decreased levels of circulating glucose in bovids^89^ and possibly other domesticated species, relative to their ancestors or close relatives in the wild. Domestication can also increase the risk of cancer in animals via the effects of inbreeding depression and the inability of selection to remove deleterious alleles^90^. So, apart from phylogeny, a shared environment/microbiome and domestication may explain the observed negative correlations between mean plasma glucose concentration and gastrointestinal neoplasia and cancer prevalence across vertebrates, though this hypothesis remains to be tested.

The majority of primary (28/36 species = 77.7%) and secondary carnivores (14/17 species = 82.3%) in our dataset are mammals and reptiles (Supplementary data). Mammals and reptiles have previously been shown to have lower median blood glucose levels than birds^43^. Given the above-mentioned association between the regulation of plasma glucose levels, the switch to fatty acid metabolism inside cells in birds, and the downstream antioxidant effects that increase longevity and decrease cancer prevalence in birds, this relative abundance of mammals and reptiles in higher trophic levels may explain the higher cancer prevalence in higher trophic levels^13^, including species with a lower percentage of plant-based food in their diet (Table 1F). Additionary, it is known that toxins bioaccumulate at higher concentrations in higher trophic levels^91–93^. In other words, lower trophic levels have relatively fewer bioaccumulated toxins. Also, a study in 95 birds has shown that eating invertebrates or a higher seed-to-fruit ratio was positively correlated with higher Trolox-equivalent antioxidant capacity and higher concentration of the potential antioxidant uric acid^94^. Overall, 1) the relative abundance of reptiles and mammals in higher trophic levels; 2) the biomagnification of toxins in higher trophic levels; and the 3) oxidative stress and DNA damage associated with eating red meat, may explain the higher cancer prevalence in higher trophic levels^13^ (Table 1F).

### Limitations & Future directions

There may be some mismatches between the diet, mean plasma glucose concentration, and cancer prevalence of species in our dataset versus the actual values of diet, mean plasma glucose concentration, and cancer prevalence in a species because each of these three components were most likely collected from different individuals of the same species and we do not known the exact health status of every individual at the time of data collection. For example, the mean plasma glucose concentrations in our study were collected from anesthetized animals (data from ZIMS), some of which may have had infections that affected their blood glucose levels, may have had higher blood glucose levels due to diabetes^95^, and/or may have been stressed during the method of capture (e.g., being darted, netted, or grabbed). Anaesthesia or capture can mildly to moderately elevate the blood glucose levels of felines^96^, dogs^97^, cattle^98^, goats^99^, mice^100^, rats^101^, chickens^102^, pigeons^103^, eagles^104^, American kestrels^105^, and possibly other species, but not of red-tailed hawks^106^. Therefore, it is important for future studies to examine whether our results hold when measuring the health status, diet, and plasma glucose levels of the exact same individuals, either in zoological institutions or from the wild.

In addition, some data from the 160 species in our dataset are missing. Specifically, we only know the gastrointestinal neoplasia prevalence of 108 species, the non-gastrointestinal neoplasia prevalence of 146 species, the adult weight of 142 species, the gestation length of 104 species, and the diet type or diet percentage of 67 species (Supplementary data). Including more vertebrate species in the dataset will help draw more robust conclusions about the associations between trophic level (>10 species in every trophic level), diet type or dietary percentage, mean plasma glucose concentration, and cancer prevalence within and across clades.

When trying to understand a biological phenomenon, both the *why* and *how* questions are important^107^. We have explained a hypothetical selective force, such as low atmospheric oxygen levels during the Permian-Triassic boundary, that may have shaped the divergence in glucose import and cancer prevalence in birds versus mammals and reptiles, but we have a long way to go to completely understand how gastrointestinal neoplasia and cancer prevalence is linked to lower mean plasma glucose levels. How do mean plasma glucose levels not change with diet? How exactly are trophic levels associated with neoplasia prevalence? Many molecular studies describe the links between diet, plasma glucose concentrations, and cancer prevalence, as mentioned in previous paragraphs. These molecular studies, however, have not been confirmed in every single species in our dataset. Comparative genomics^108,109^ and transcriptional analyses can provide some answers on the diet-, glucose-, and cancer-related gene pathways that are present/absent or differentially regulated in different taxa.

### Conclusion

To our knowledge, this is the first time an association between plasma glucose levels and gastrointestinal neoplasia and cancer prevalence has been reported across vertebrates. The fact that this association is negative indicates that these results may be partly explained by Satoh’s framework on the divergence of birds versus mammals and reptiles during the Permian-Triassic boundary^60^ and/or a possibly higher uptake and faster metabolism of glucose by healthy avian bird cells relative to healthy mammalian^110^ and reptilian cells. Future studies should test whether our correlations across species hold when measuring all three components (diet, plasma glucose levels, and cancer prevalence) from the exact same individuals, either in animals under human care or wild animals, and broadening these comparisons to many more vertebrate species. The rise of comparative phylogenomics^108,109^ can also bring insights into how particular glucose-transport-related genes and pathways are associated with cancer-related genes and pathways across the animal kingdom. These approaches will bring us closer to explaining the diversity of cancer prevalence across species and designing more targeted gene therapies that possibly help us ‘live like a bird’.

## Methods

### Data Collection and Classification

#### Trophic Level, Food Type, and Diet Type Data

We identified trophic levels (herbivore, invertivore, primary carnivore, or secondary carnivore) for n=160 species based on records from previous studies (classified for each species based on their primary diet)^13^. We collected food type percentages (endothermic vertebrates, ectothermic vertebrates, fish, fruit, invertebrates, nectar, plant, scavenger, seed, and unknown vertebrates) from Jim Song et. al. (2020) for n=67 species^88^. In the analyses of these food type percentages (with a percentage of >10%), we removed food types (fish, nectar, scavenger, unknown vertebrates) with a sample size smaller than 10 species to improve the power of the analyses. Next, in the analyses of the percentage of animal-based vs. plant-based foods, we summed the food type percentage to determine the percentage of animal-based (invertebrates, endothermic vertebrates, ectothermic vertebrates, unknown vertebrates, scavenger, and fish) and plant-based foods (fruit, nectar, seed, and plant) in each diet for each species. Lastly, we assigned diet types (carnivore: 100% animal-based diet, herbivore: 100% plant-based foods, and omnivore: contains both animal- and plant-based foods) based on the food type percentage. In all the following analyses, we excluded species for which the percentage of that food type in their diet was 0% because: (1) we were interested in the effects of eating a particular food type on neoplasia prevalence and plasma glucose concentrations; and (2) one can already identify the 0% data from our figures and analyses of the other diet types - for example, if a species consumes 100% fruit, it does not consume any of the other food types, in other words the consumption of endothermic vertebrates, ectothermic vertebrates, fish, invertebrates, nectar, plants, scavenging items, seeds, and unknown vertebrates is 0%. We then matched trophic level, food types, animal/plant-based percentage, and diet types to cancer data for analysis.

#### Plasma Glucose Concentration, Cancer, and Life-history Data

We collected records of mean plasma glucose concentrations (measured in mmol/L) for each species as recorded in the ZIMS database (https://www.species360.org/). Data on mean plasma glucose concentration were the most often reported measure on glucose in ZIMS. We used the mean concentration of glucose rather than median concentration of glucose because there were no significant outliers in the mean concentration of glucose data across vertebrates, within mammals, and within birds according to Grubbs’ test^111,112^, and the mean concentration of glucose data within mammals and birds followed a normal distribution (according to Shapiro’s test^113^).

Data for neoplasia (malignant and benign tumors) and malignancy prevalence, including gastrointestinal neoplasia and malignancy prevalence as well as non-gastrointestinal neoplasia and malignancy prevalence, were collected from previous studies^13^. Each species’ neoplasia and malignancy prevalence is based on data from at least 20 individuals in that species. These data are from several zoological and veterinary institutions.

Body mass (grams) and gestation length (months) have been previously shown to significantly correlate with neoplasia prevalence and malignancy prevalence, respectively, across vertebrates^10^. Therefore, to control for the effect of body mass and gestation length on neoplasia prevalence and malignancy prevalence across vertebrates, we collected body mass and gestation length data for each species from published resources^10,114,115^.

## Statistical Analyses

We conducted all analyses in R version 4.0.5^116^. We transformed the adult weight, gestation length, and plasma glucose values, to normalize the distribution of data. We chose the most relevant transformation for each variable based on the “transformTukey” function (“rcompanion” R package). Specifically, in cases where the distribution of data was not normal (Shapiro’s test^113^), we transformed the weight values (–1 x weight ^ –0.125), the gestation length values to the power of 0.2, and the glucose concentration values to the power of 0.425. In the analyses between neoplasia and malignancy prevalence with plasma glucose concentration, we also centred the independent variable by subtracting it by the mean for normalization of the data. We used the R packages CAPER^117^, phytools^118^, geiger^119^, and tidyverse^120^, and then performed phylogenetic generalised least squares (PGLS) regressions in all analyses to control for the phylogenetic non-independence among species using the NCBI tree creator^122^. In analyses where one of the variables was malignancy prevalence or neoplasia prevalence, we weighted the analyses by 1/(square root of the number of necropsies per species) (from Revell^118^).

We classified diet type (carnivore, herbivore, and omnivore) and trophic level (Herbivore, Invertivore, Primary Carnivore, and Secondary Carnivore) as categorical variables. Next, we classified percentage of food type (endothermic vertebrates, ectothermic vertebrates, fruit, invertebrates, plants, seeds), and percentage of animal-based and plant-based foods, malignancy prevalence, neoplasia prevalence, and glucose concentration as numerical variables. We conducted univariate and multivariate analyses, with malignancy/neoplasia prevalence across tissues, gastrointestinal malignancy/neoplasia prevalence, or non-gastrointestinal malignancy/neoplasia prevalence as dependent variables, and mean glucose concentration, trophic level, diet type, percentage of animal and plant-based foods, food type, gestation length, and body weight as independent variables. We also tested whether the *P*-values passed the False Discovery Rate (FDR) correction in each of these six groups of analyses [1. correlations between diet and mean plasma glucose concentrations across vertebrates (10 tests); 2. correlations between cancer and mean plasma glucose concentrations across vertebrates (30 tests); 3. correlations between cancer and diet across vertebrates (108 tests); 4. correlations between cancer and mean plasma glucose concentrations within mammals (24 tests); 5. correlations between cancer and mean plasma glucose concentrations within birds (24 tests); 6. correlations between cancer and mean plasma glucose concentrations within reptiles (24 tests)].

## Supplementary Data

The data used in this study will be made available upon acceptance of the manuscript for publication.

## Supplementary Figure Legends

**Supplementary Figure 1. Relationship between malignancy prevalence or neoplasia prevalence and mean plasma glucose concentration.** (**A**) Mean plasma glucose concentration is not significantly correlated with overall neoplasia prevalence for 160 species (PGLS: *P*-value = 0.13). **(B)** Mean plasma glucose concentration is negatively correlated with gastrointestinal neoplasia prevalence for 108 species (PGLS: *P*-value = 0.007) even after correcting for multiple testing (Table 1B). (**C**) Mean plasma glucose concentration is not significantly correlated with non-gastrointestinal malignancy prevalence for 146 species (PGLS: *P*-value = 0.53), nor (**D**) non-gastrointestinal neoplasia prevalence for 146 species (PGLS: *P*-value = 0.18). Each dot represents the neoplasia prevalence across tissues (A), the gastrointestinal neoplasia prevalence (B), the non-gastrointestinal malignancy prevalence (C), the non-gastrointestinal neoplasia prevalence (D), and the average plasma glucose concentration of one species; Amphibia: Dark Blue; Aves: Blue; Elasmobranchii: Green; Mammalia: Orange; Reptilia: Red. We show images of significant outlier species (Rosner’s test). Animal silhouettes from PhyloPic (http://www.phylopic.org/).

**Supplementary Figure 2**. **No significant correlation between cancer or neoplasia prevalence and the percentage of food type in a species diet after correcting for multiple testing.** (**Α**) The percentage of plants in a species’ diet is negatively correlated with neoplasia prevalence across tissues for 39 species (PGLS: *P*-value = 0.006), but not when controlling for the variance in their plasma glucose concentrations (PGLS: *P*-value > 0.05) or after correcting for multiple testing (Table 1F). There is no significant correlation between the percentage of fruit, invertebrates, seeds, endothermic vertebrates (Vendotherms), or ectothermic vertebrates (Vectotherms) in a species’ diet and neoplasia prevalence across tissues for 25, 35, 18, 15 or 11 species, respectively (PGLS: *P*-value > 0.05). (**B**) The percentage of plants in a species’ diet is negatively correlated with gastrointestinal neoplasia prevalence for 22 species (PGLS: *P*-value < 0.05) after correcting for multiple testing, but not when controlling for the variance in their plasma glucose concentrations (Table 1F). The percentage of endothermic vertebrates (Vendotherms) in a species’ diet is negatively correlated with gastrointestinal neoplasia prevalence for 13 species (PGLS: *P*-value < 0.05), but not after correcting for multiple testing (Table 1F). There is no significant correlation between the percentage of fruit and invertebrates in a species’ diet and gastrointestinal neoplasia prevalence for 19 and 23 species, respectively (PGLS: *P*-value > 0.05). (**C**) There is no significant correlation between the percentage of fruit, plants, invertebrates, seeds, or endothermic vertebrates (Vendotherms) in a species’ diet and non-gastrointestinal malignancy prevalence for 22, 36, 31, 16, or 13 species, respectively (PGLS: *P*-value > 0.05). (**D**) There is no significant correlation between the percentage of fruit, plants, invertebrates, seeds, or endothermic vertebrates (Vendotherms) in a species’ diet and non-gastrointestinal neoplasia prevalence for 22, 36, 31, 16, or 13 species, respectively (PGLS: *P*-value > 0.05). Each dot shows the neoplasia prevalence across tissues (A), the gastrointestinal neoplasia prevalence (B), the non-gastrointestinal malignancy prevalence (C), the non-gastrointestinal neoplasia prevalence (D), and the percentage of food type in the diet of one species. We added minimal jitter in the plots in order to better visualize individual data points.

**Supplementary Figure 3. No significant correlation between malignancy prevalence or neoplasia prevalence and percentage of plant-based or meat-based food in species’ diet after correcting for multiple testing.** (**A**) The percentage of plant-based food in a species’ diet is negatively correlated with neoplasia prevalence across tissues for 50 species without controlling for the variance in their plasma glucose concentrations (PGLS: *P*-value = 0.01), but not after correcting for multiple testing (Table 1F). There is no significant correlation between the percentage of animal-based food in a species’ diet and neoplasia prevalence across tissues for 44 species (PGLS: *P*-value > 0.05). (**B**) The percentage of plant-based food in a species’ diet is negatively correlated with gastrointestinal neoplasia prevalence for 29 species with (PGLS: *P*-value = 0.02) or without (PGLS: *P*-value = 0.01) controlling for the variance in their plasma glucose concentrations, but not after applying corrections for multiple testing (Table 1F). There is no significant correlation between the percentage of animal-based food in a species’ diet and gastrointestinal neoplasia prevalence for 32 species (PGLS: *P*-value > 0.05). (**C**) The percentage of plant-based or animal-based food in a species’ diet is not significantly correlated with non-gastrointestinal malignancy prevalence for 46 or 39 species, respectively (PGLS: *P*-value > 0.05). (**D**) The percentage of plant-based or animal-based food in a species’ diet is not significantly correlated with non-gastrointestinal neoplasia prevalence for 46 or 39 species after correcting for multiple testing. Each dot shows the neoplasia prevalence across tissues (A), the gastrointestinal neoplasia prevalence (B), the non-gastrointestinal malignancy prevalence (C), the non-gastrointestinal neoplasia prevalence (D), and the percentage of plant-based or meta-based food in the diet of one species. We added minimal jitter in the plots in order to better visualize individual data points.

**Supplementary Figure 4**. **No significant correlation between malignancy prevalence or neoplasia prevalence and diet type.** Diet type is not significantly correlated with neoplasia across tissues (**A**), gastrointestinal neoplasia prevalence (**B**), non-gastrointestinal malignancy prevalence (**C**), or non-gastrointestinal neoplasia prevalence (**D**) (PGLS: *P*-value > 0.05). Each dot shows the neoplasia prevalence across tissues (A), the gastrointestinal neoplasia prevalence (B), the non-gastrointestinal malignancy prevalence (C), the non-gastrointestinal neoplasia prevalence (D), and the diet type of one species. N shows the number of species per diet category. The horizontal black line in each diet category shows the median neoplasia prevalence across tissues (A), the median gastrointestinal neoplasia prevalence (B), the median non-gastrointestinal malignancy prevalence (C), or the median non-gastrointestinal neoplasia prevalence (D), in that diet category.

**Supplementary Figure 5**. **No significant correlation between cancer or neoplasia prevalence and mean plasma glucose concentration in mammals.** There is no significant correlation between mean plasma glucose concentration and (**A**) malignancy prevalence across tissues for 68 mammalian species, (**B**) neoplasia prevalence across tissues for 68 mammalian species, (**C**) gastrointestinal malignancy prevalence for 48 mammalian species, (**D**) gastrointestinal neoplasia prevalence for 48 mammalian species, (**E**) non-gastrointestinal malignancy prevalence for 67 mammalian species, and (**F**) non-gastrointestinal neoplasia prevalence for 67 mammalian species (PGLS: *P*-value > 0.05). Each dot shows the malignancy prevalence across tissues (A), the neoplasia prevalence across tissues (B), the gastrointestinal malignancy prevalence (C), the gastrointestinal neoplasia prevalence (D), the non-gastrointestinal malignancy prevalence (E), the non-gastrointestinal neoplasia prevalence (F), and the average plasma glucose concentration of one species. We show images of significant outlier species (Rosner’s test). Animal silhouettes from PhyloPic (http://www.phylopic.org/).

**Supplementary Figure 6**. **No significant correlation between cancer or neoplasia prevalence and mean plasma glucose concentration in birds.** There is no significant correlation between mean plasma glucose concentration and (**A**) malignancy prevalence across tissues for 51 bird species, (**B**) neoplasia prevalence across tissues for 51 bird species, (**C**) gastrointestinal malignancy prevalence for 31 bird species, (**D**) gastrointestinal neoplasia prevalence for 31 bird species, (**E**) non-gastrointestinal malignancy prevalence for 41 bird species, and (**F**) non-gastrointestinal neoplasia prevalence for 41 bird species (PGLS: *P*-value > 0.05). Each dot shows the malignancy prevalence across tissues (A), the neoplasia prevalence across tissues (B), the gastrointestinal malignancy prevalence (C), the gastrointestinal neoplasia prevalence (D), the non-gastrointestinal malignancy prevalence (E), the non-gastrointestinal neoplasia prevalence (F), and the average plasma glucose concentration of one species. We show images of significant outlier species (Rosner’s test). Animal silhouettes from PhyloPic (http://www.phylopic.org/).

**Supplementary Figure 7**. **No significant correlation between cancer or neoplasia prevalence and mean plasma glucose concentration in reptiles.** There is no significant correlation between mean plasma glucose concentration and (**A**) malignancy prevalence across tissues for 35 reptile species, (**B**) neoplasia prevalence across tissues for 35 reptile species, (**C**) gastrointestinal malignancy prevalence for 25 reptile species, (**D**) gastrointestinal neoplasia prevalence for 25 reptile species, (**E**) non-gastrointestinal malignancy prevalence for 32 reptile species, and (**F**) non-gastrointestinal neoplasia prevalence for 32 reptile species (PGLS: *P*-value > 0.05). Each dot shows the malignancy prevalence across tissues (A), the neoplasia prevalence across tissues (B), the gastrointestinal malignancy prevalence (C), the gastrointestinal neoplasia prevalence (D), the non-gastrointestinal malignancy prevalence (E), the non-gastrointestinal neoplasia prevalence (F), and the average plasma glucose concentration of one species. We show images of significant outlier species (Rosner’s test). Animal silhouettes from PhyloPic (http://www.phylopic.org/).

## Acknowledgements

We acknowledge the following institutions: Akron Zoo, Atlanta Zoo, Audubon Nature Institute, Bergen County Zoo, Birmingham Zoo, Buffalo Zoo, Capron Park Zoo, Central Florida Zoo, Dallas Zoo, El Paso Zoo, Elmwood Park Zoo, Fort Worth Zoo, Gladys Porter Zoo, Greensboro Science Center, Henry Doorly Zoo, Utah’s Hogle Zoo, Jacksonville Zoo, John Ball Zoo, Los Angeles Zoo, Louisville Zoo, Mesker Park Zoo, Miami Zoo, Oakland Zoo, Oklahoma City Zoo, Philadelphia Zoo, Phoenix Zoo, Pueblo Zoo, San Antonio Zoo, Santa Ana Zoo, Santa Barbara Zoo, Sedgwick County Zoo, Seneca Park Zoo, The Brevard Zoo, The Detroit Zoo, The Oregon Zoo, and Toledo Zoo. Thanks to Diego Mallo, Ping-Han Huang, and Walker Mellon for help with the statistical analyses. This work was supported in part by NIH grants U54 CA217376, U2C CA233254, P01 CA91955, and R01 CA140657 as well as CDMRP Breast Cancer Research Program Award BC132057 and the Arizona Biomedical Research Commission grant ADHS18-198847. The findings, opinions and recommendations expressed here are those of the authors and not necessarily those of the universities where the research was performed or the National Institutes of Health.

## Author Contributions

A.J.B. and S.E.K. designed the study, analysed the cancer data, the diet type data, and wrote the first draft. A.J.B. obtained additional adult weight data via AnAge. S.E.K. conceived the idea to compare glucose concentration data with cancer prevalence data, created the figures in R, collected and analysed the ZIMS glucose data, and performed the statistical analyses. Z.T.C. provided help with the phylogenetic analyses. S.M.R., Z.T.C., E.G.D., T.M.H., and A.M.B. helped in the collection of malignancy prevalence and neoplasia prevalence, adult weight, gestation length, and necropsies for each species. K.L.S. and C.C.M. provided guidance during the project. All authors commented on the final versions of the manuscript.

