## Supplementary figures and images for "The relationship between diet, plasma glucose, and cancer prevalence across vertebrates"

### Supplementary Figure 1

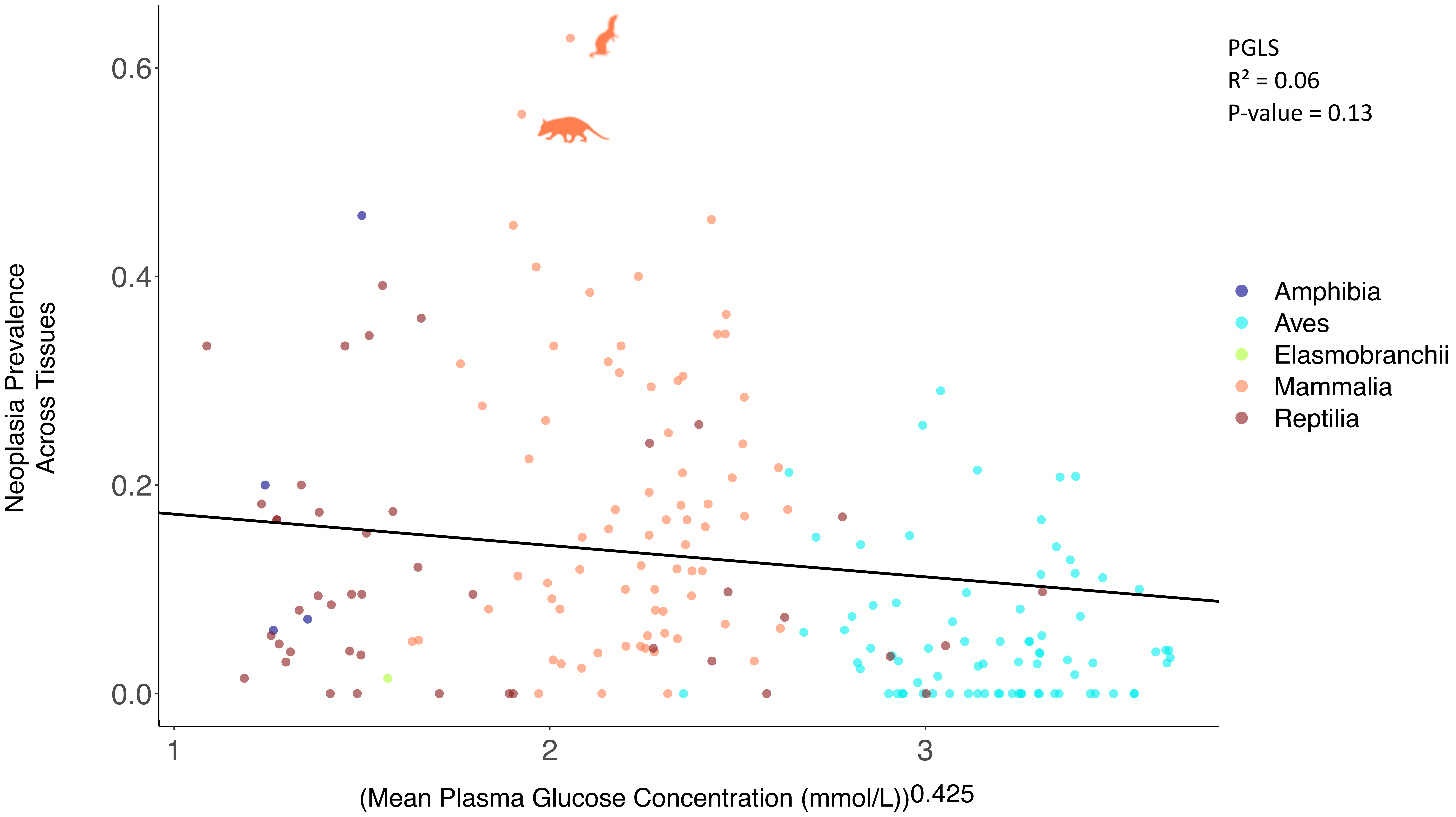

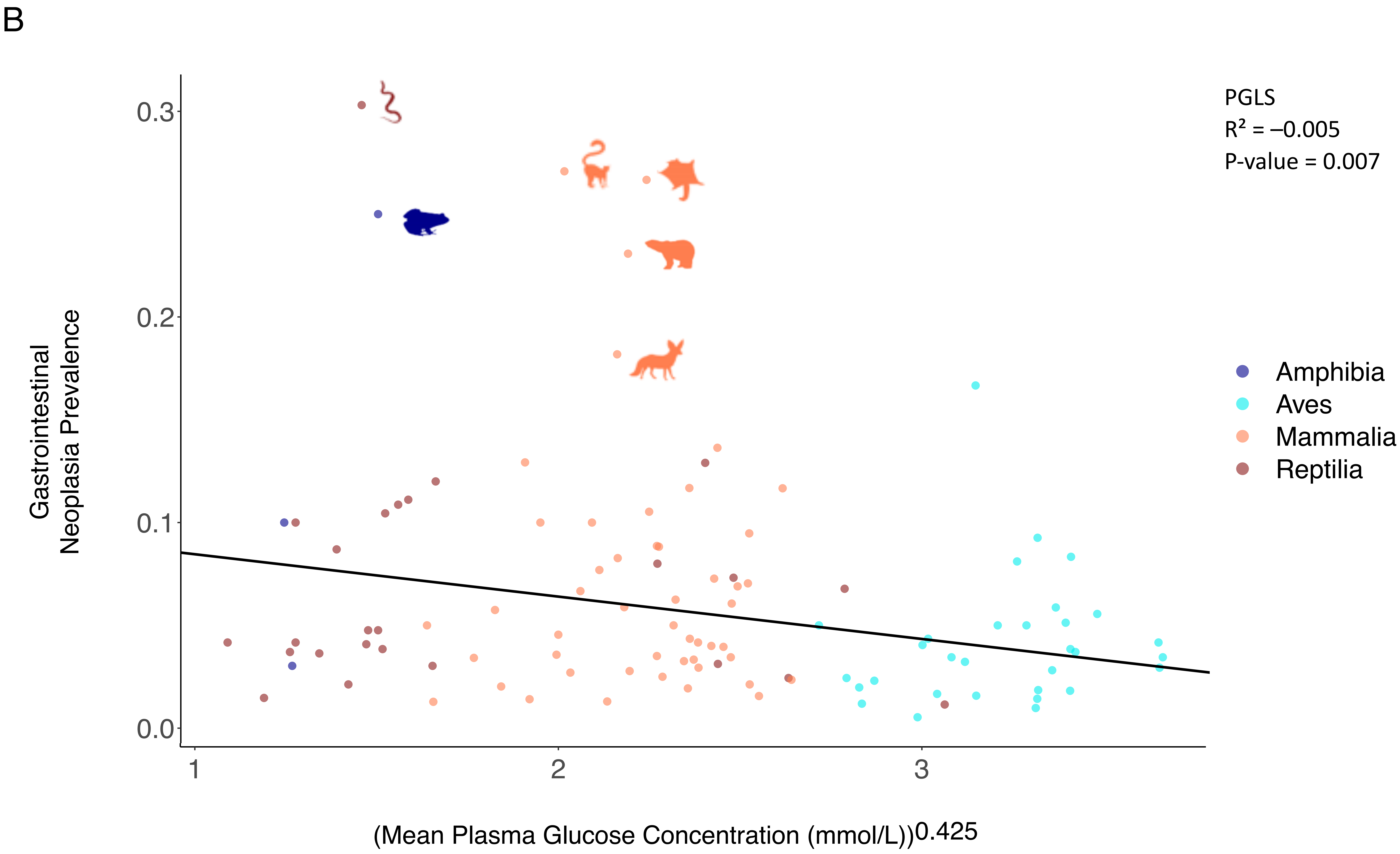

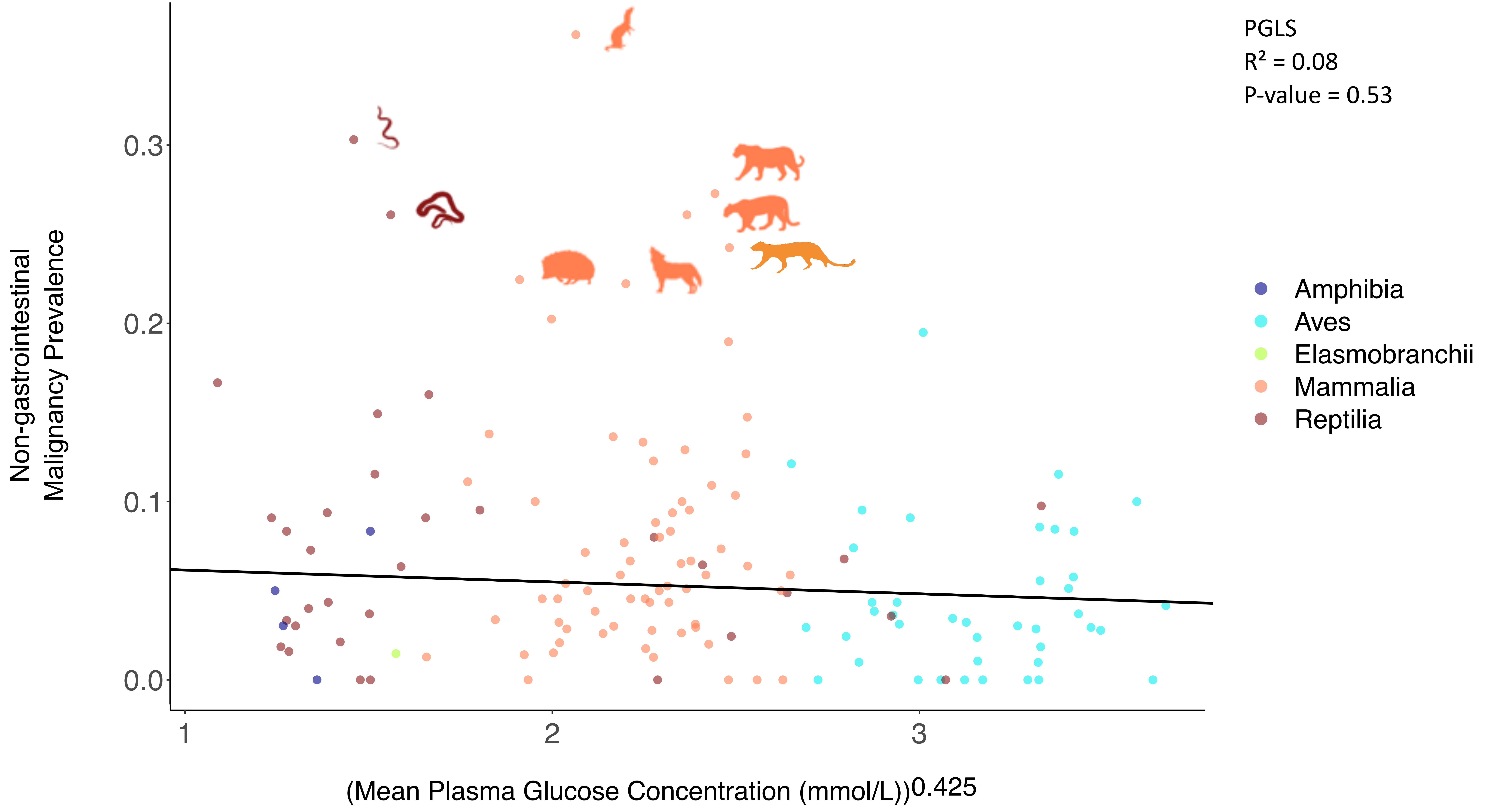

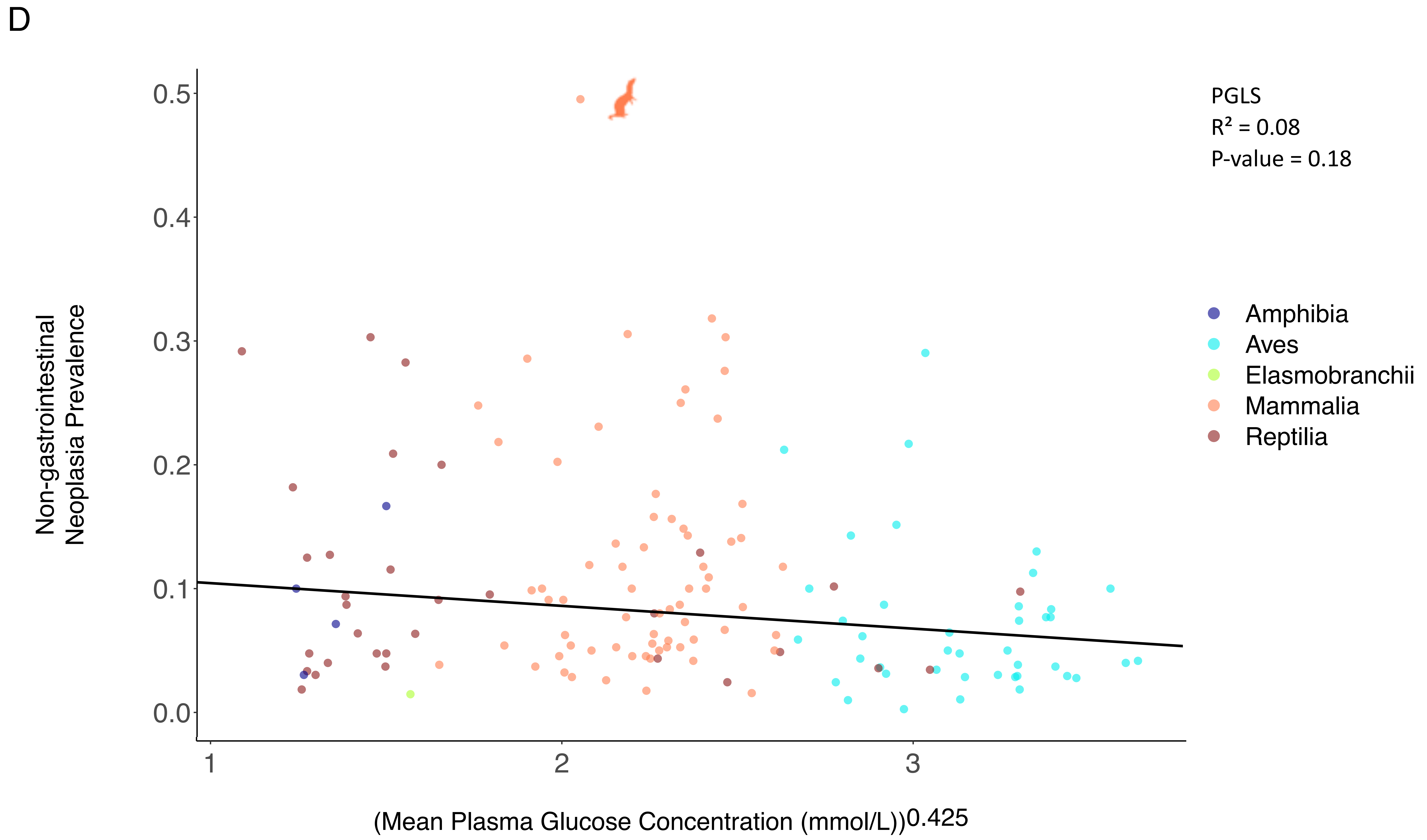

### Supplementary Figure 3

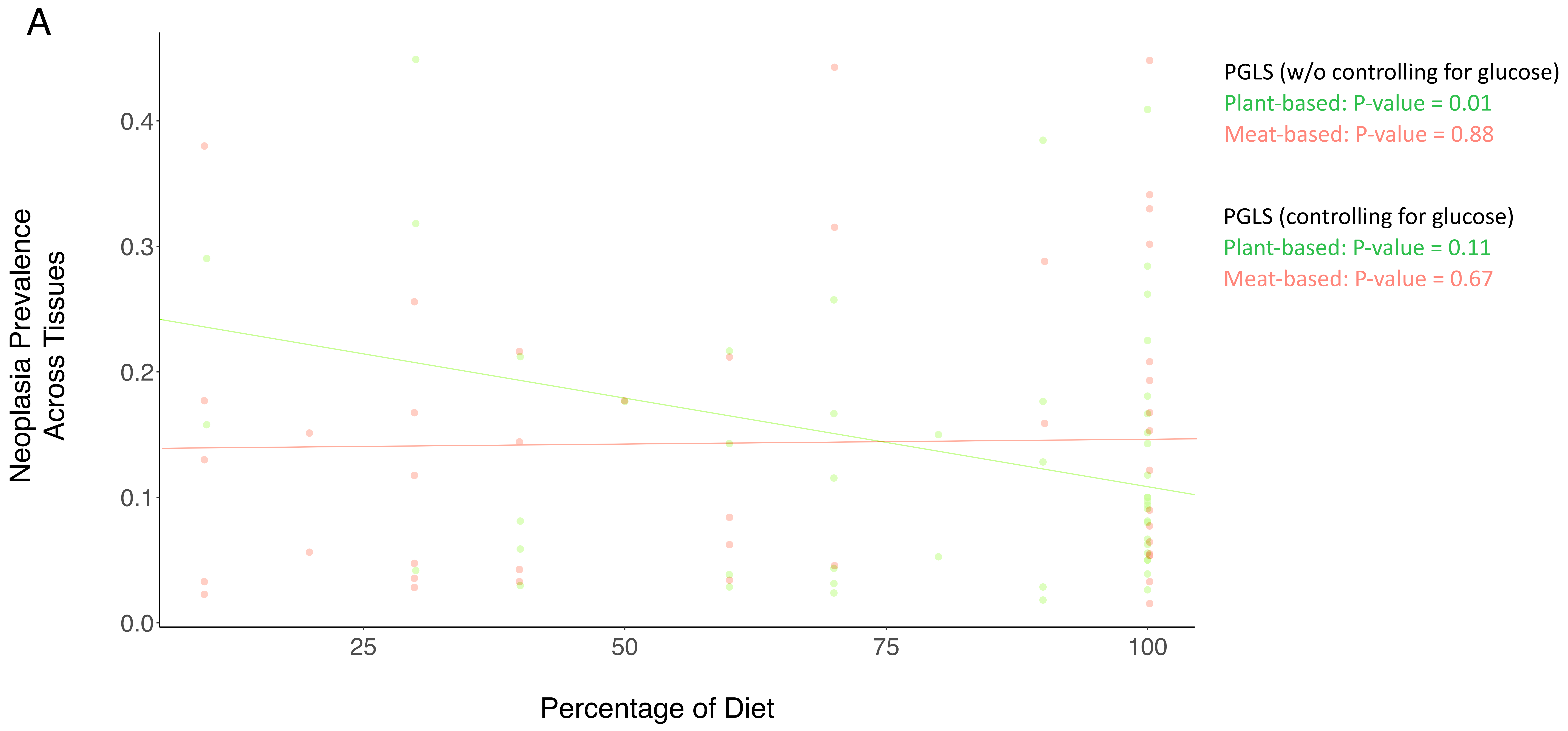

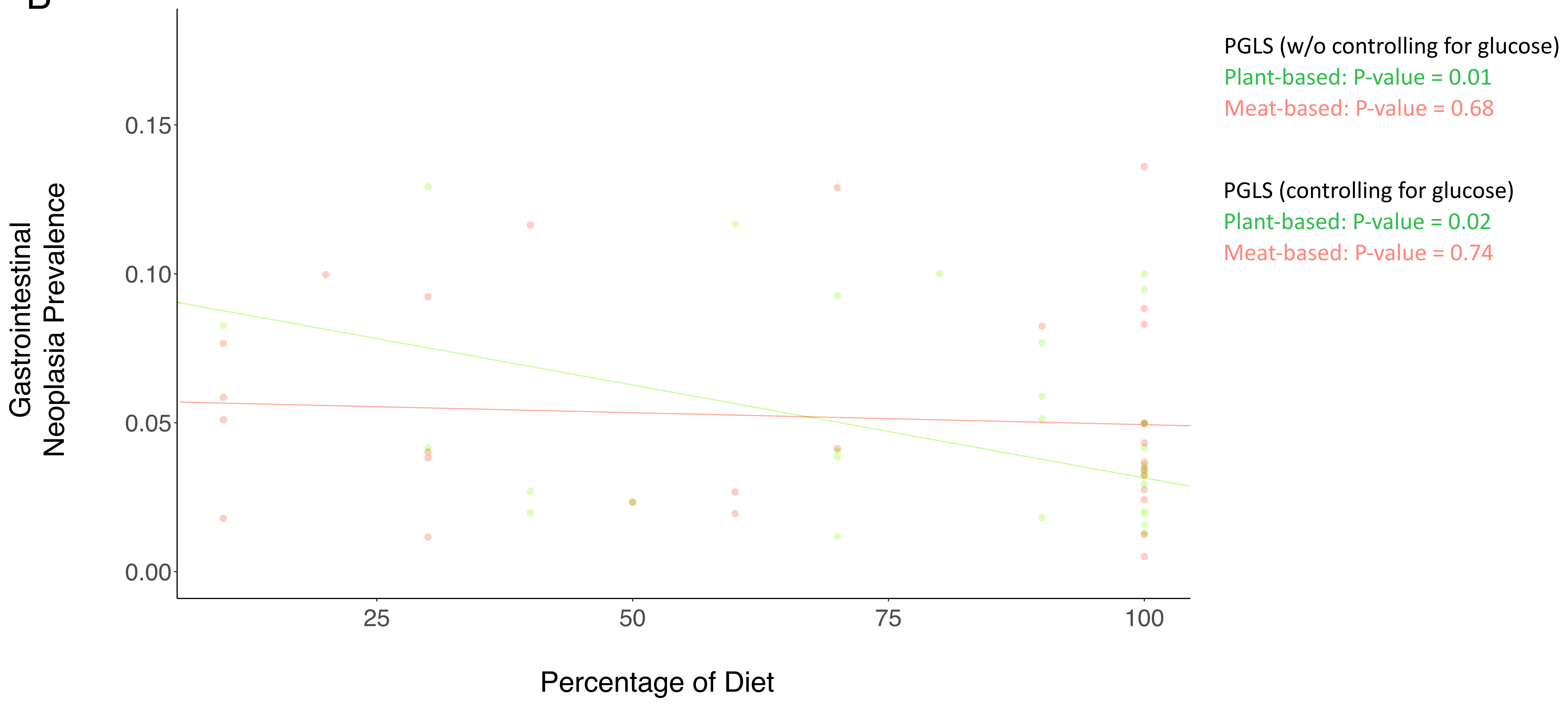

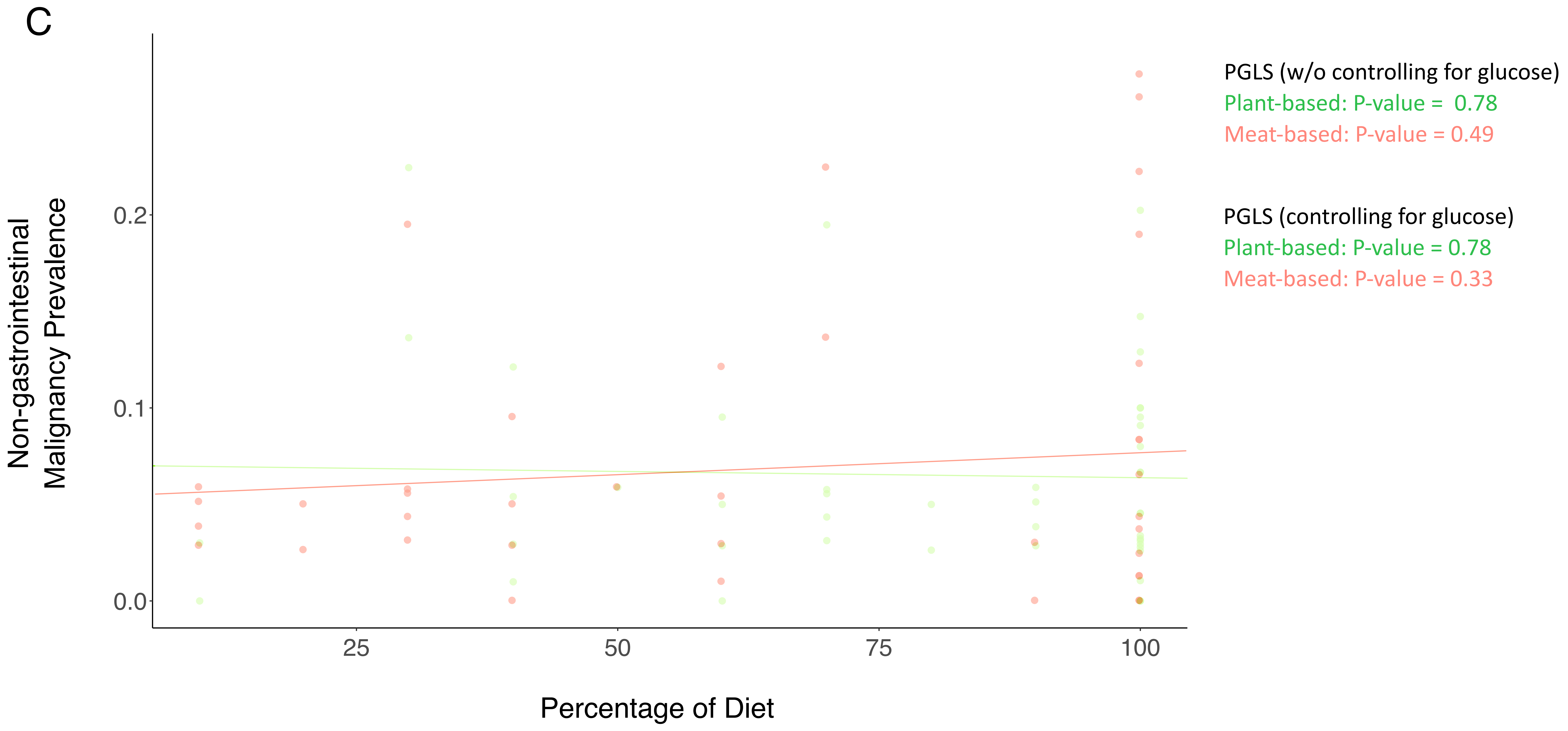

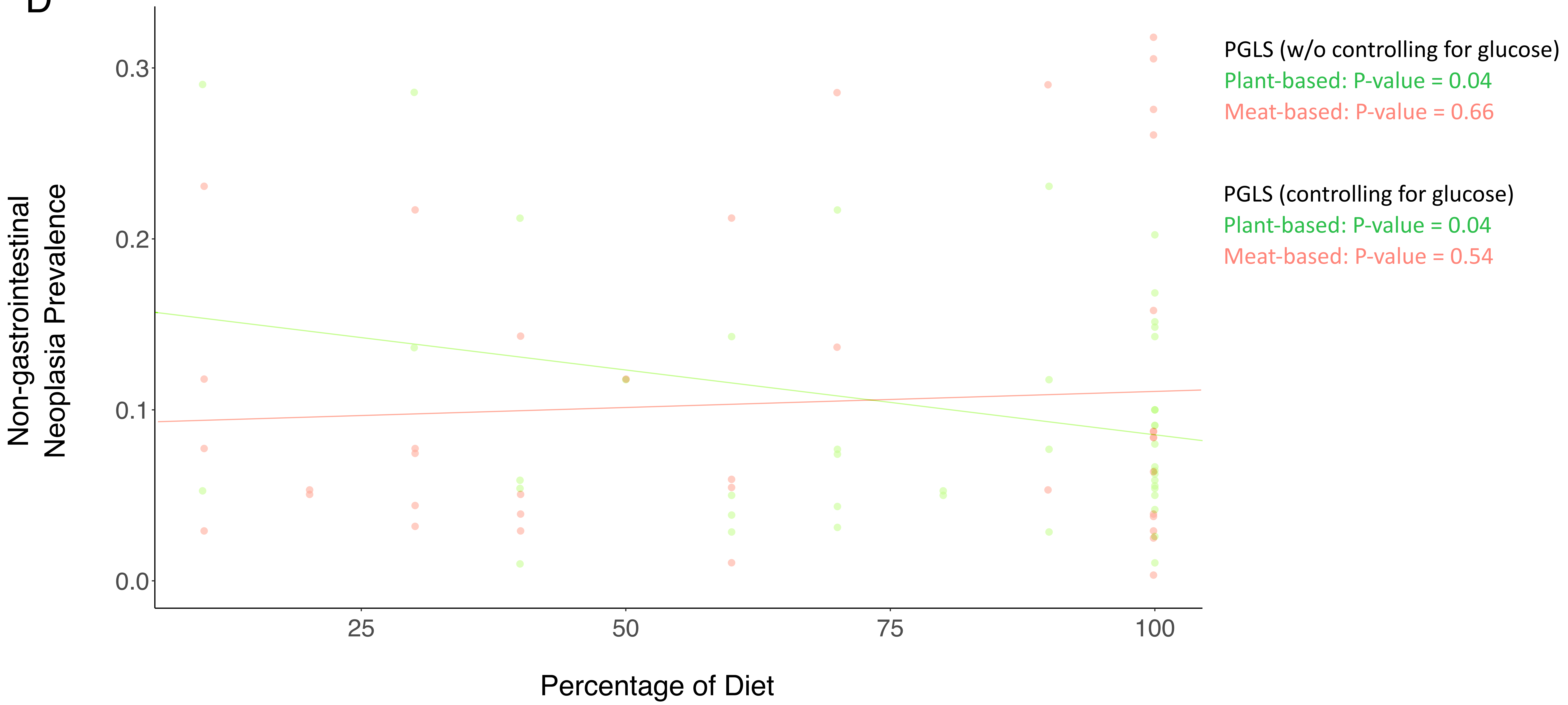

### Supplementary Figure 4

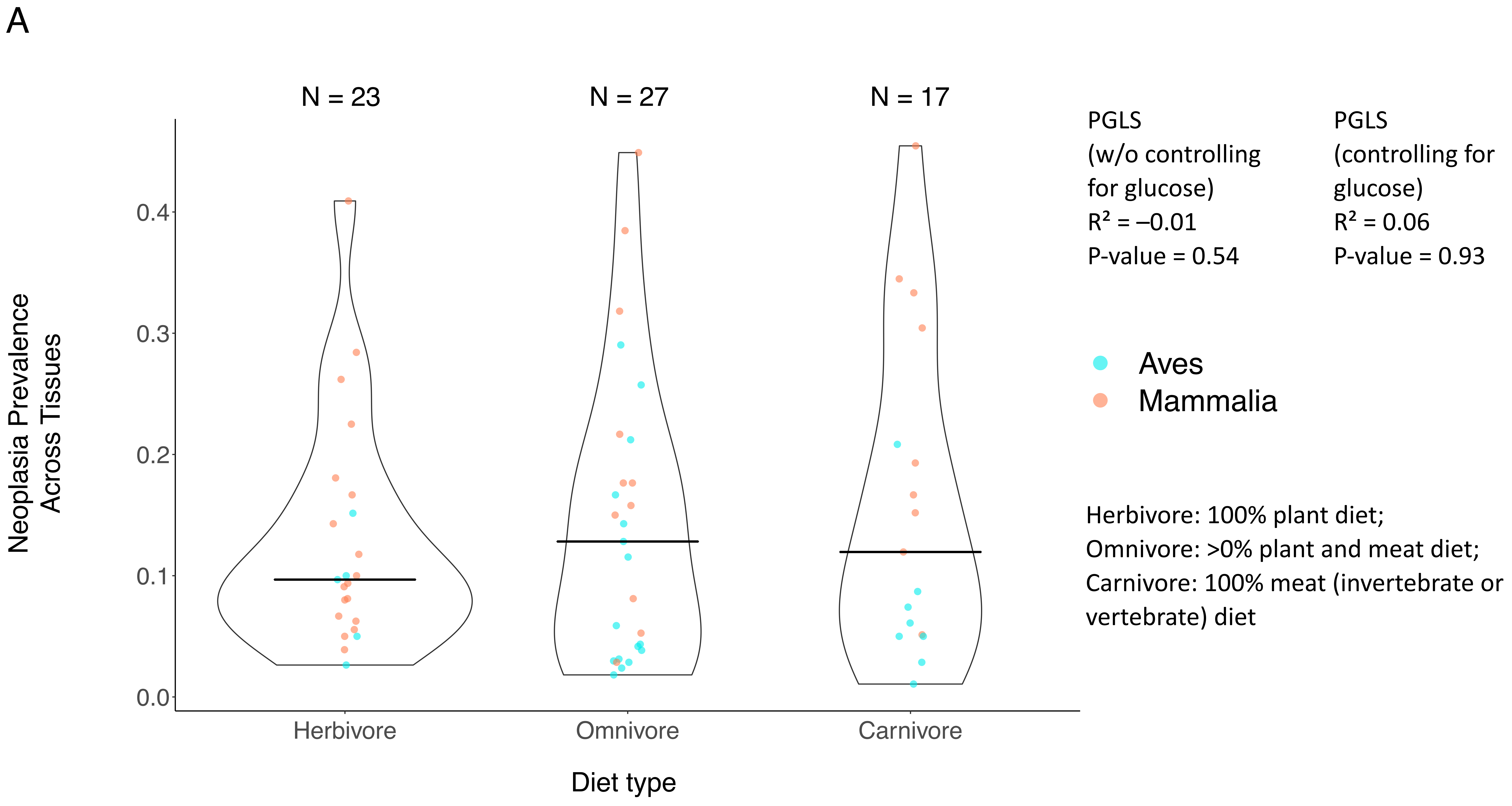

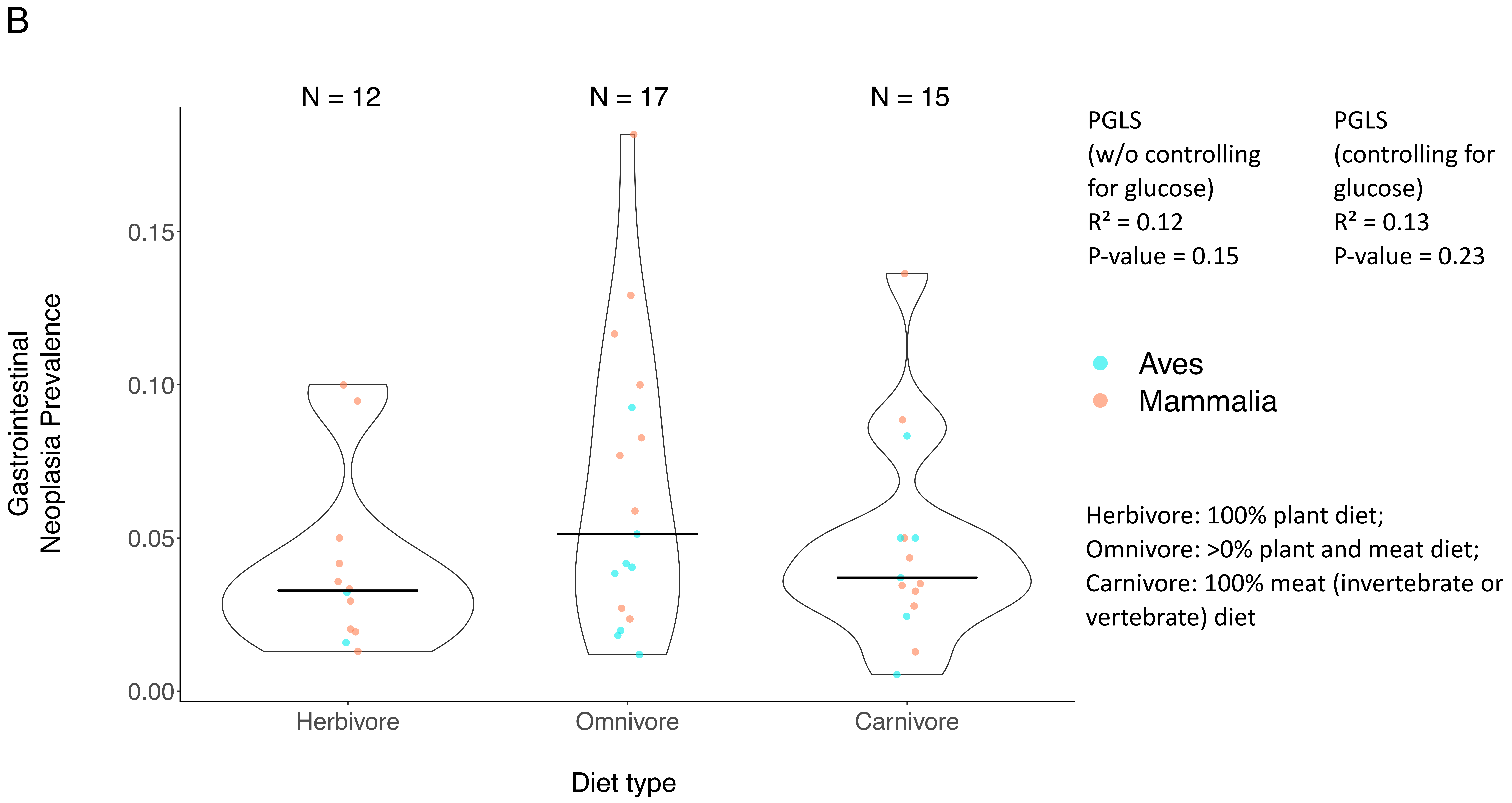

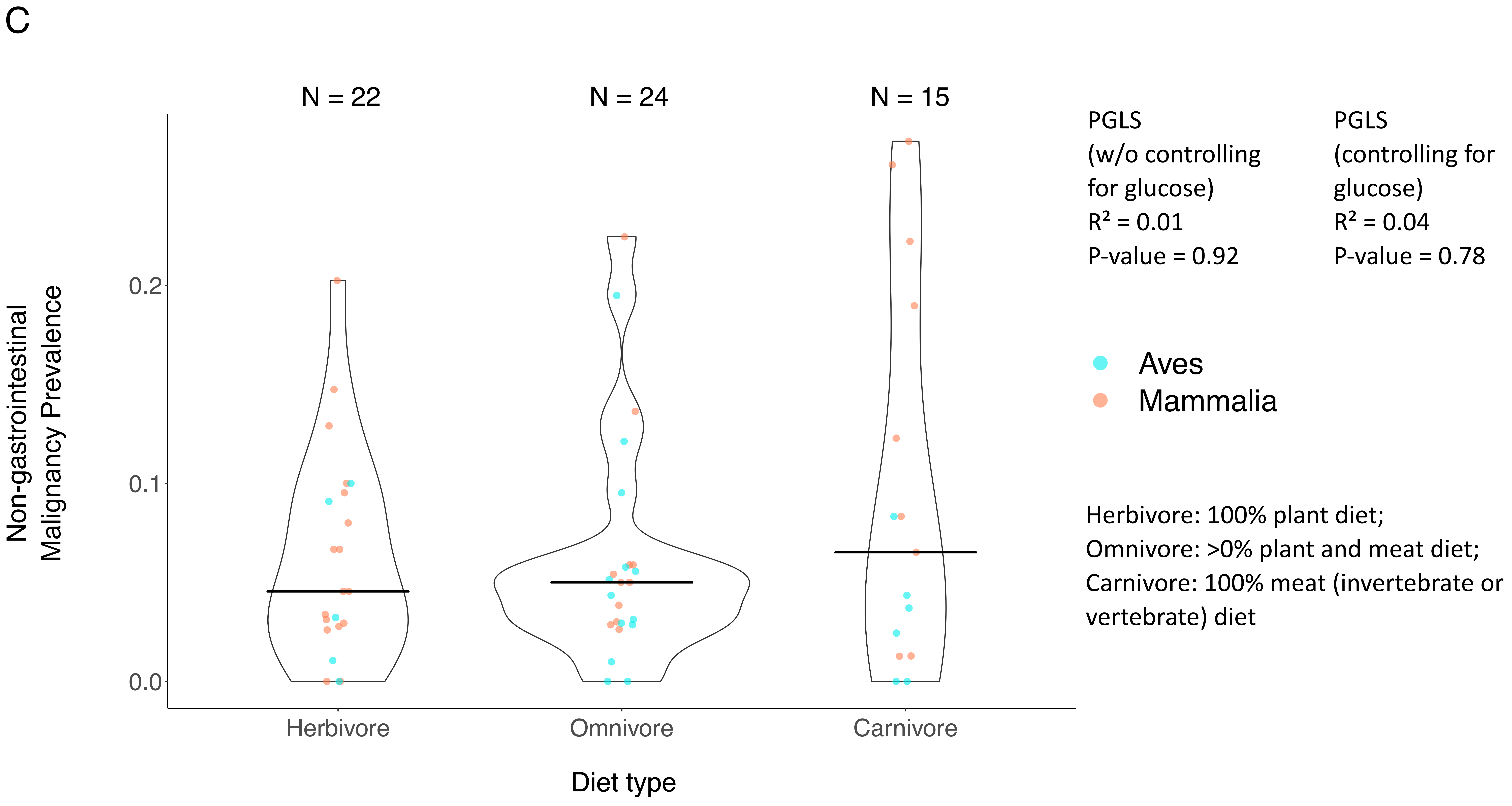

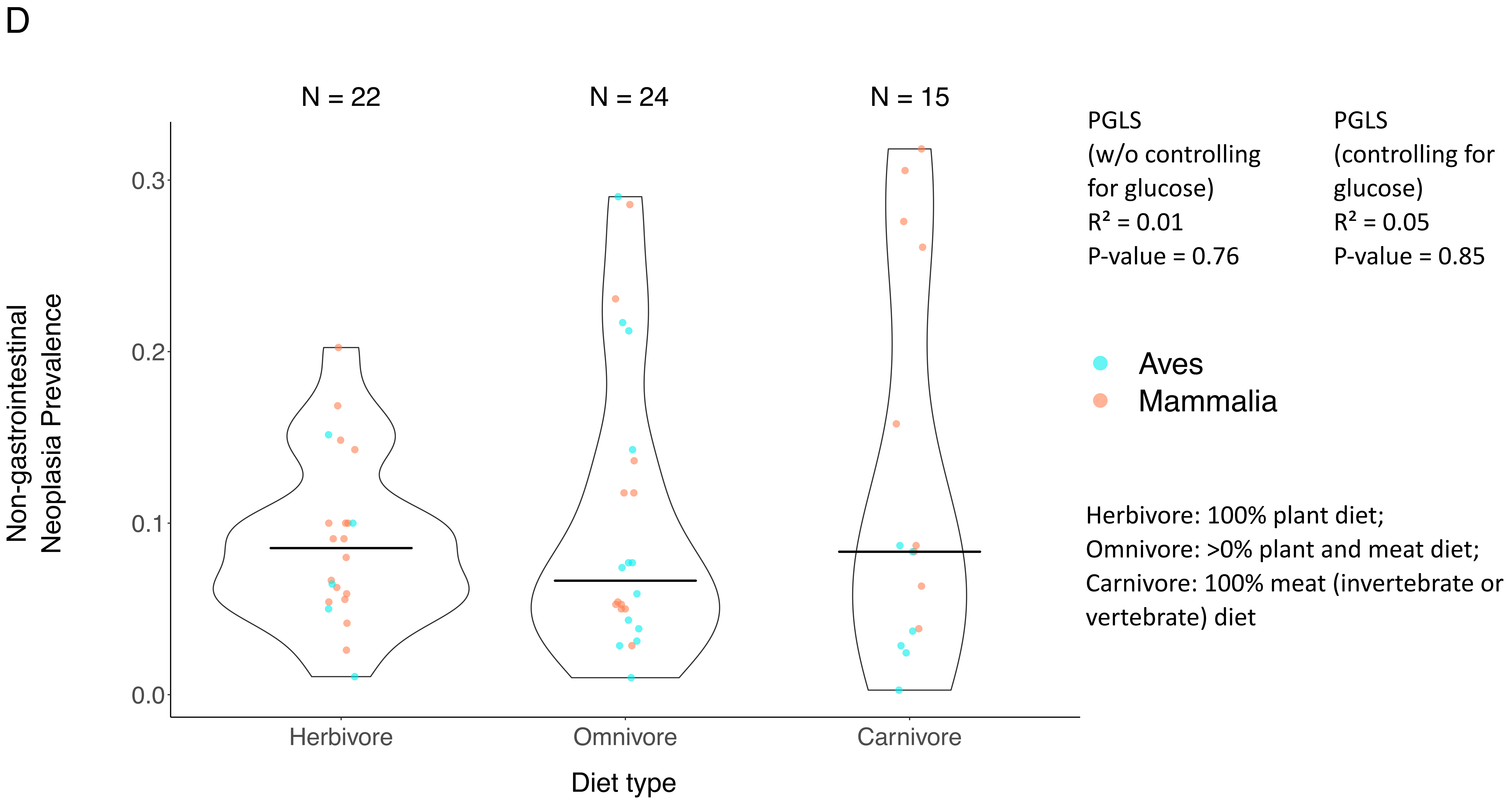

### Supplementary Figure 5

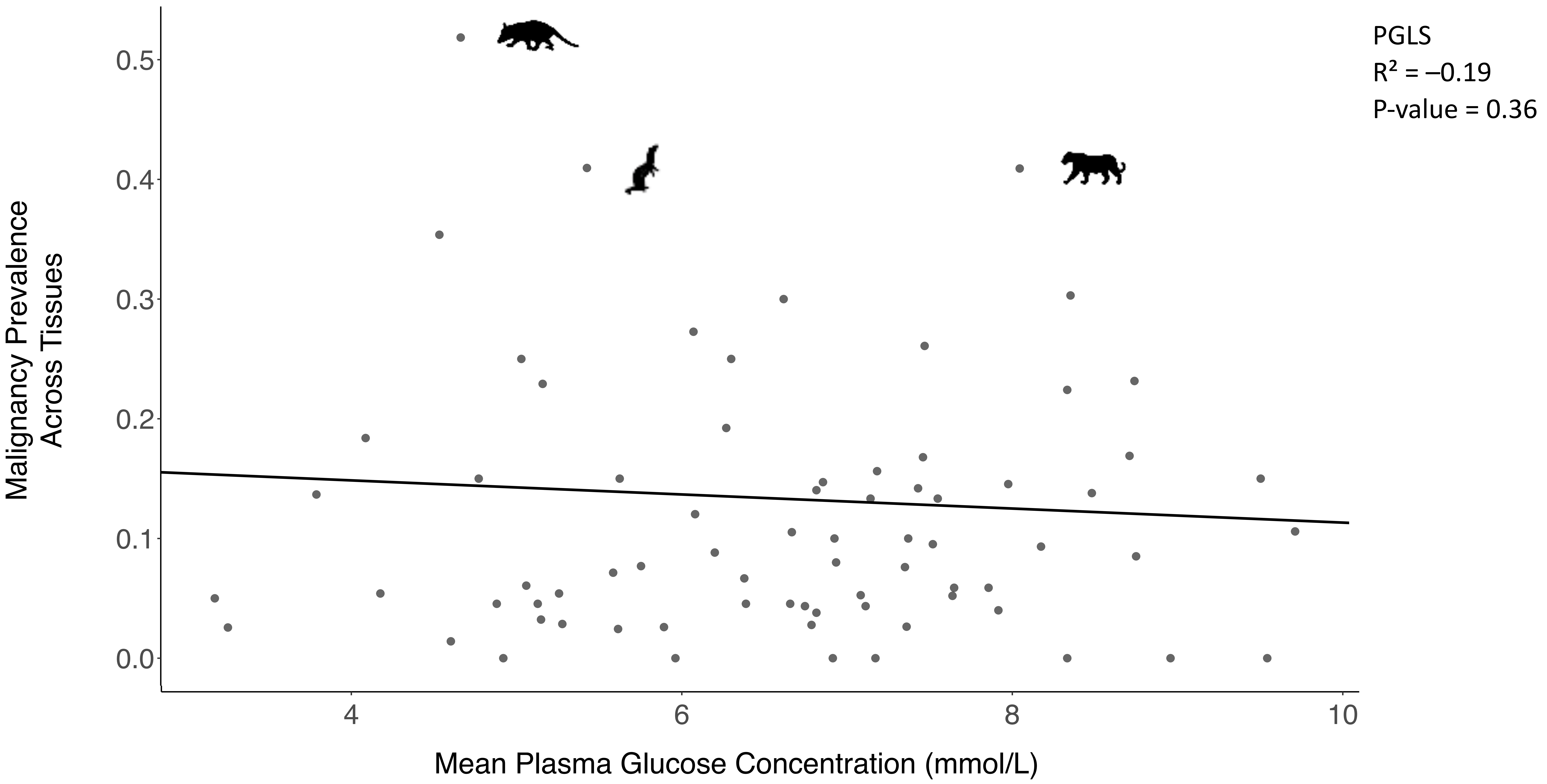

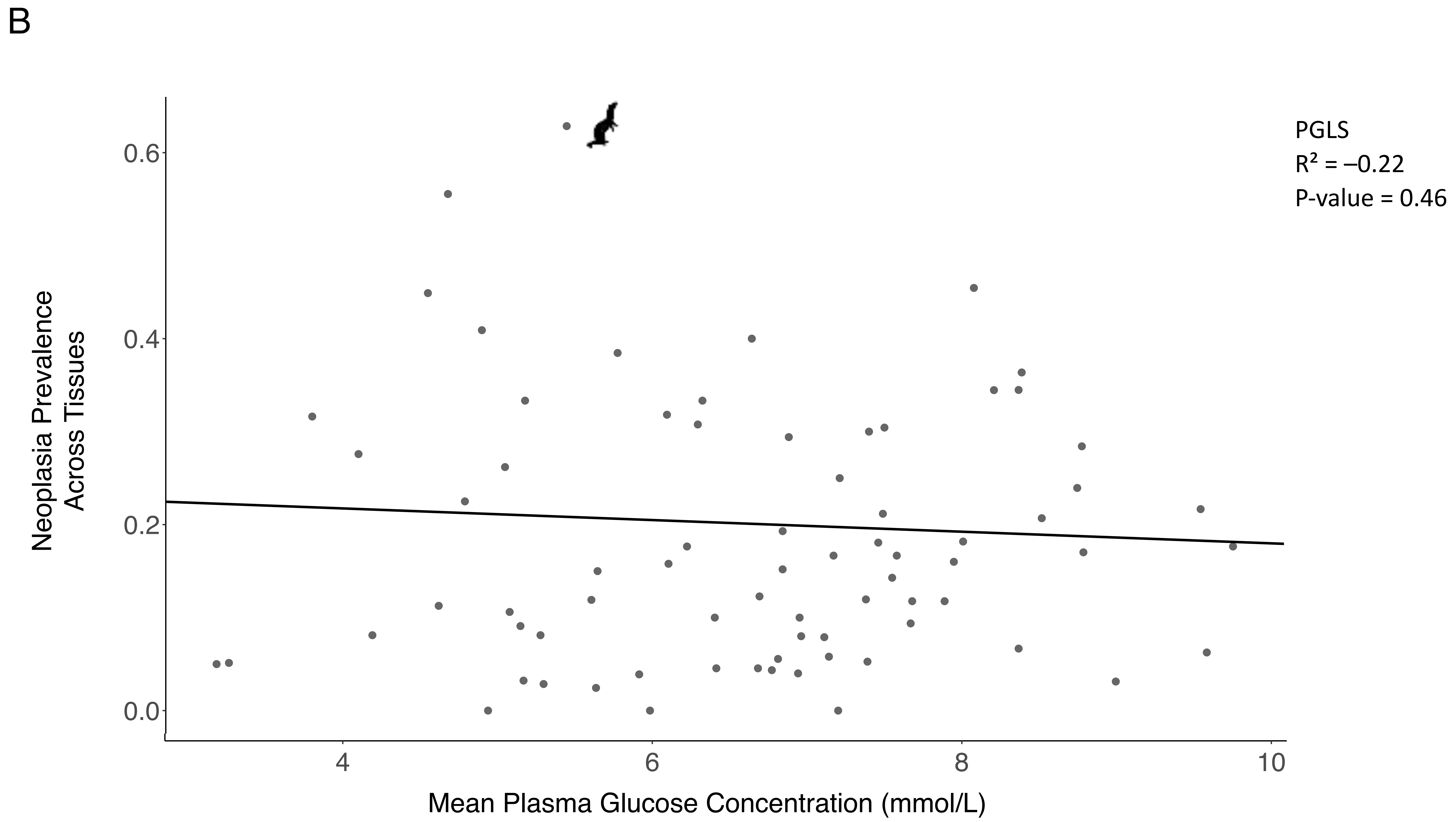

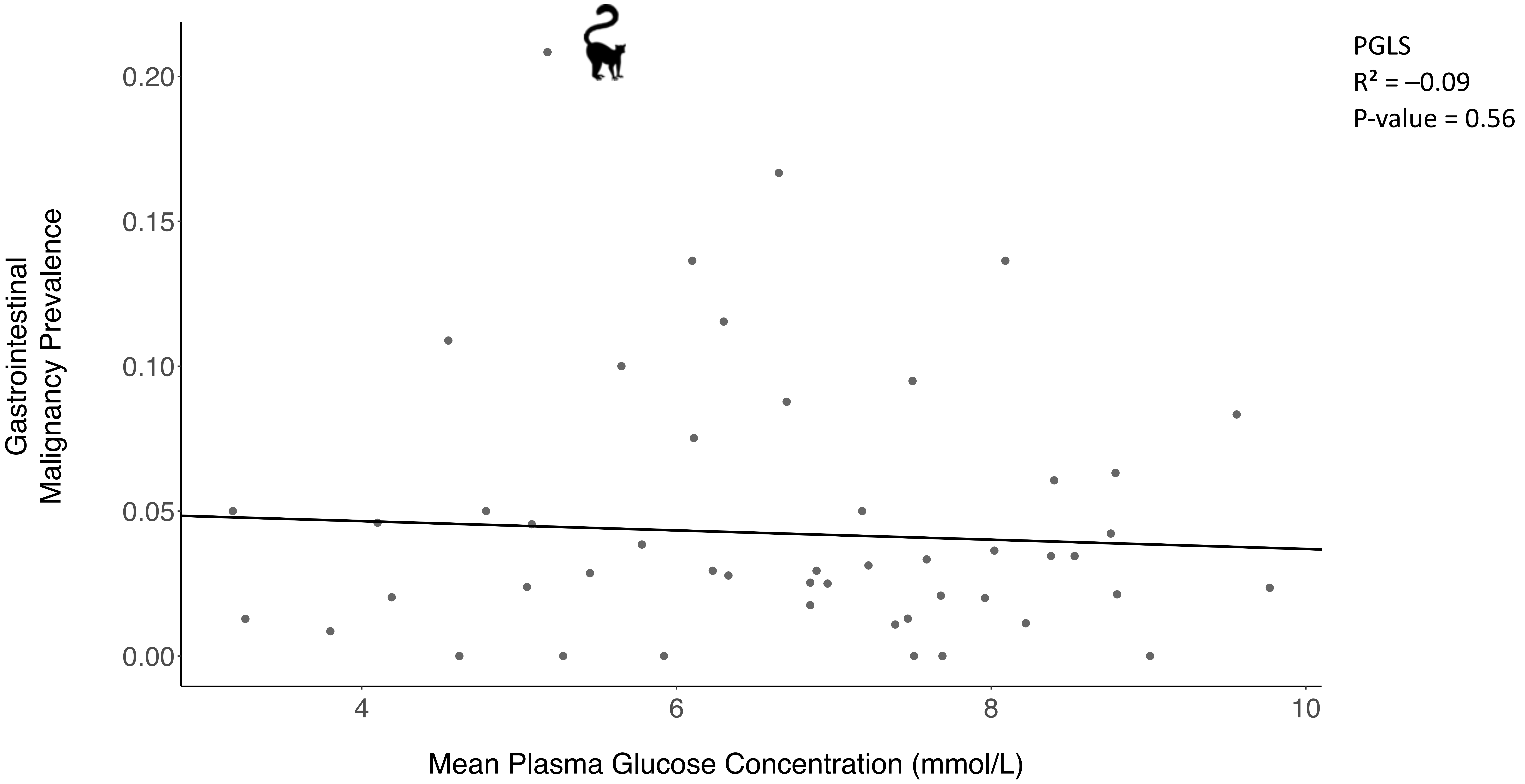

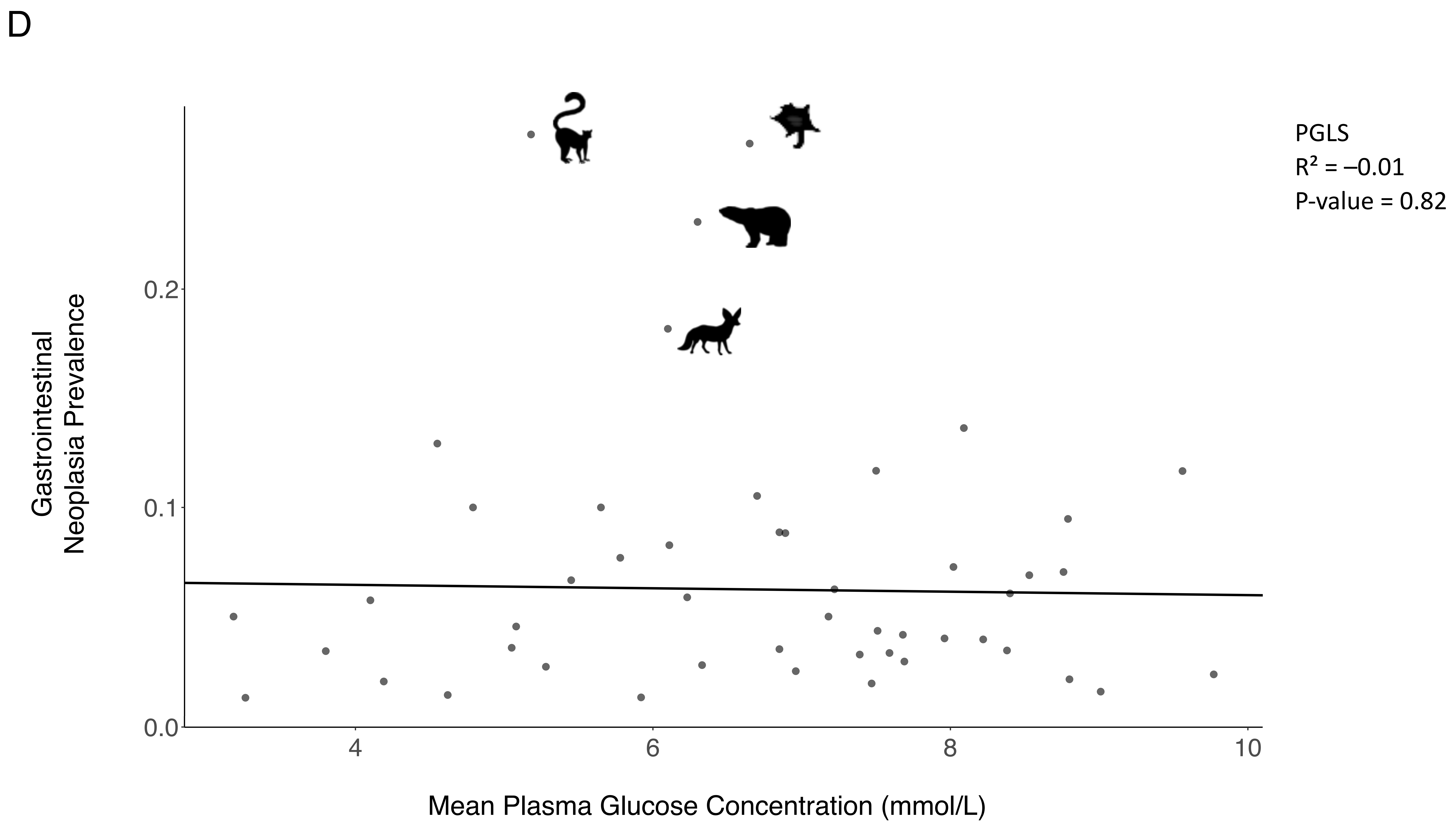

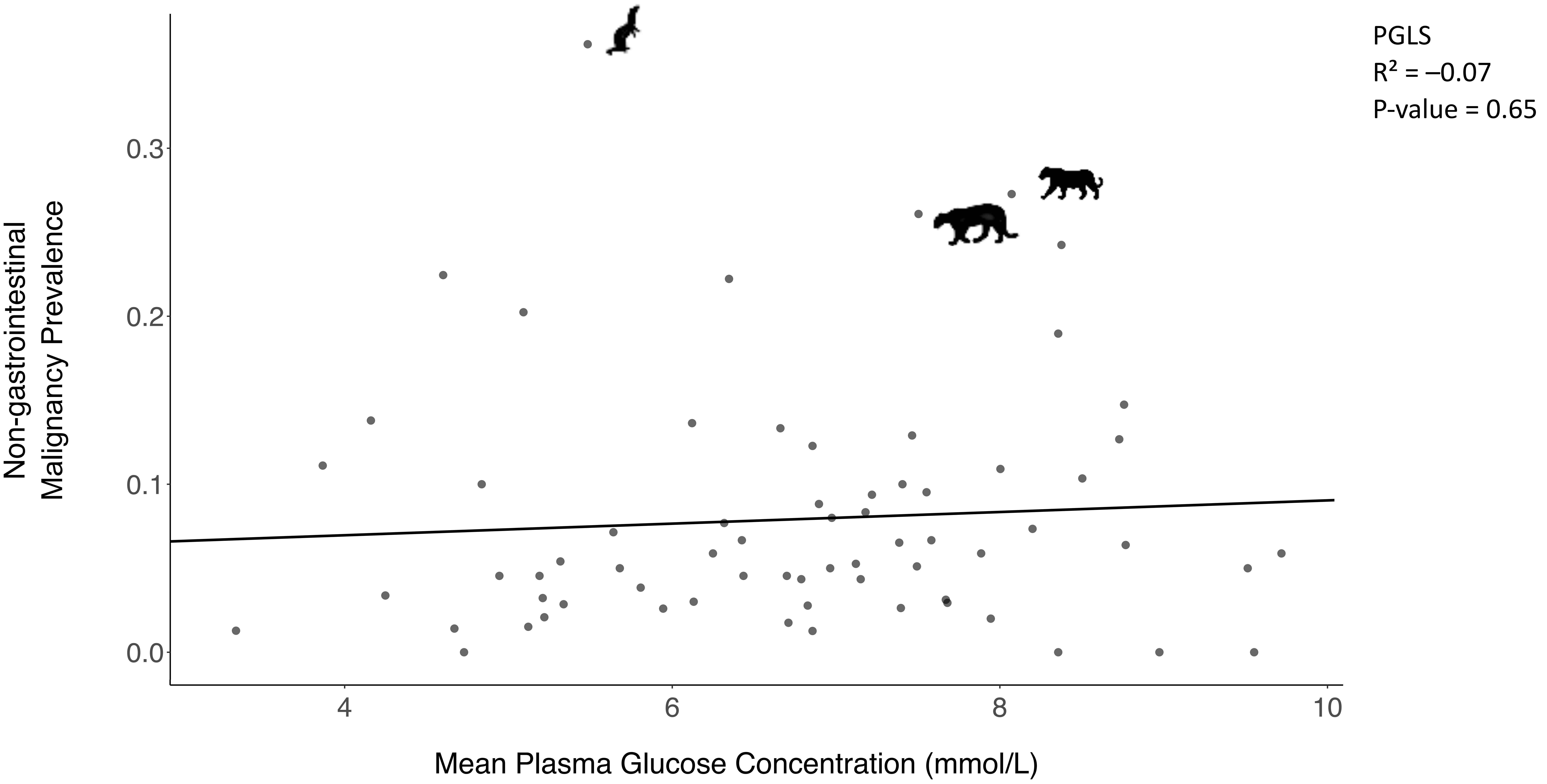

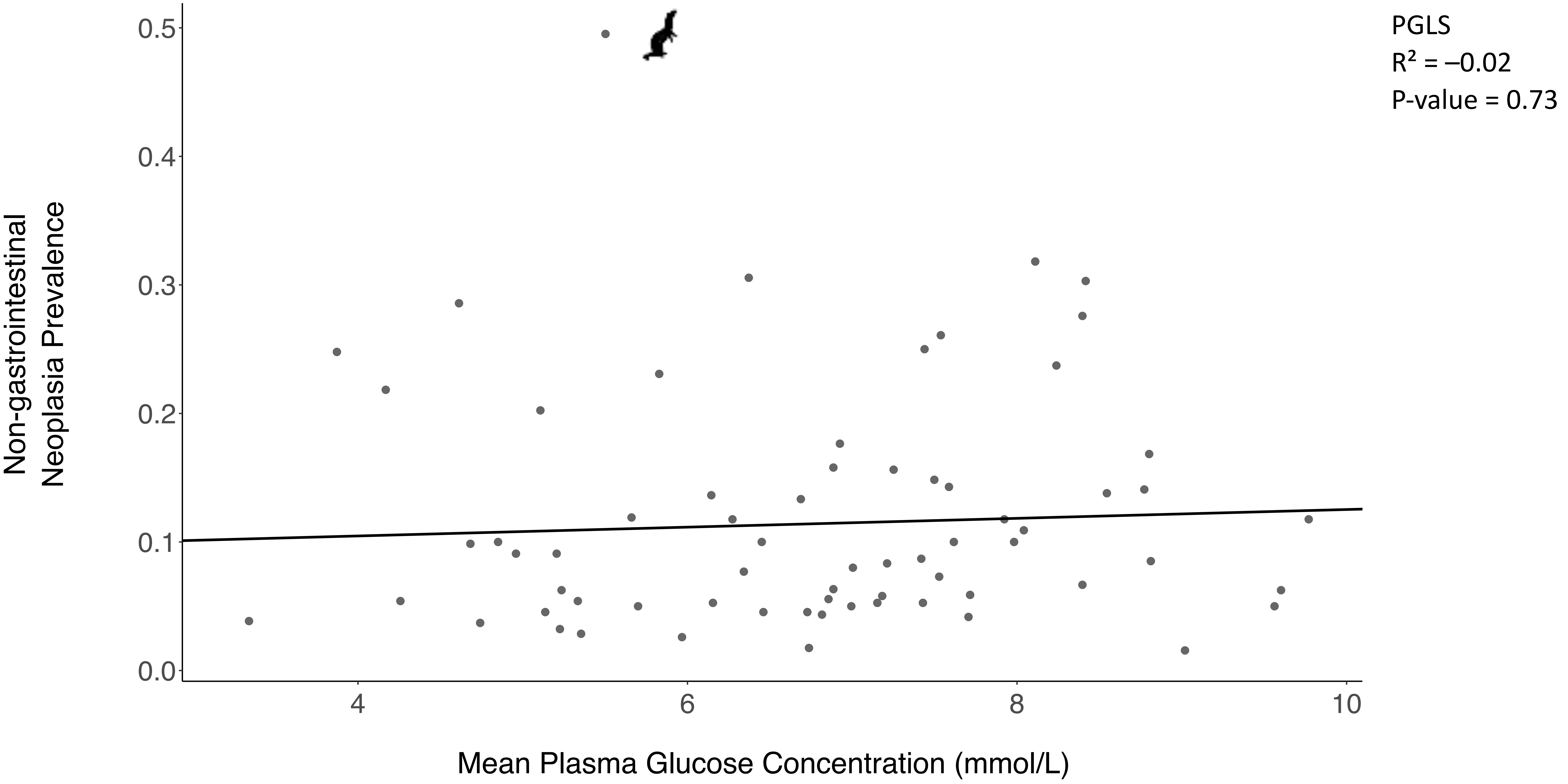

### Supplementary Figure 6

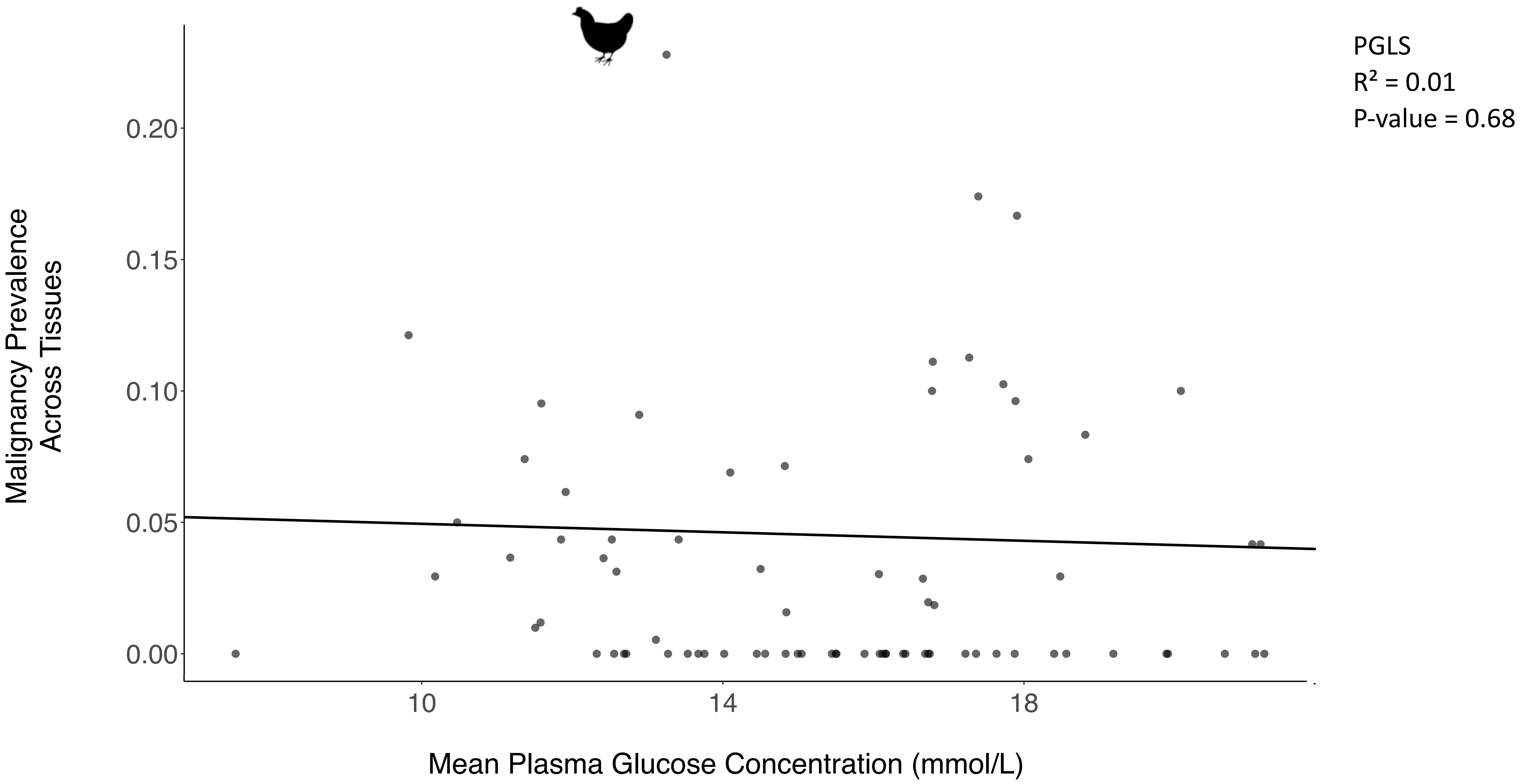

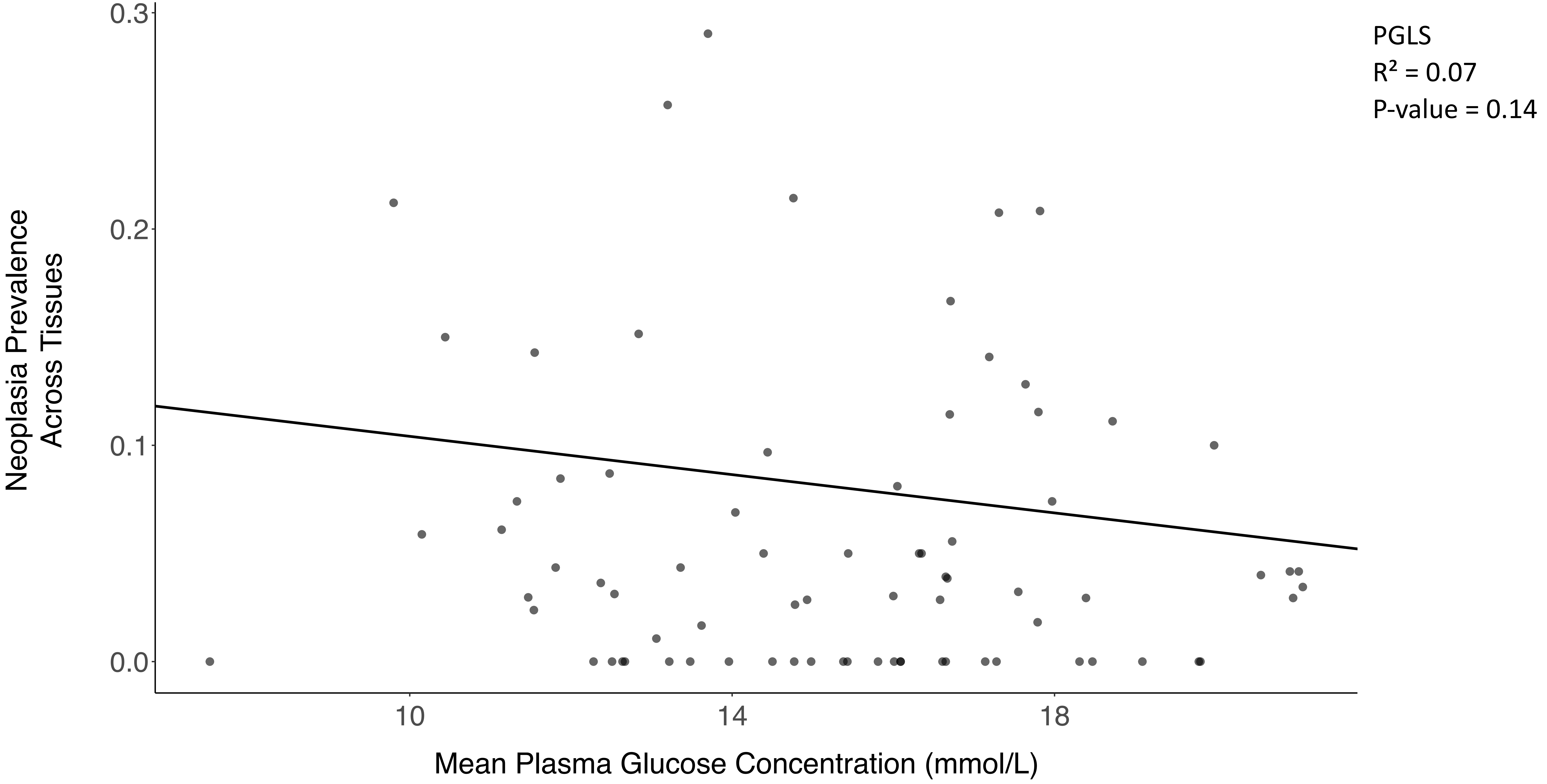

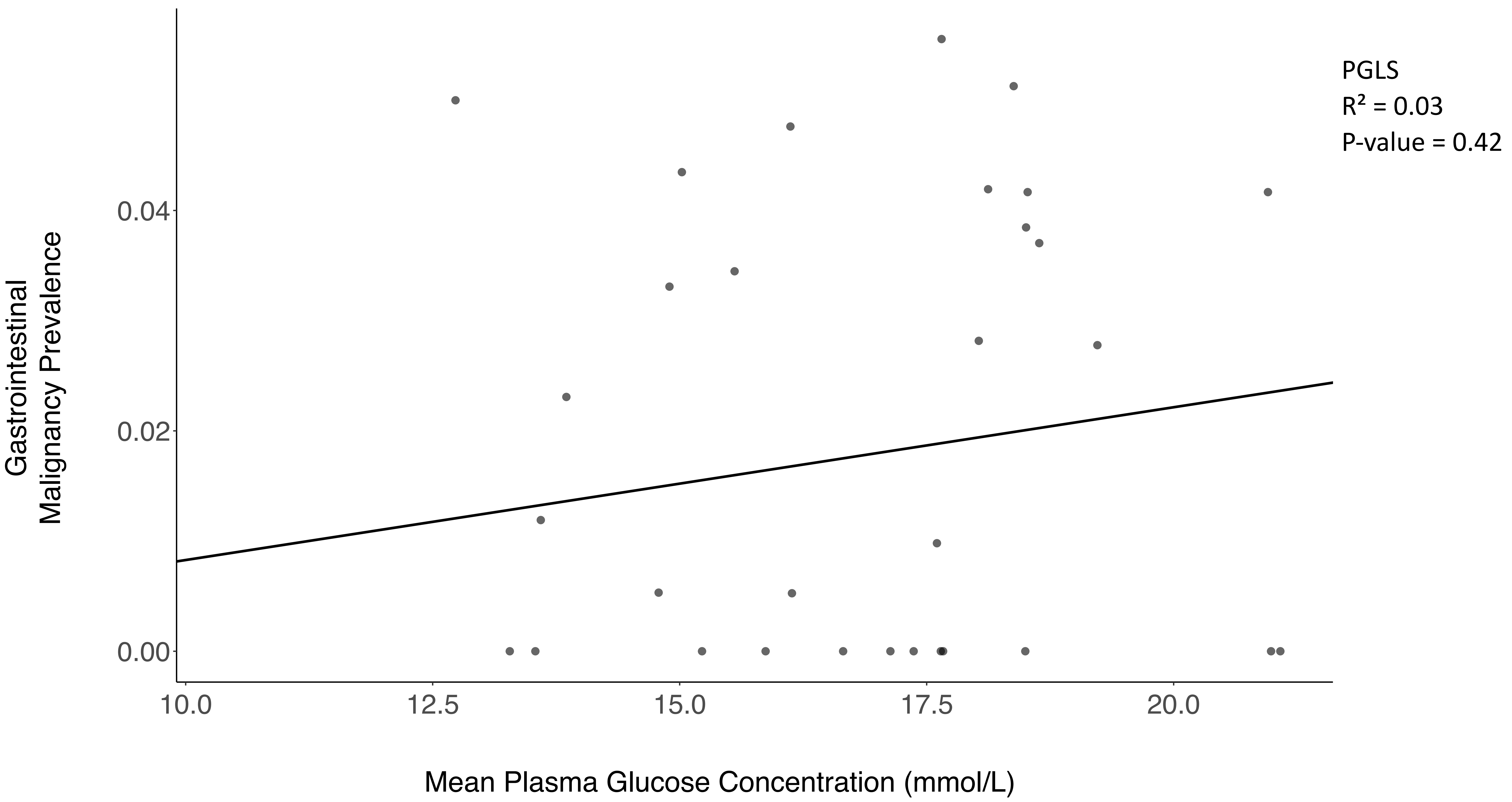

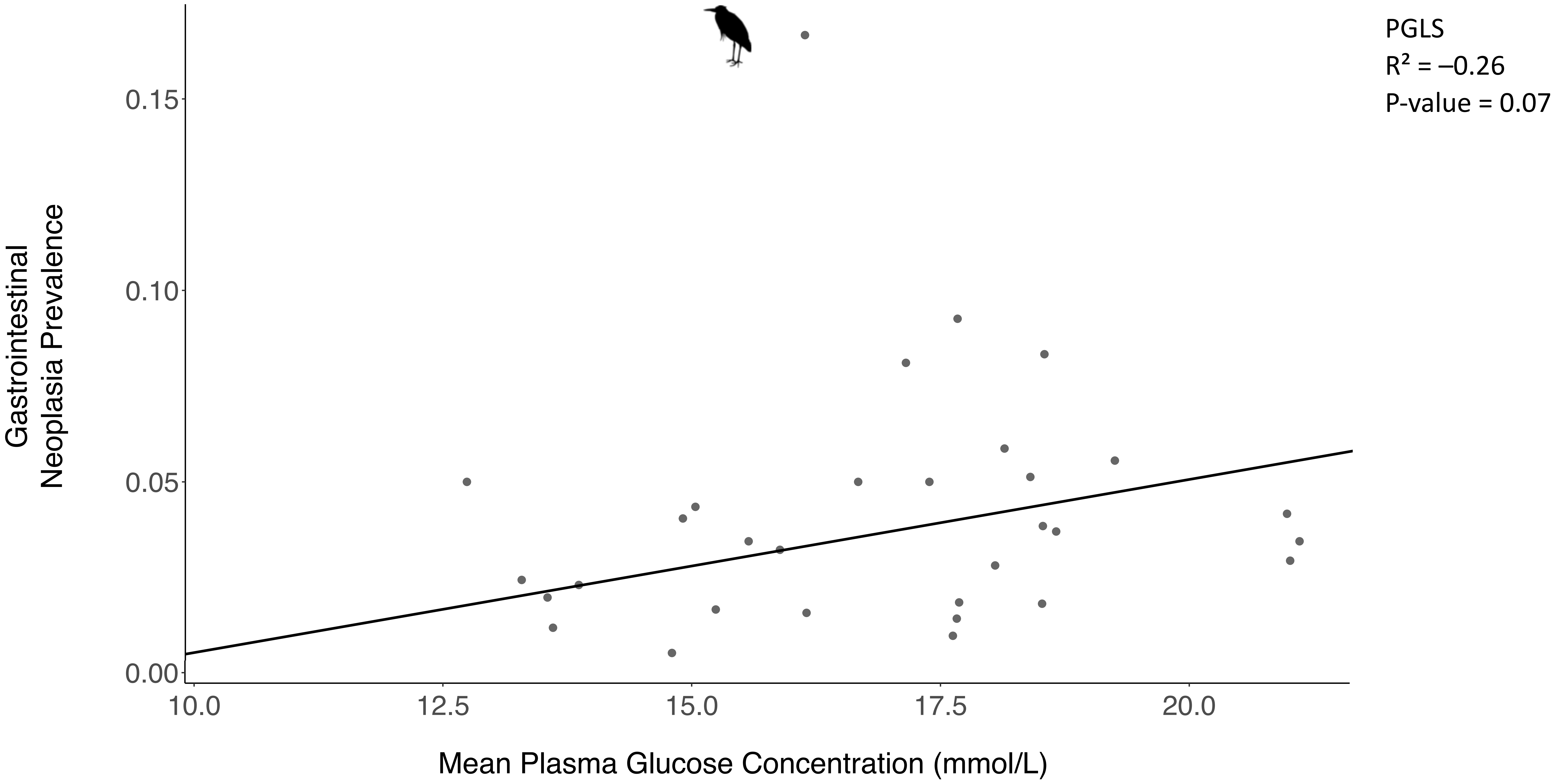

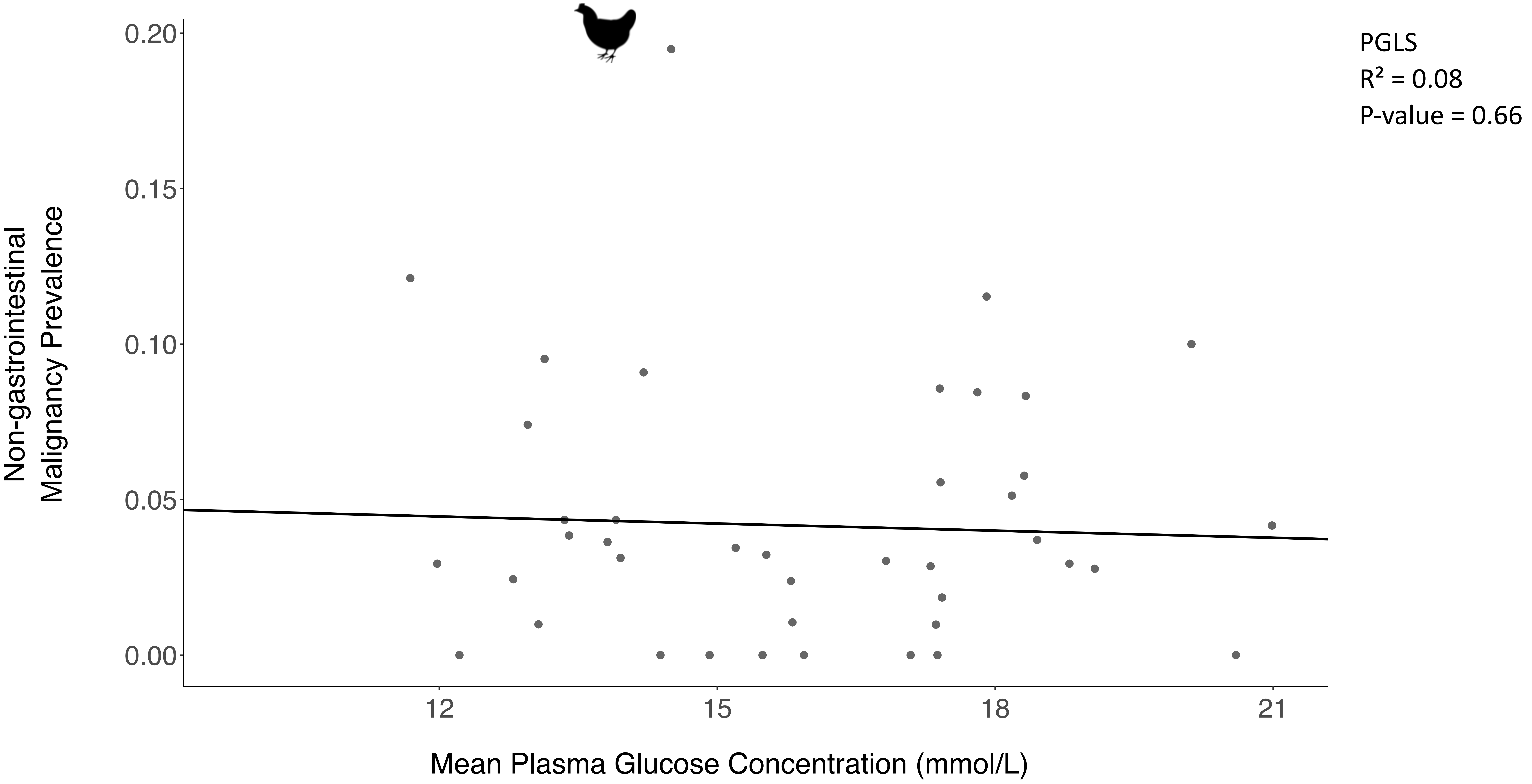

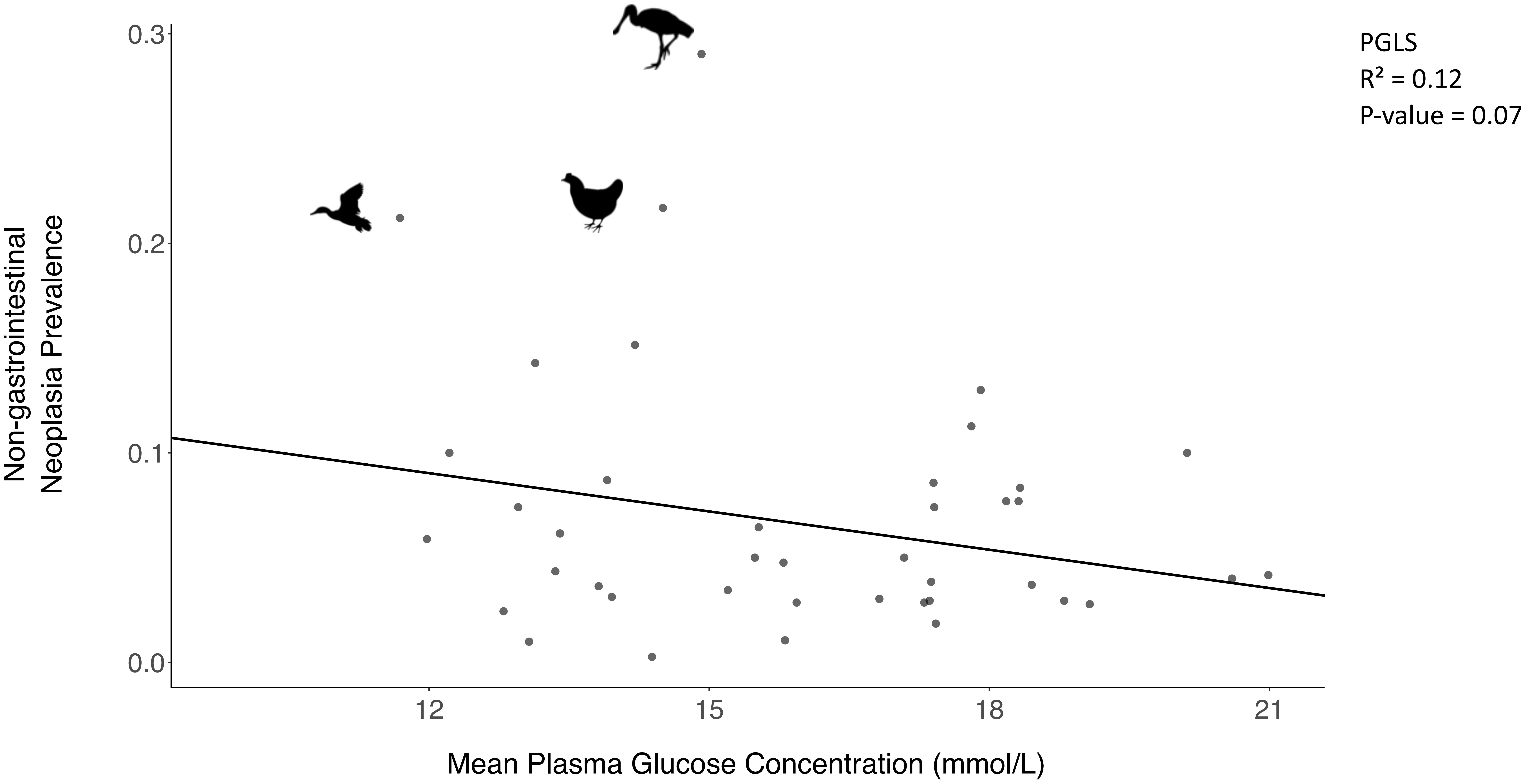

### Supplementary Figure 7

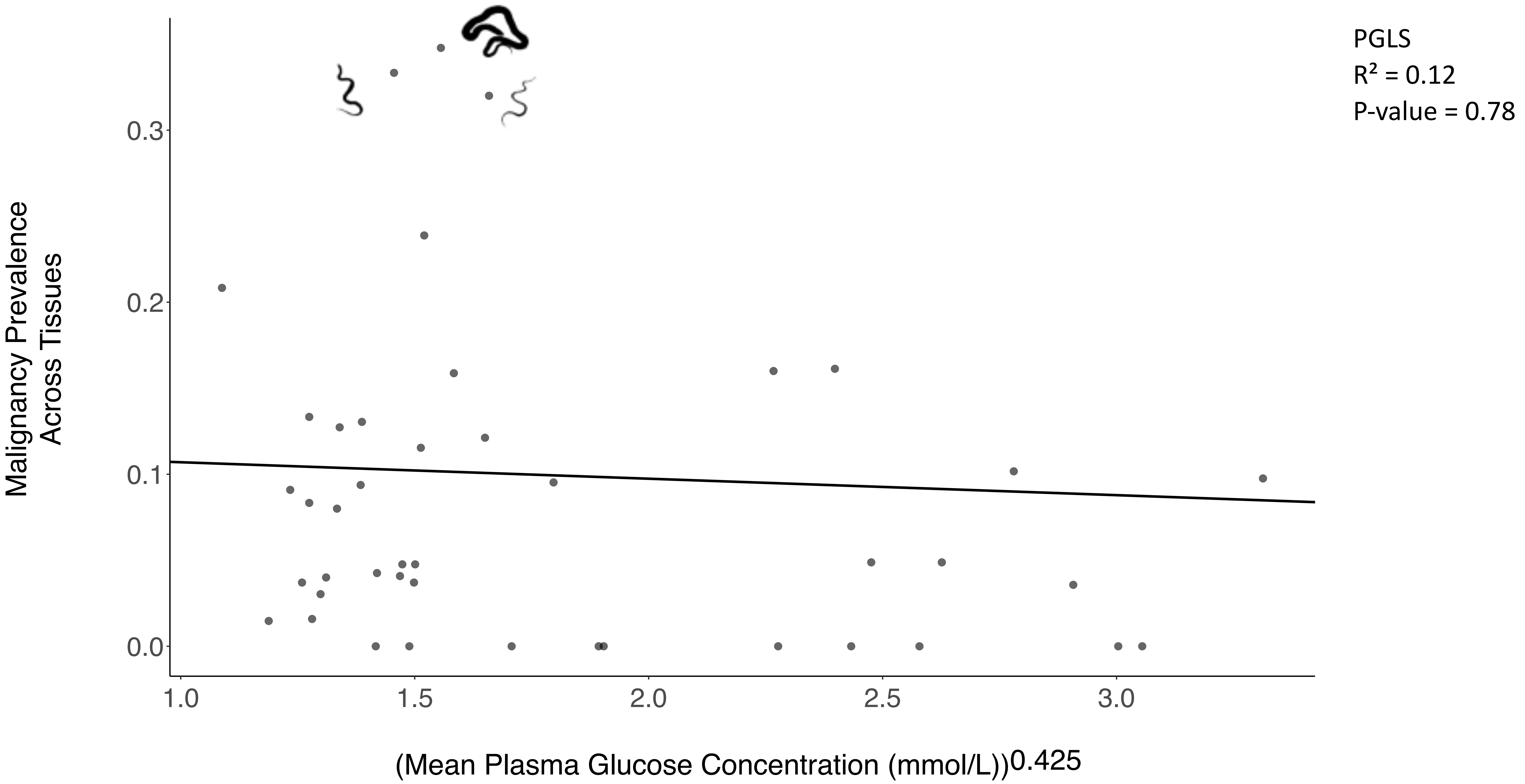

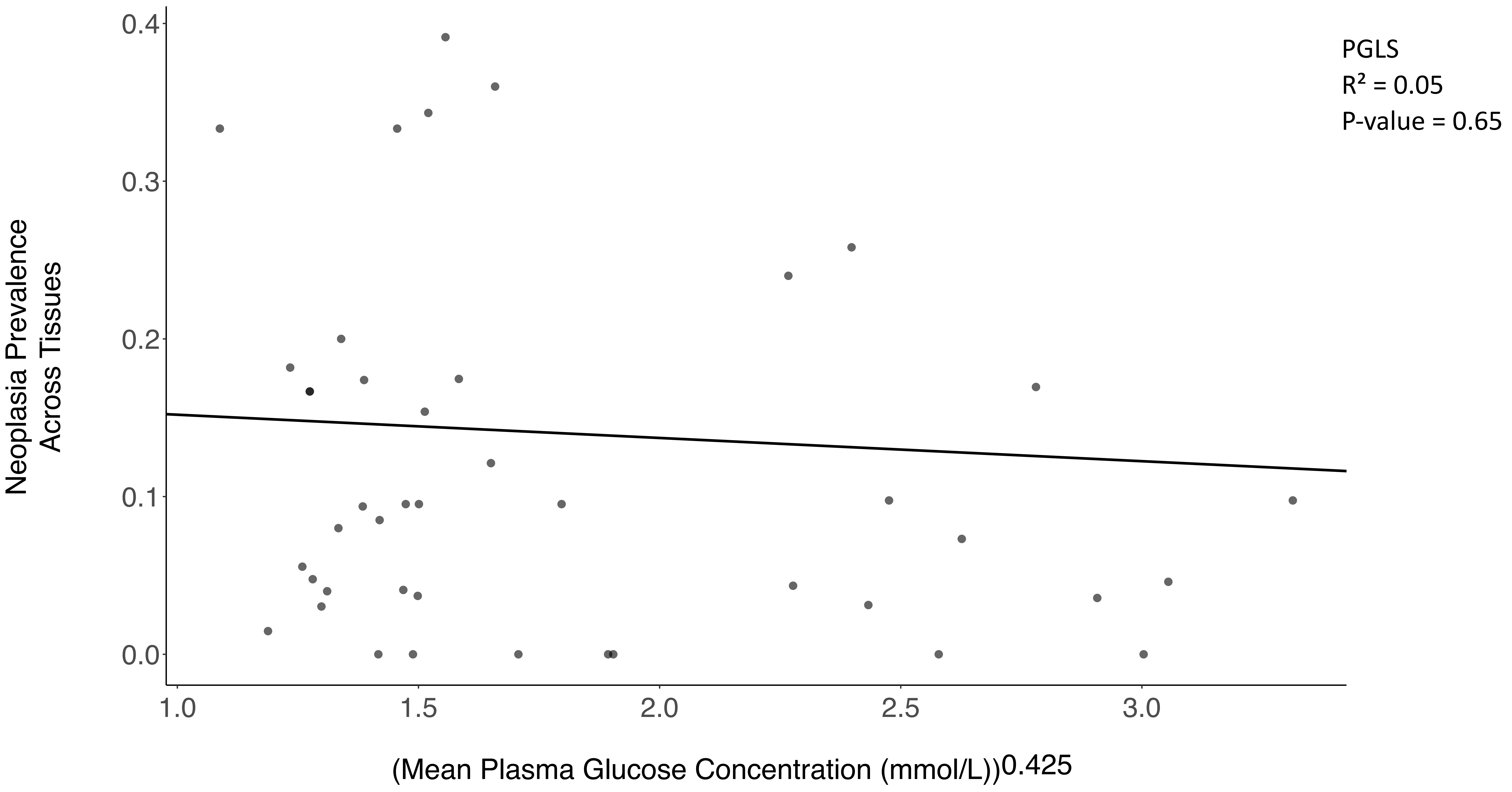

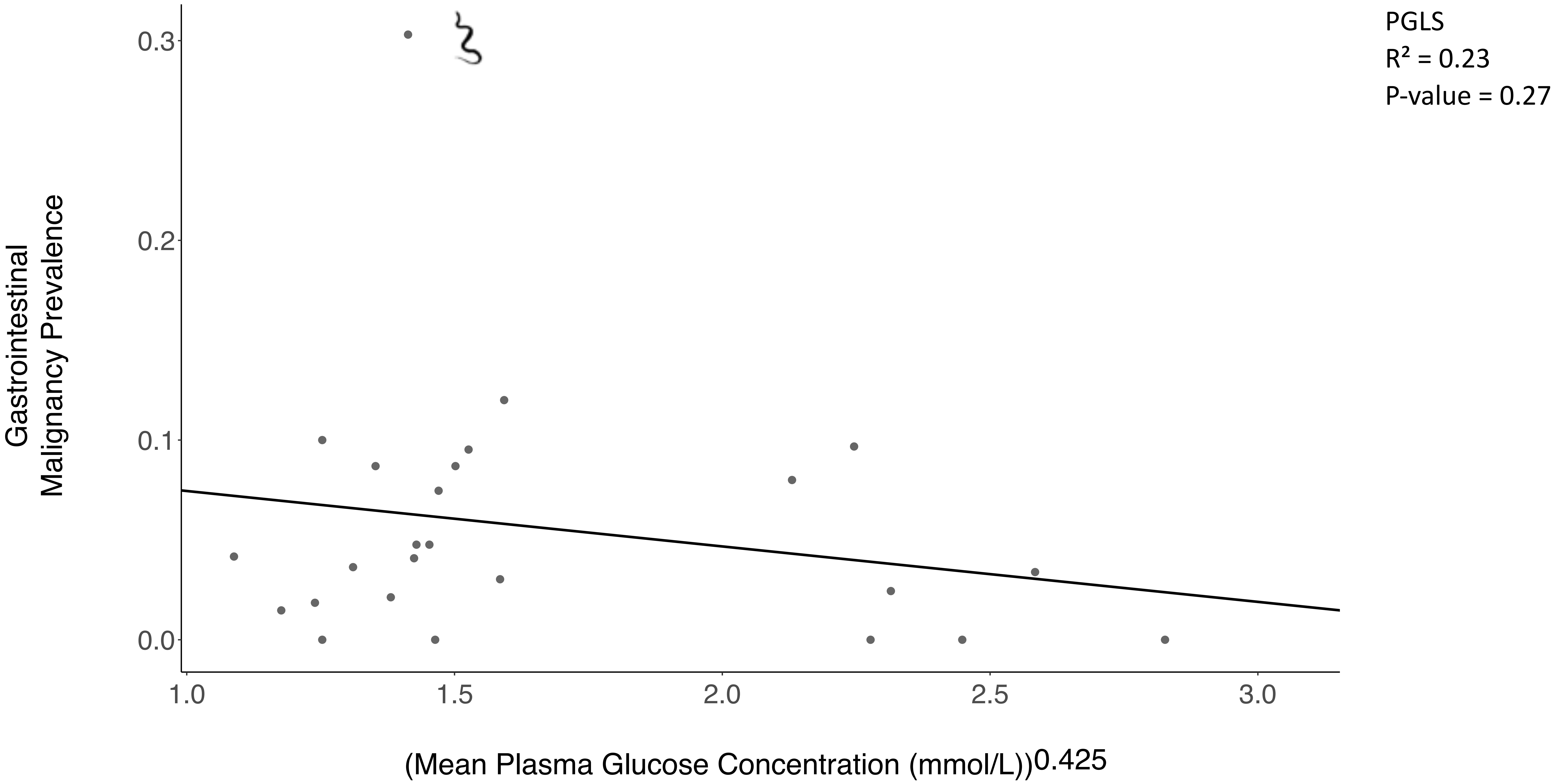

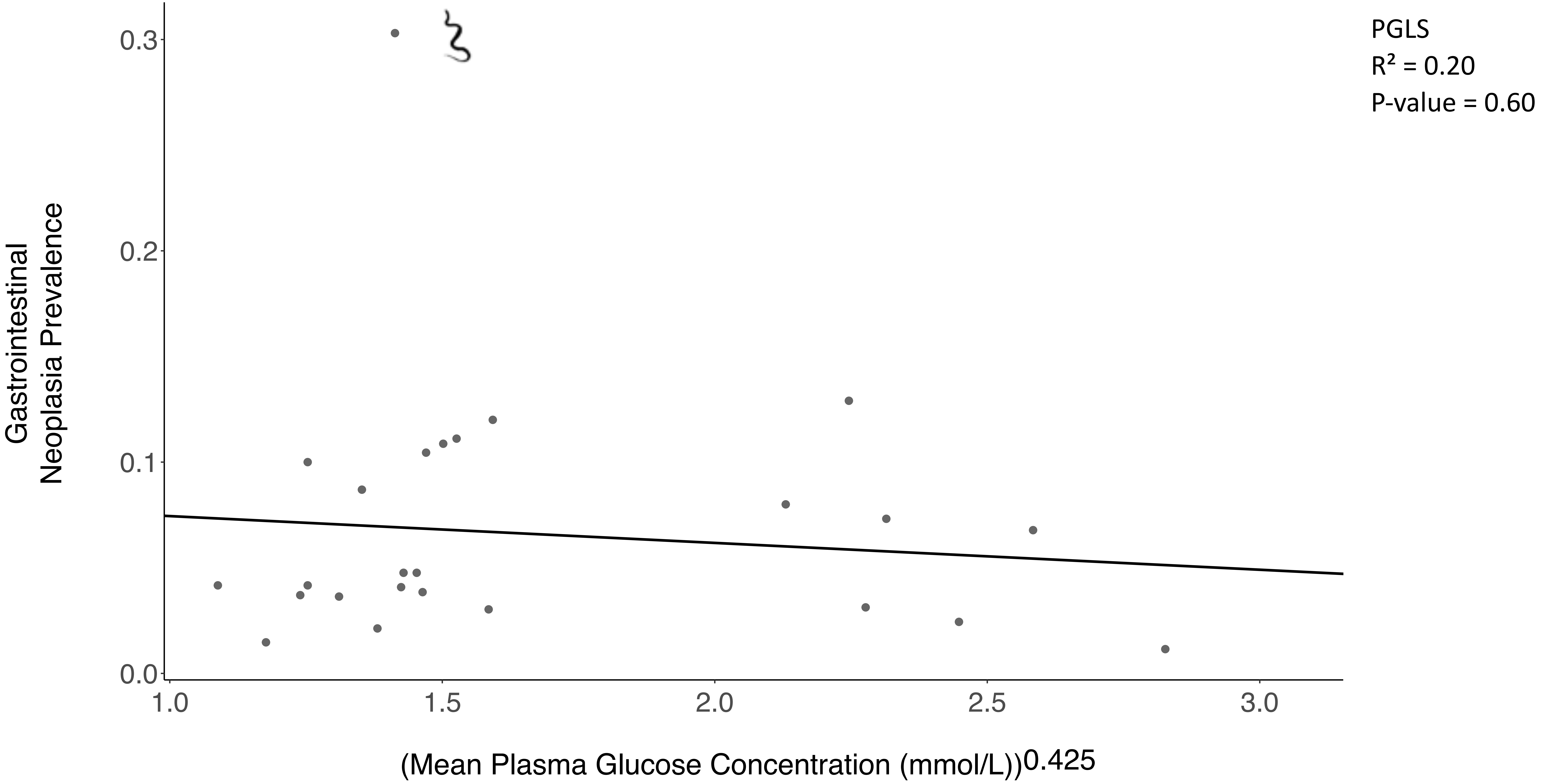

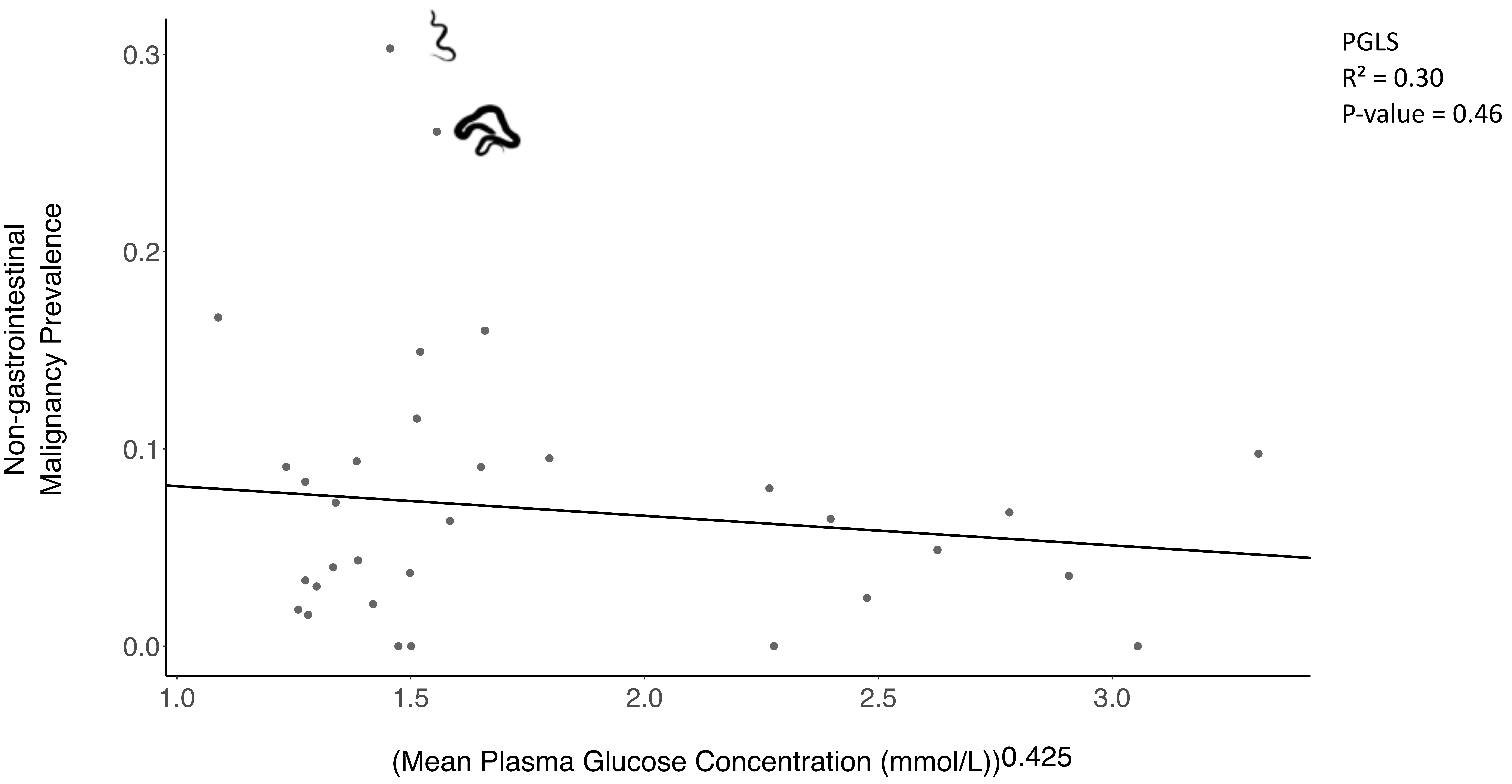

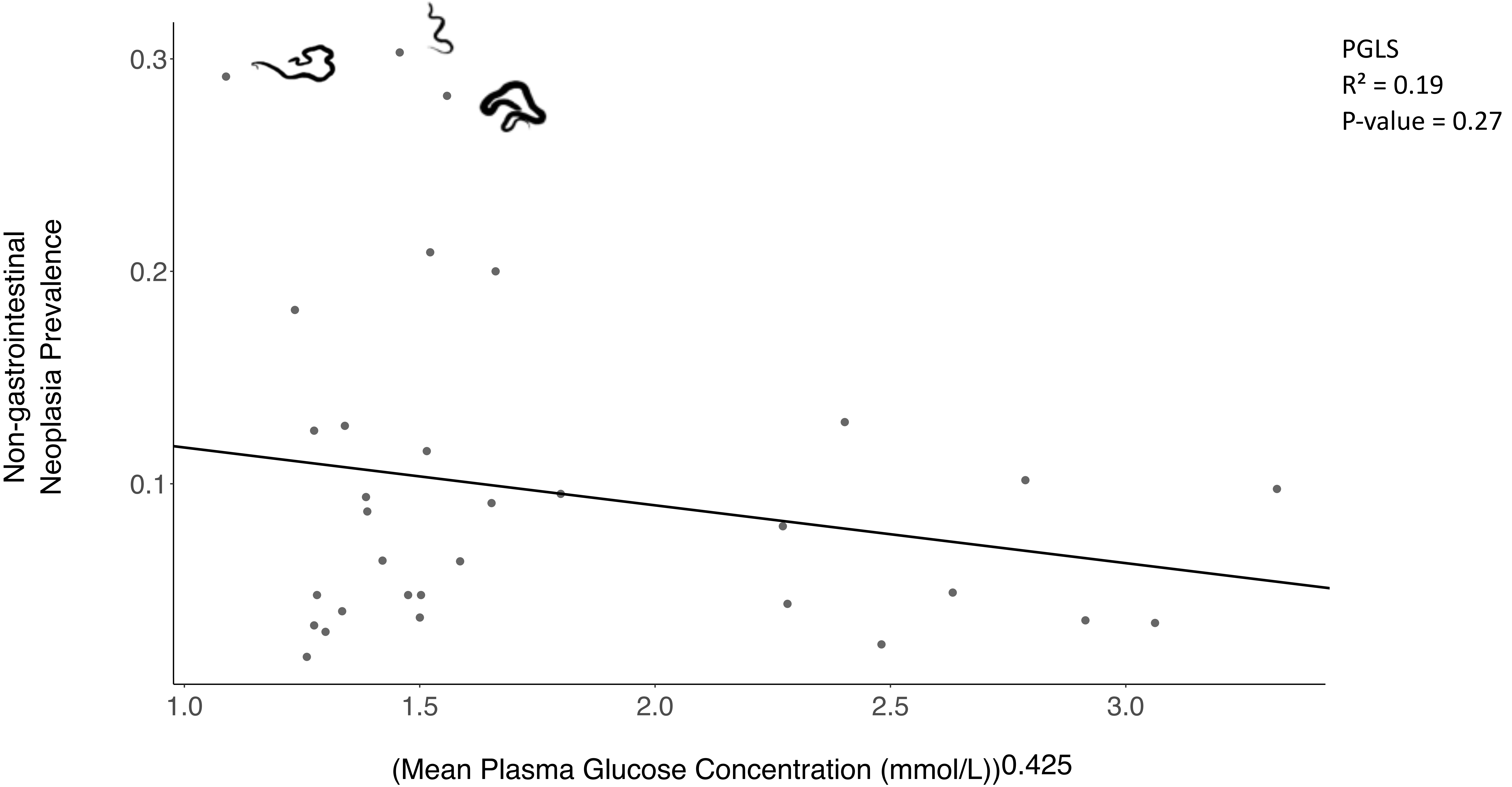
