## Supplementary Figure 2 for "The relationship between diet, plasma glucose, and cancer prevalence across vertebrates"

A

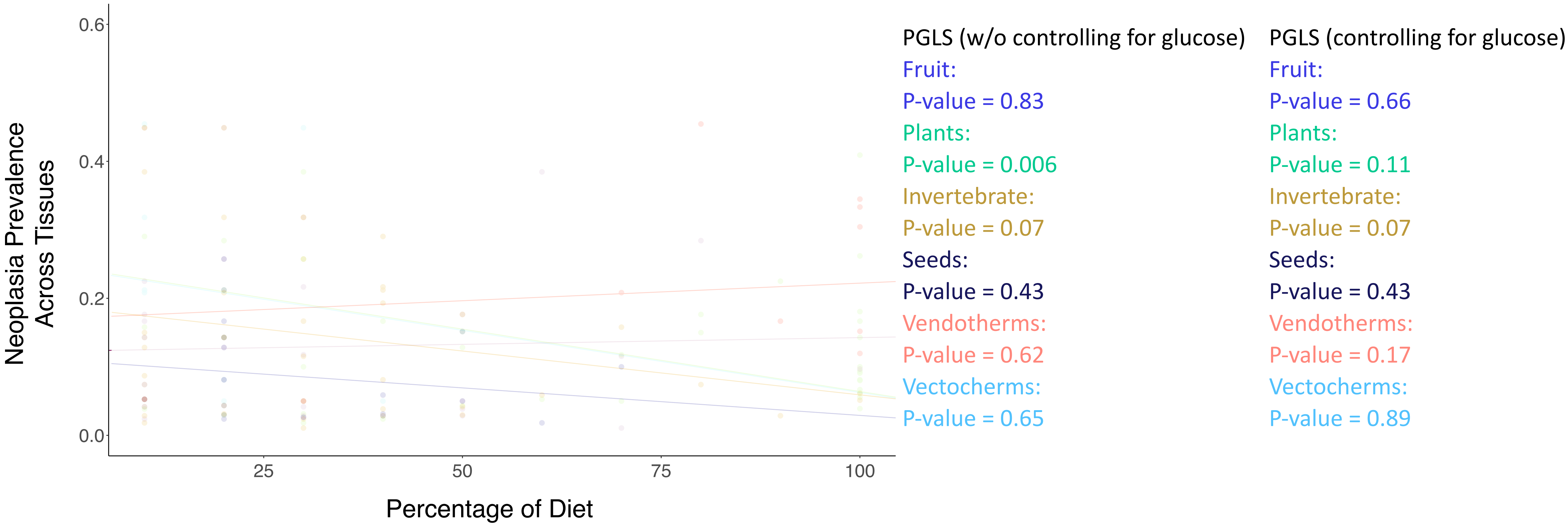

Neoplasia Prevalence Across Tissues
0.6
0.4
0.2
0.0
25
50
75
100
Percentage of Diet
PGLS (w/o controlling for glucose)
Fruit:
P-value = 0.83
Plants:
P-value = 0.006
Invertebrate:
P-value = 0.07
Seeds:
P-value = 0.43
Vendotherms:
P-value = 0.62
Vectocherms:
P-value = 0.65
PGLS (controlling for glucose)
Fruit:
P-value = 0.66
Plants:
P-value = 0.11
Invertebrate:
P-value = 0.07
Seeds:
P-value = 0.43
Vendotherms:
P-value = 0.17
Vectocherms:
P-value = 0.89

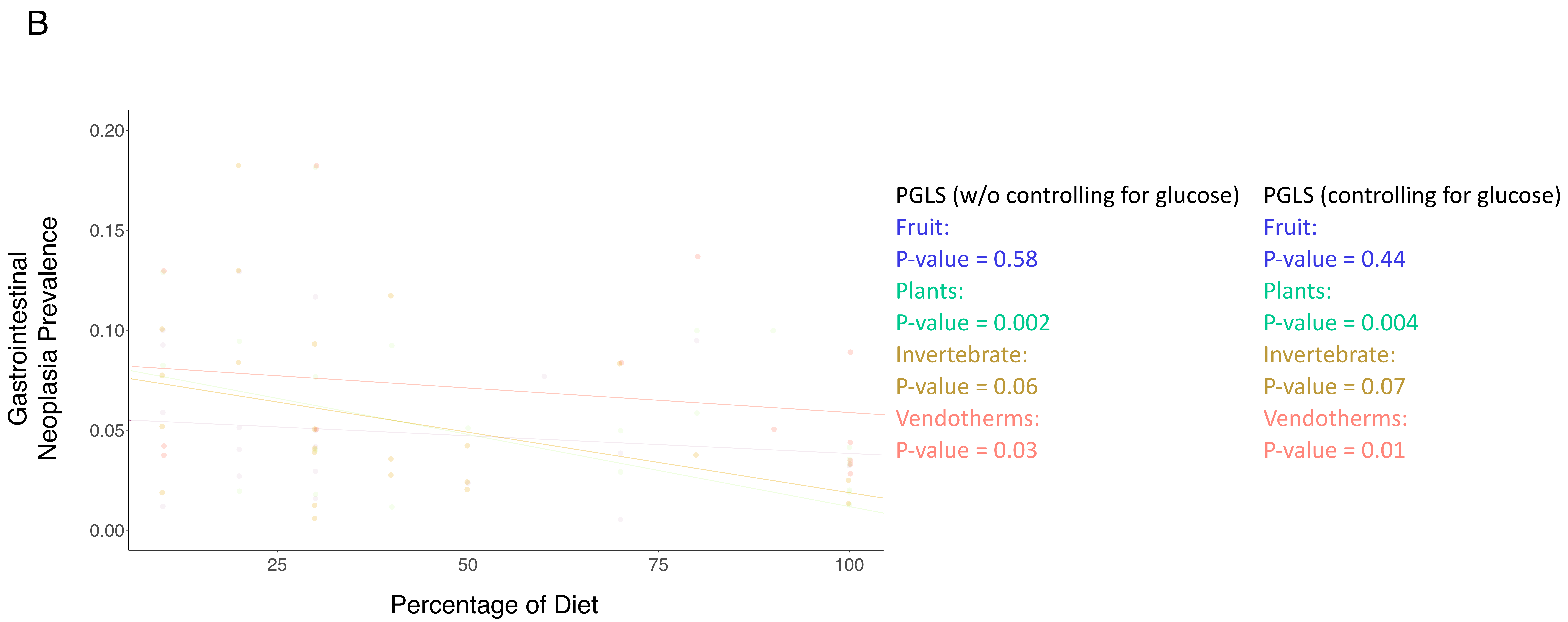
B
Gastrointestinal
Neoplasia Prevalence
0.20
0.15
0.10
0.05
0.00
25
50
75
100
Percentage of Diet
PGLS (w/o controlling for glucose)
Fruit:
P-value = 0.58
Plants:
P-value = 0.002
Invertebrate:
P-value = 0.06
Vendotherms:
P-value = 0.03
PGLS (controlling for glucose)
Fruit:
P-value = 0.44
Plants:
P-value = 0.004
Invertebrate:
P-value = 0.07
Vendotherms:
P-value = 0.01

C

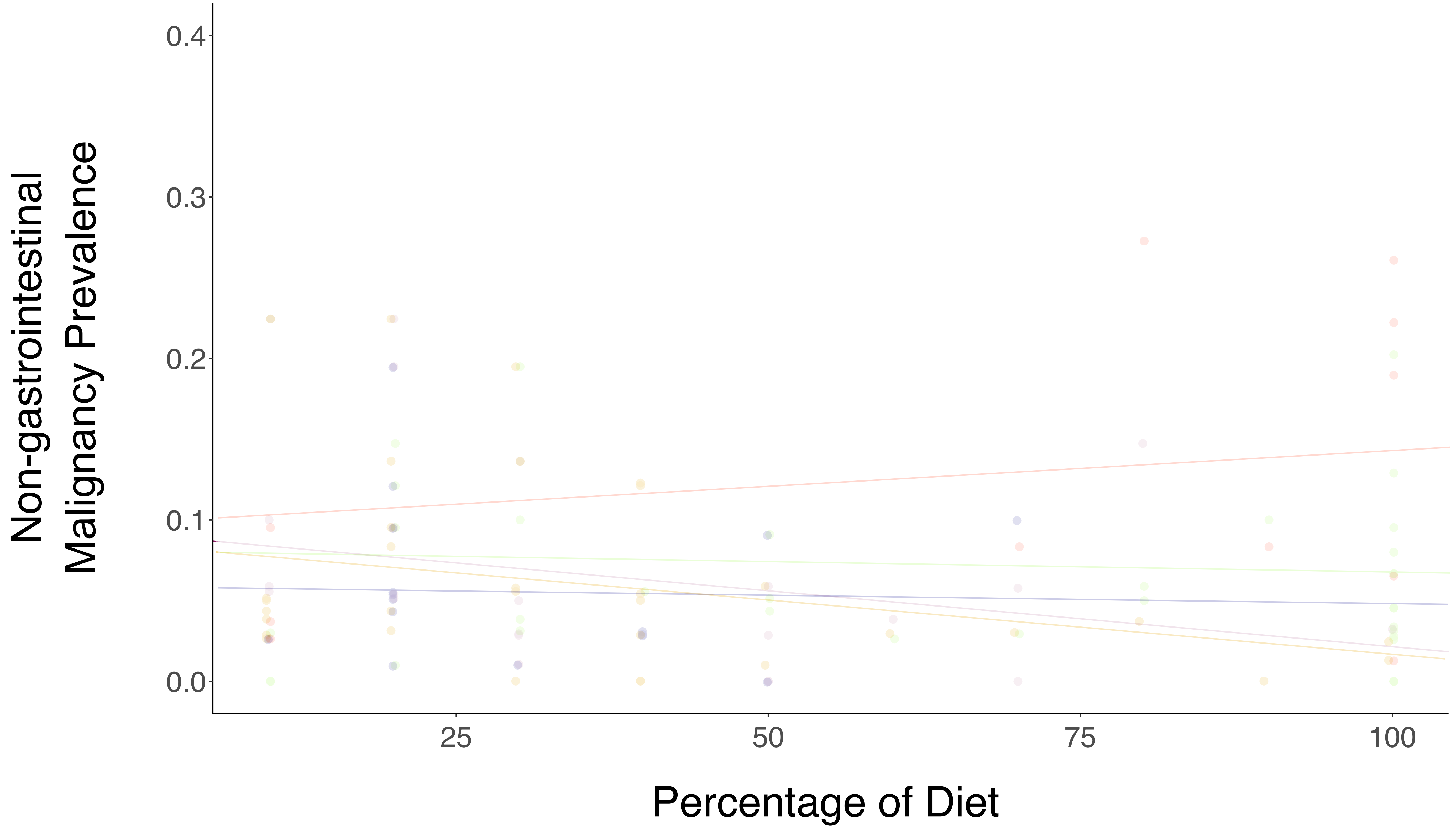


PGLS (w/o controlling for glucose)
Fruit:
P-value = 0.21
Plants:
P-value = 0.57
Invertebrate:
P-value = 0.07
Seeds:
P-value = 0.90
Vendotherms:
P-value = 0.48

PGLS (controlling for glucose)
Fruit:
P-value = 0.19
Plants:
P-value = 0.56
Invertebrate:
P-value = 0.06
Seeds:
P-value = 0.90
Vendotherms:
P-value = 0.45

D

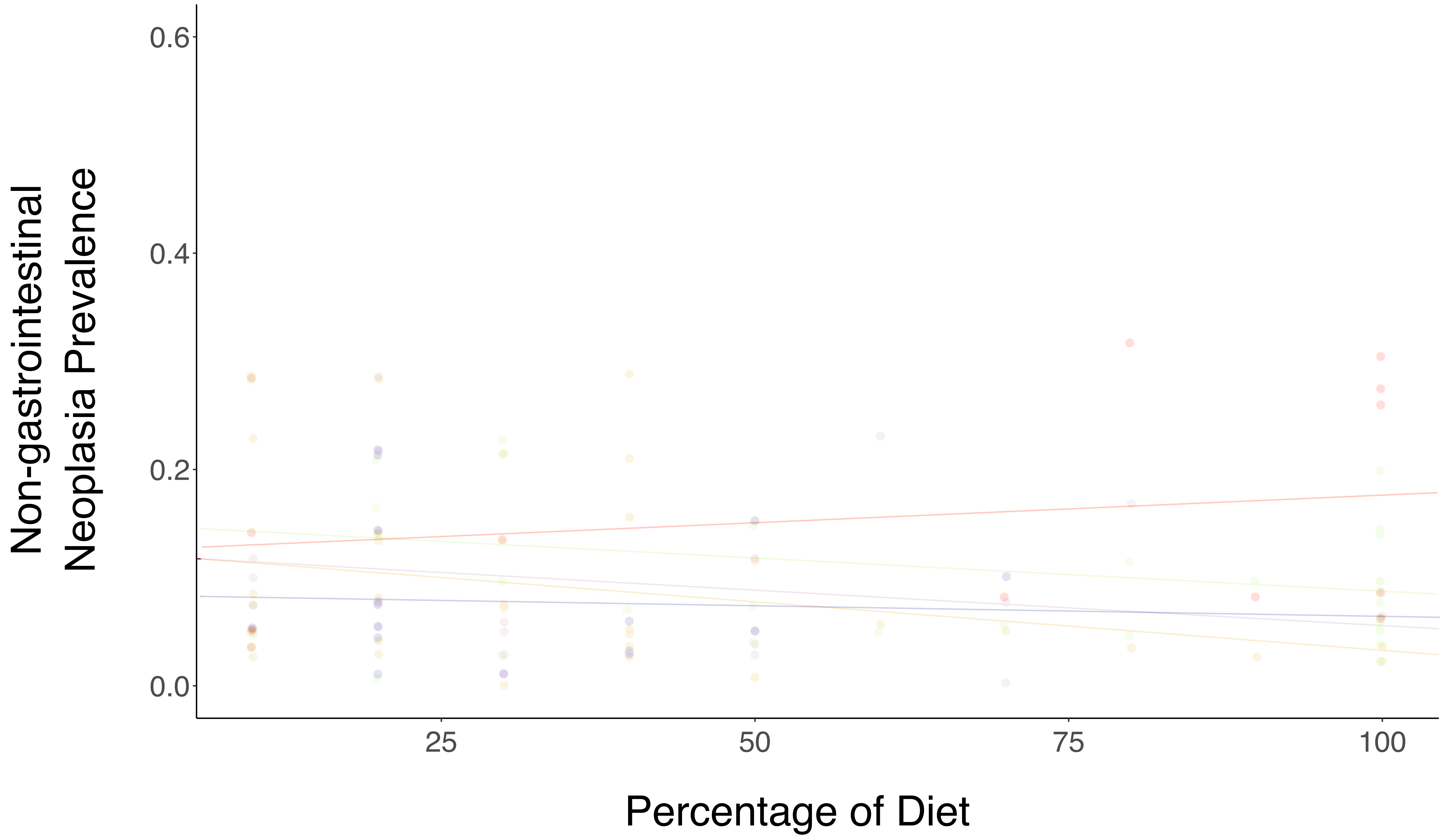


PGLS (w/o controlling for glucose)
Fruit:
P-value = 0.34
Plants:
P-value = 0.06
Invertebrate:
P-value = 0.09
Seeds:
P-value = 0.85
Vendotherms:
P-value = 0.50

PGLS (controlling for glucose)
Fruit:
P-value = 0.30
Plants:
P-value = 0.05
Invertebrate:
P-value = 0.08
Seeds:
P-value = 0.86
Vendotherms:
P-value = 0.44
